# Gene regulatory network inference and analysis of multidrug-resistant *Pseudomonas aeruginosa*

**DOI:** 10.1101/610493

**Authors:** Fernando Medeiros Filho, Ana Paula Barbosa do Nascimento, Marcelo Trindade dos Santos, Ana Paula D’Alincourt Carvalho-Assef, Fabricio Alves Barbosa da Silva

**Affiliations:** Programa de Computação Científica, Fundação Oswaldo Cruz, Rio de Janeiro, RJ, Brasil, 21040-900; Laboratório Nacional de Computação Científica, Petrópolis, RJ, Brasil; Laboratório de Pesquisa em Infecção Hospitalar, Fundação Oswaldo Cruz, Rio de Janeiro, RJ, Brasil

**Keywords:** Pseudomonas aeruginosa, gene regulatory network, multidrug resistance

## Abstract

**Background:** Healthcare-associated infections caused by bacteria such as *Pseudomonas aeruginosa* are a major public health problem worldwide. Gene regulatory networks computationally represent interactions among regulatory genes and their targets, an important approach to understand bacterial behavior and to provide novel ways of overcoming scientific challenges, including the identification of potential therapeutic targets and the development of new drugs.

**Objectives:** Our goal in this manuscript is to present a reconstruction of multidrug-resistant *P. aeruginosa* gene regulatory network and to analyze its topological properties.

**Methods:** The methodology was based on gene orthology inference by the reciprocal best hit method. We used the genome of *P. aeruginosa* CCBH4851 as the basis of the reconstruction process. This multidrug-resistant strain is representative of an endemic outbreak in Brazilian territory belonging to ST277.

**Findings:** As the main finding, we obtained a network with a larger number of regulatory genes, target genes and interactions compared to previous work. Topological analysis results are accordant to the complex network representation of biological processes.

**Main conclusions:** The network properties are consistent with *P. aeruginosa* biological features. To the best of our knowledge, the *P. aeruginosa* gene regulatory network presented here is the most complete version available to date.

## Introduction

Healthcare-associated infections (HAI) are one of the major public health problems worldwide, increasing the morbidity and mortality rates of hospitalized individuals. HAI infections are often caused by multidrug-resistant (MDR) bacteria such as *Pseudomonas aeruginosa*, especially in immunocompromised patients. In Brazil, *P. aeruginosa* was ranked as the fifth most common causative agent of HAI in patients hospitalized in adult and pediatric intensive care units, and nearly 35% of the reported strains are resistant to carbapenems, a class of antibiotics widely used in *P. aeruginosa* infections therapy^(1)^. In fact, individuals infected with MDR *P. aeruginosa* clones have a higher mortality rate (44.6%) compared to those with non-MDR infection^(2)^.

*P. aeruginosa* is a versatile pathogen that cause several types of infections affecting the lower respiratory tract, skin, urinary tract, eyes, leading to bacteremia, endocarditis, and other complications. *P. aeruginosa* infections are difficult to treat as the therapeutic choices has becoming ever more limited. Biofilm formation and the presence of intrinsic resistance-associated genes are examples of the *P. aeruginosa* arsenal against chemotherapy. In addition, this bacterium can become multidrug resistant to a broad range of antibiotics through the acquisition of new resistance mechanisms by horizontal gene transfer^(3–5)^.

In 2000, the genome sequence of *P. aeruginosa* PAO1 strain was published, providing data concerning its genome sequence, genetic complexity and ecological versatility^(6)^.The PAO1 strain is sensitive to most clinically used antimicrobial agents and has been extensively studied ever since.

In 2003, the first clinical isolate of an MDR P. aeruginosa carrying the carbapenemase gene named *bla*_SPM-1_ was identified in Brazilian territory. The SPM-1 protein is a metallo-*β*-lactamase that confers resistance to almost all classes of beta-lactams^(7)^. Most of SPM-producing isolates belong to clone ST277, as indicated through multilocus sequence typing (MLST). This clone has been associated with hospital outbreaks in several Brazilian states, and have already been found in hospital sewage and rivers^(8–10)^.

Over the past years, researchers have applied mathematical methods in order to generate computational models used to study several organisms’ behavior, contributing to the development of new products, improvement and acceleration of existing health policies, and research of novel ways of overcoming scientific challenges. This approach is often based on the construction of biological networks and pathway analysis comprising gene regulatory, metabolic, signal transduction and/or protein-protein interactions^(11)^.

A gene regulatory network (GRN) is a collection of transcription factors that interact with each other and with other molecules in the cell to regulate the levels of mRNA and protein expression. In 2011, Galán-Vásquez *et al*. ^(12)^ published the first *P. aeruginosa* GRN, analyzing its main topological properties and interactions between its regulatory components.

In this work, a reconstruction of the *P. aeruginosa* GRN of an MDR strain is described, including as much curated biological data as available to date. This reconstruction is based on the *P. aeruginosa* CCBH4851, a strain representative of an endemic outbreak in Brazilian territory caused by clones belonging to the ST277. This strain shows resistance to all antimicrobials of clinical importance except for polymyxin B, has several mechanisms of resistance and mobile genetic elements^(13)^. The implications of the choice of an MDR strain as the basis of the GRN reconstruction presented in this manuscript are discussed. In addition, GRN topological properties are analyzed, characterizing regulators, target genes, transcription factors auto-activation mechanisms, influential genes and network motifs.

## Materials and Methods

### Bacterial strains

In this manuscript, a gene regulatory network reconstruction for *P. aeruginosa* CCBH4851 is described. This strain is deposited in the Culture Collection of Hospital-Acquired Bacteria (CCBH) located at the Laboratório de Pesquisa em Infecção Hospitalar - Instituto Oswaldo Cruz/Fiocruz (WDCM947; 39 CGEN022/2010) and its genome is available in the GenBank database (accession number CP021380)^(13)^. In order to perform the orthology analysis, we used *P. aeruginosa* PAO1 ^(6)^, *P. aeruginosa* PA7 ^(14)^ and *P. aeruginosa* UCBPP-PA14 (PA14)^(15)^ as reference strains.

### Orthology-based model generation

Fitch^(16)^ defines orthologs as genes diverging after a speciation event, sharing a common ancestor. The most common approach to find orthologs is the reciprocal best hits (RBH) method^(17)^. The regulatory interaction between a transcription factor (TF) and a target gene (TG) belonging to *P. aeruginosa* PAO1, *P. aeruginosa* PA14 and *P. aeruginosa* PA7 strains were propagated to *P. aeruginosa* CCBH4851 reconstructed network if both TF and TG form RBHs. The criteria to define an orthology relationship is the existence of RBHs between the two genomes. Two genes x and x’ of the genomes X and X’, respectively, are considered orthologs if they are also RBHs, *i.e.*, if aligning the sequence of x against the gene list of X’ we obtain x’ as the best alignment, and if aligning the sequence of x’ against the gene list of X we obtain x as the best hit. Once the complete set of genomes RBHs between X and X’ is obtained, a regulatory interaction between a TF (the gene x) and a TG (the gene y) was propagated from the reference network to CCBH4851, if both TF and TG have their respective RBHs in the CCBH4851 genome. The propagation of a regulatory interaction x-y from the reference genome X holds if a pair x’-y’ exists in the genome X’ such that both (x, x’) and (y, y’) are RBH pairs. One disadvantage of RBH method is the incapacity to detect multi-to-multi orthologous relationships. In this case, RBH only picks the hit with the best score alignment, resulting in false negatives. In order to solve these false negatives, when a gene presented no orthologous in genome X’, manual curation was performed as follows: the protein sequence encoded by gene x of the genome X was searched against the genome X’ using the BLASTX algorithm. If the search returned two or more hits, the neighborhood of each hit was assessed to determine which gene in the X’ genome was orthologous to that specific protein, matching its genomic context. If the search returned no hits, the gene had no ortholog in the genome X’. This test for the propagation of regulatory interactions was performed with all interactions known in PAO1, PA7 and PA14. The all-against-all alignments were performed by the BLASTP program using stringent parameters as follows: identity 90%, coverage 90% and E value cutoff of 1e-5. Figure 1 presents an overview of the reconstruction processes.

**Fig. 1.**
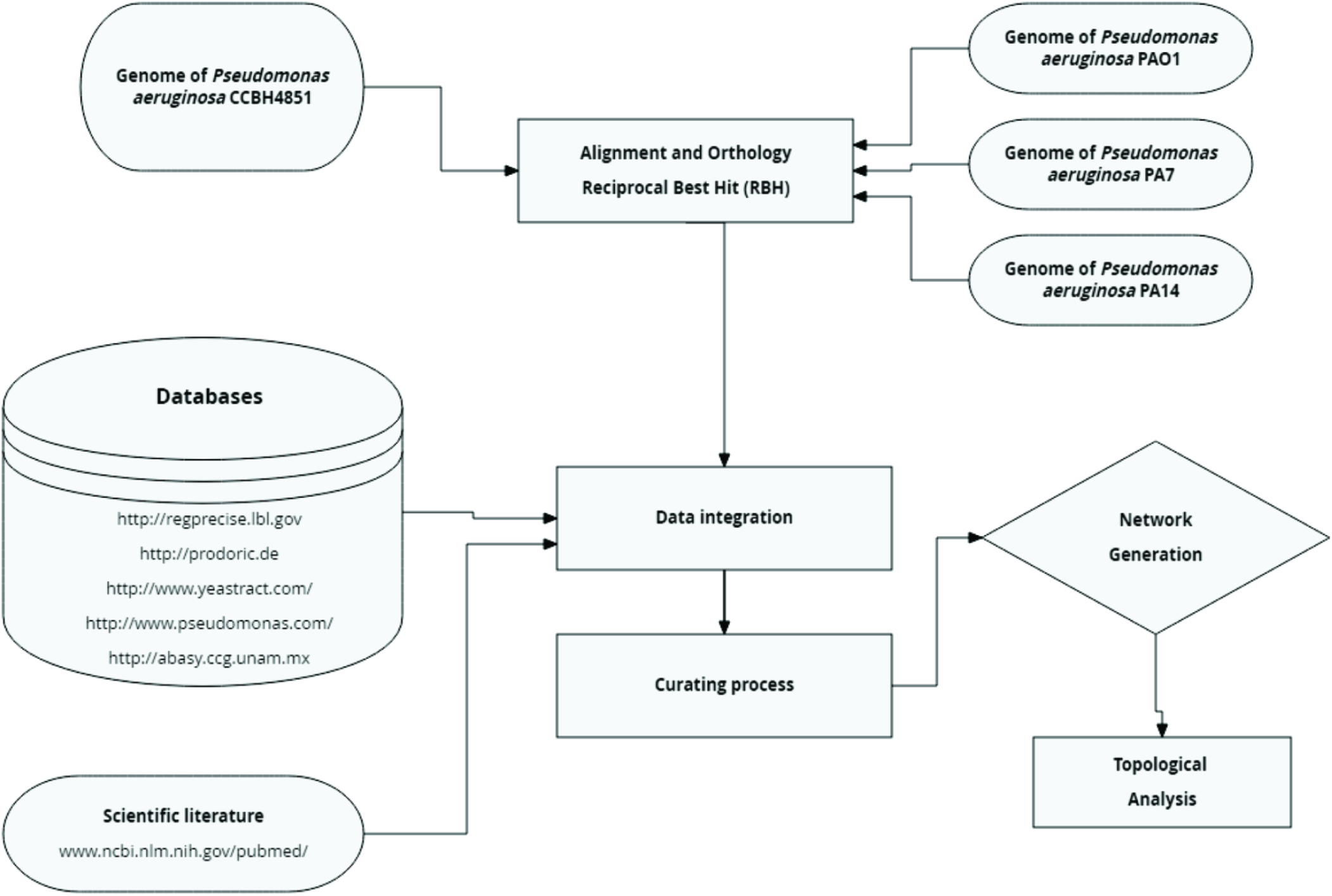
Overview of general strategy for reconstruction of the *P. aeruginosa* GRN. The process started with the alignment of *P. aeruginosa* CCBH4851 genome and the three reference strains. Next, the RBH method was applied and the resulting genes were compared against gene regulatory databases listed in the “Databases” box. Then, data obtained from these databases were integrated and submitted to the curation process, which aims to solve network inconsistencies. Finally, the GRN was generated and its topology was analyzed.

### Identification of RBHs

An algorithm was implemented using the Python programming language to automate and generate the list of RBHs in a tabular format. The last step was to identify and separate the regulators and target genes in a single table, extending the work done by Galán-Vásquez *et al*. ^(12)^.

### Data integration

The data integration process brings together biological information from all strains with the aim of organizing biological knowledge. The final network table is available as supplementary material. This table is organized into 6 columns: “Regulatory gene”, “Ortholog of the regulatory gene”, “Target gene”, “Ortholog of the target gene”, “Mode of regulation” and “Reference”. The first column lists regulatory genes of *P. aeruginosa* CCBH4851, the second column contains orthologous of regulatory genes in the reference strain (PAO1, PA7 or PA14), the third column refers to the target gene in CCBH4851, the fourth column lists orthologous of target genes in the reference strain, the fifth column describes the mode of regulation, and the sixth column indicates the corresponding reference.

### Curation process

Our group has developed a web application to support the curation of biological networks. This web application, called CurSystem^(18)^ (available from: http://pseudomonas.procc.fiocruz.br:8185/CurSystem) provides support for distributed, asynchronous interaction among specialists. Through this tool, it was possible to select specific gene interactions, discuss their main peculiarities and determine if they would be part of the network or not. This stage was fundamental to exclude doubtful biological information from the network.

### Network generation and computational analysis

The R language and Rstudio free software were used in network generation and computational analysis^(19)^. Analysis of degree, centrality, clustering coefficient, connectivity, cycles, paths and hierarchical levels were made according to previous works^(12,20)^.We used the dplyr, tibble, readr, igraph and scales packages. The package igraph was used for computation of feed-forward loop (FFL) motifs (function triad_census). The measurement of network degree-entropy was made according Breitkreutz *et al.* ^(21)^.

All data and code are available as supplementary files.

## Results

### General features of the gene regulatory network

The *P. aeruginosa* network reconstruction resulted in a total of 1046 genes, of which 42 behave as regulatory genes, 96 both as regulatory and target genes (*i.e.*, a TF is influenced by another TF in the network), and 908 target genes. We found 1576 regulatory interactions between regulators and their target genes. Altogether, the genes represent approximately 16.52% of the *P. aeruginosa* CCBH4851 genome used as the model organism in this work. Despite the apparent small coverage, we have included most transcription factors with described function among the 138 regulators in the *P. aeruginosa* CCBH4851 network. The number of regulatory genes, target genes and interactions represent an increase of 44.9%, 34.69% and 35.27% compared to previous work, respectively^(12)^. Network enrichment was not the only aspect observed in the *P. aeruginosa* CCBH4851 gene regulatory network reconstruction. As the reconstruction was based on the RBH method, comparing the CCBH4851 genome annotation with reference strains, it was not possible to infer an orthology relationship for some genes, particularly *oprD* and *mexZ*, which are genes involved in antibiotic resistance mechanisms. The curation process revealed that these genes were either fully absent or annotated as pseudogenes in CCBH4851. A pseudogene is a DNA sequence that resembles a gene from the reference genome, but has suffered modifications such as point mutations, insertions, deletions, premature stop codons or frameshifts, being impossible to attest if its product is still functional in the target organism without proper experimentation. The lack of orthology resulted in the exclusion of these genes from *P. aeruginosa* CCBH4851 GRN. In addition, some notations were kept as listed in the previous network^(12)^ and in databases and/or scientific literature used. For example, *ihf* (for integration host factor) represents not a single gene, but a complex composed of the product of *himA* and *himD* genes that together act as a TF of several target genes. On the other hand, regulatory systems such as quorum sensing or two component systems are often formed by a pair of genes, but only one of them is able to bind in the promoter region. However, both genes are listed as regulatory genes. This way we could maintain an equivalent notation to previous networks^(12)^.

### Basic network topological analysis: number of vertices, number of edges and density

We identified 1576 edges in the CCBH4851 network. These interactions were classified in four types: activation (“+”), repression (“-”), dual (“d”) (when the regulatory gene can act as an activator or repressor, depending on certain conditions) and unknown (“?”). Figure 2 is an illustration of the CCBH4851 GRN. Network density is a measure of interconnectivity between vertices. It is the ratio of the actual number of edges in the network by the maximum possible number of edges. The regulatory network of the CCBH4851 strain has a density (1.44e-03) which is slightly lower than the observed density for the PAO1 strain (2.12e-03) but maintains the same order of magnitude. A network diameter indicates the path length between the two most distant nodes. The CCBH4851 GRN has a diameter of 12 nodes while the previous network has a diameter of 9 nodes. Another measure, the average path distance, also called average shortest path, is the average distance between two nodes. While the previous network presented an average of 4.08, CCBH5851 GRN presented an average of 4.80 ^(12,22)^.

**Fig. 2.**
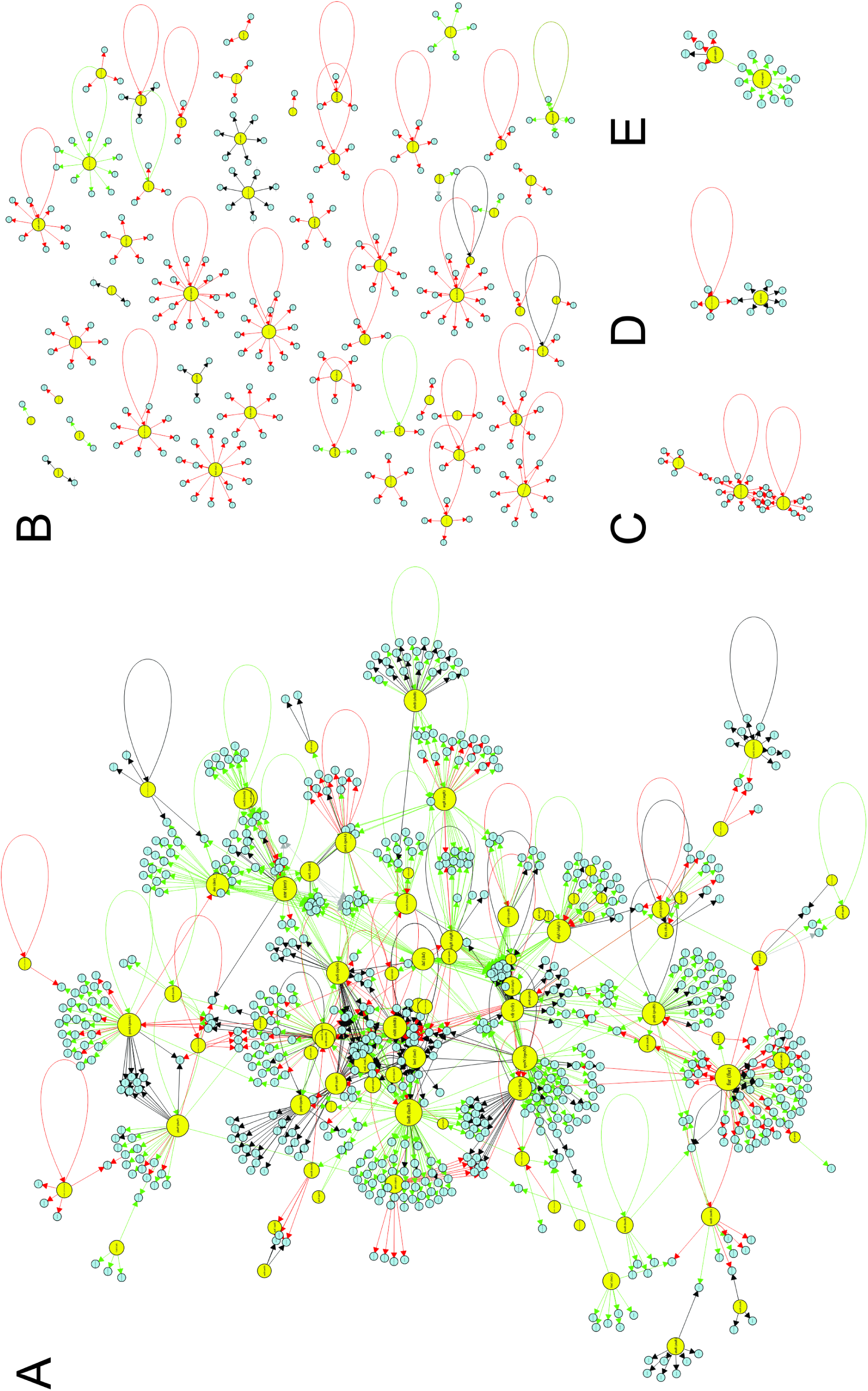
Visualization of the *P. aeruginosa* CCBH4851 gene regulatory network. The yellow circles represent regulatory genes, light blue circles represent target genes, black lines indicate an unknown mode of regulation, green lines indicate activation, red lines indicate repression and gray lines a dual mode of regulation. (A) The GRN large highly connected network component; (B) All regulatory and target genes that have no connections with the component depicted in A; (C, D, E) Clusters of lower connectivity compared to the component depicted in A. All figures are available with better resolution in the supplementary material.

The degree *k(i)* of a vertex *i* is defined as its number of edges. Edges in directed networks can be of two types: they can “depart” from or “arrive” at node *i*, defining its “incoming” (*k-in*) and “outgoing” (*k-out*) degrees respectively. It was observed for the CCBH4851 GRN that, on average, each vertex is connected to 3 other vertices, same value reported for PAO1 GRN. Figure 3 illustrates incoming (3A-B) and outgoing (3C-D) degree distributions for the CCBH4851 GRN.

**Fig. 3.**
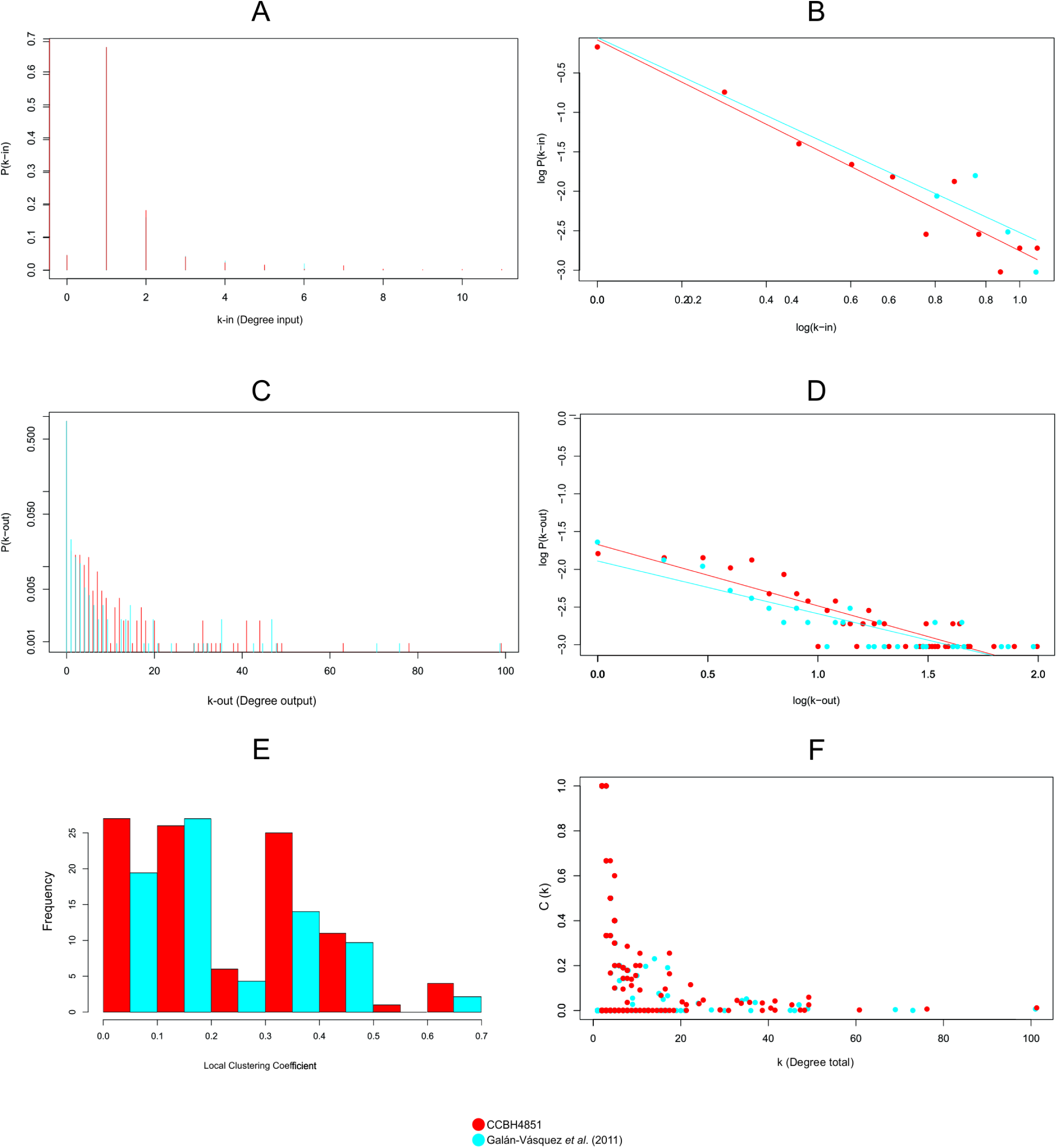
Graphic representation of topological measurements of the *P. aeruginosa* CCBH4851 gene regulatory network (red) compared to the previously published network (blue) ^(12)^. (A) and (B) Incoming degree distribution of the *P. aeruginosa* CCBH4851 GRN. (C) and (D) Outgoing distribution of the *P. aeruginosa* CCBH4851 GRN. For clarity, the distributions are plotted both on a linear (A, C) and on logarithmic scale (B, D). (E) Local clustering coefficient distribution. (F) Clustering coefficient by degree.

Scale-free is a common topology classification associated with biological networks, corresponding to complex networks which degree distribution follows a power law. In scale-free networks, most nodes (vertices) have few connections and few nodes have a large number of connections. In this way, scale-free networks are dominated by a relatively small number of high degree nodes, generally called hubs^(23)^.

The degree distribution can be approximated by:

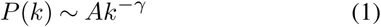

Equation 1 corresponds to a power-law distribution and the exponent *γ* is its degree exponent^(24)^. The degree distribution in figures 3B and 3D is shown on double logarithmic axis, and the straight line is consistent to a power-law distribution. For the k-in, the estimated value for *γ* was 2.89, very close to the value reported by the reference work (*γ* =2.717)^(12)^.

### Clustering coefficient distribution

Given a node *i* with *m(i)* neighbors in a directed network, the maximum number of edges connecting the elements of this neighborhood is given by *m*_max_*(i) = m(i)(m(i)-1)*. The local clustering coefficient *C(i)* is defined as the ratio between the actual number of edges *N(i)* occurring in node *i* neighborhood and *m*_max_*(i)* ^(25)^. The local clustering coefficient is defined as *C(i)=N(i)/ m*_max_*(i)*. In GRNs, the local clustering coefficient *C(i)* is interpreted as the interaction between genes forming regulatory groups. The distribution of local clustering coefficients can be seen in Figure 3E.

On the other hand, the global clustering coefficient is proportional to the number of triangles present in the network, disregarding the directionality of the edges. A tringle is a set of three nodes with at least two connections between them. We can have closed triangles, with three connections within the set, and open triangles, with only two edges. The global clustering coefficient *C* is the ratio between the number of closed triangles and the total number of triangles (closed or open) in the network. The CCBH4851 network has a global clustering coefficient equal to 3.2e-02. Another interesting feature to observe is the correlation between the local clustering coefficient *C(i)* and the degree *k(i)*, as shown by the scatter plot in Figure 3F. The observed correlation is negative, and the figure also shows that the vertices with high degree *k* correspond to the same vertices with null clustering coefficients, while the vertices that form clusters have low degrees. From this observation, it is confirmed that strongly cohesive groups are exceptions in the network and are formed by small number of genes. These results were obtained for both the CCBH4851 and the previously published *P. aeruginosa* GRN^(12)^.

### Connectivity

Network connectivity is a concept that reflects the associations between every pair of genes. Nodes were considered part of a connected component when they interacted through a direct or an indirect link (intermediate connections). In the connectivity analysis, network interactions were considered undirected. Similar to the reference GRN, the CCBH4851 network was disconnected, it presented one large connected component (including 751 nodes) and more 48 small connected components, a larger number when compared to previous work^(12)^. However, the fact that network is disconnected at specific points could have several causes: (i) natural behavior of the organism, *i.e.*, not all genes in a complex genome are linked, since cellular processes can be compartmentalized or global, constitutive or growth phase-dependent, (ii) lack of sufficient biological information to infer interactions, (iii) overall, *P. aeruginosa* genomes maintain a conserved core component which accounts to the majority of the genome; on the other hand, additional strain-specific blocks of genes are acquired by horizontal gene transfer as the result of evolutionary events which can reflect in a decreased similarity rate with reference strains, therefore an increased similarity rate with newly reported strains; this process can reflect on loss of existing interactions or gain of interactions still not fully described, thus lacking connection with other components in the network.

### Dominant activity and autoregulation

The analysis of the frequency of the different modes of regulation indicated that activation is the predominant type of regulation mode in the CCBH4851 network, with frequency values very similar to those previously observed for the *P. aeruginosa* GRN. Overall, 48.92% of the interactions are of the activation mode, 28.8% repression mode, while 22.27% is dual or unknown mode. Although the distribution pattern was maintained, a significant enrichment was observed in the negative and unknown regulation modes. When considering autoregulation, *i.e.*, a gene regulating its own expression, the CCBH5851 GRN presented a predominance of negative autoregulatory motifs, unlike the findings of Galán-Vásquez et al. ^(12)^.

### Motifs

The existence of cycles or motifs in biological networks is a necessary condition for the existence of multiple stationary states or attractors. In gene regulatory networks, the most common 3-genes motif is the feed-forward loop (FFL). The FFL motif comprises a gene A that regulates gene B. Then, both A and B regulate gene C. There are two types of FFL motifs: (i) coherent, when the regulatory effect of both paths, direct and indirect, are the same; (ii) incoherent, when the regulatory effects are different. In this work, we computed the total number of FFL motifs, the number of coherent type I FFL motifs, where all interactions are activations, and the number of incoherent type II motifs, where all interactions are repressions^(26)^. The CCBH4851 has a larger number of FFL motifs (when considering all variations), when compared to the GRN published by Galán-Vásquez *et al*. The coherent type I FFL motif was the most abundant in both networks, with 82 representatives in the PAO1 GRN and 79 in the CCBH4851 GRN. On the other hand, the incoherent type II FFL motifs were 4 in CCBH4851 GRN against 3 in the previous network^(12)^.

### Hubs

Identifying the most influential genes in a gene transcription network is a key step in determining therapeutic targets against an infectious agent. One way to identify possible targets is to identify so-called network hubs. Different definitions for the word hub can be applied in the context of complex network theory: one of them is to verify which vertices have the highest *k-out* degrees in order to identify, in the case of a gene regulatory network, the genes with the greatest influence on target regulation. According to Vandereyken *et al.* ^(27)^, the exact number of interactions that characterizes a hub, also called the degree threshold, differs among studies. Some works show that the minimum number is 5, others mention 8, 10, 20 or even 50. In this work, the degree threshold was defined as the average of the number of connections of all nodes having at least two edges. The application of this procedure results in the cutoff value of 16 connections. Table 1 shows the 30 most influential hubs in the *P. aeruginosa* GRN. After pinpointing the hubs, an analysis was performed to check whether they are interconnected (through direct or indirect interactions) or not. It was observed only two hubs are not interconnected: *np20* and PA4851_19380 (homologous to PA1520). The remaining hubs have a direct (when a hub affects the regulation of another hub) or indirect (when hubs affect the regulation of the same group of target genes) connection to other hubs (Figure 4). Node interactions that are not common among hubs were hidden to better visualization in Figure 4.

**Table 1.** The 30 most influential hubs of the *P. aeruginosa* CCBH4851 GRN.

| GENE | *k-out* | GENE | *k-out* |
| --- | --- | --- | --- |
| *lasR* | 99 | *cbrB* | 31 |
| *fur* | 78 | *algR* | 31 |
| *rpoN* | 63 | *ihf* | 30 |
| *anr* | 49 | *phoP* | 29 |
| *mexT* | 48 | *qscR* | 25 |
| *rhlR* | 44 | *cysB* | 21 |
| *fleQ* | 44 | *exsA* | 20 |
| *algU* | 41 | *psrA* | 20 |
| *pmrA* | 41 | *pprB* | 18 |
| *argR* | 39 | *roxR* | 18 |
| *pvdS* | 38 | *rsaL* | 17 |
| *rpoS* | 35 | *roxS* | 17 |
| *dnr* | 34 | *np20* | 17 |
| *vfr* | 33 | *narL* | 16 |
| *lasI* | 32 | PA4851_19380 | 16 |

**Fig. 4.**
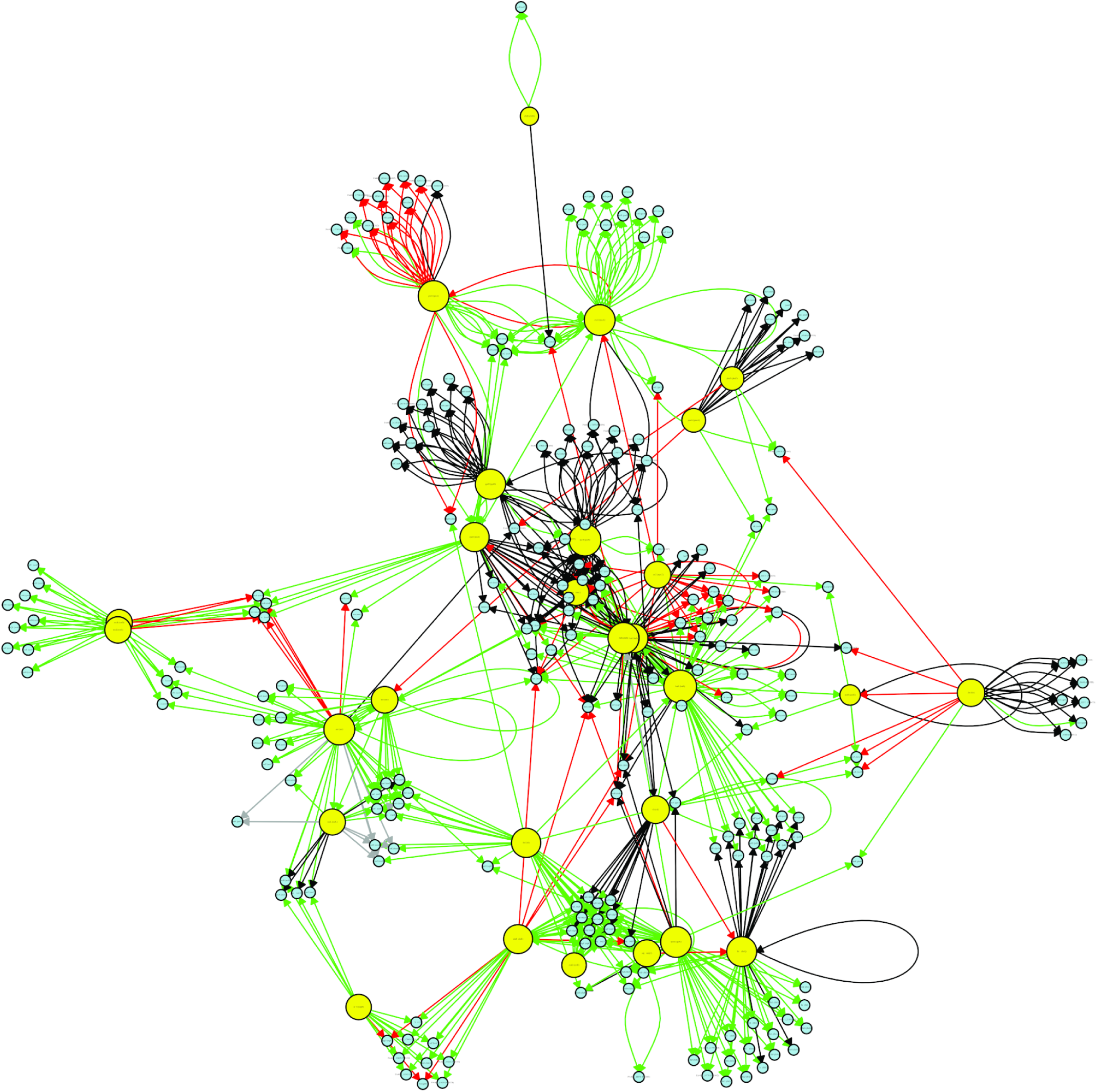
Connectivity relationships among the 30 most influential hubs of the *P. aeruginosa* CCBH4851 GRN. The yellow circles represent regulatory genes considered hubs, light blue circles represent target genes, black lines indicate an unknown mode of regulation, green lines indicate activation, red lines indicate repression and gray lines a dual mode of regulation.

The summarized results comprising network statistics is presented in Table 2, which contains standard measures, such as the number of nodes, number of edges, number of autoregulatory motifs, diameter of the network and average path length. Other relevant measures are the number of coherent and incoherent feed-forward motifs, clustering coefficients, and network entropy. Also, a comparison with data from previous network^(12)^ was included in Table 2.

**Table 2.** Comparison of topological statistic measures between the GRN published by Galán-Vásquez *et al.* ^(12)^ and the *P. aeruginosa* CCBH4851 GRN.

|  | Galán-Vásquez *et al.* <sup>(12)</sup> GRN | CCBH4851 GRN |
| --- | --- | --- |
| Vertices | 690 | 1046 |
| Edges | 1020 | 1576 |
| Regulatory genes | 76 | 138 |
| Target genes | 593 | 908 |
| Positive regulation | 779 | 772 |
| Negative regulation | 218 | 454 |
| Dual regulation | 11 | 13 |
| Unknown regulation | 12 | 337 |
| Autoregulation (total) | 29 | 72 |
| Positive autoregulation | 16 | 21 |
| Negative autoregulation | 13 | 39 |
| Unknown self-regulation | - | 12 |
| Feed-forward loop motifs (total)* | 137 | 218 |
| Coherent type I feed-forward loop motifs* | 82 | 79 |
| Incoherent type II feed-forward loop motifs* | 3 | 4 |
| Density | 2.12e-03 | 1.44e-03 |
| Diameter | 9 | 12 |
| Average path length | 4.08 | 4.80 |
| Global clustering coefficient | 2.28e-02 | 3.2e-02 |
| Local clustering coefficient | 2.5e-01 | 1.92e-01 |
| Entropy | 1.92 | 2.34 |

\*Number of feed-forward loop motifs determined using the igraph package.

## Discussion

The importance of gene regulation on metabolic, adaptive, pathogenic and antibiotic resistance capabilities is well known. The GRN reconstruction and analysis of a versatile pathogen such as *P. aeruginosa*, in particular when based on an MDR strain, contribute to increase the knowledge of related cellular processes. Multidrug resistance can be conferred by a combination of factors varying according to the antimicrobial class. For instance, carbapenems resistance in *P. aeruginosa* is manly given by mutations in *oprD* and/or presence of MBLs. Mutations or differential expression of efflux system genes are also a contributing factor for both carbapenems and aminoglycosides resistance. Overall, multidrug resistance can also be provided by other mechanisms, including acquisition of genes through horizontal transfer and punctual mutations, in multiple combinations, when compared to several non-susceptible strains^(28)^. In addition, *P. aeruginosa* has the ability to form biofilm, which can play a role on antibiotic penetration, antibiotic tolerance, formation of persister cells and protection from the host immune system^(4)^. Due a natural limitation of graph representation, data such as gene expression variation, point mutations or genes lacking experimental evidence are not eligible to be included in a gene regulatory network graph. Overall, we could exclude from our network genes such as *oprD, mexZ, pilA*. Since *oprD* is a target gene, its exclusion has a minor impact in the network topology, but it is extremely important for the cell because *oprD* codifies an outer membrane porin important for the absorption of carbapenems. The lack of OprD leads to low outer membrane permeability. On the other hand, *mexZ* is a regulatory gene and its exclusion from the network results in the exclusion of its node as well as the interactions (edges) with their target genes. The *mexZ* product represses the transcription of *mexX* and *mexY* genes. MexXY are part of an efflux pump system whose overexpression leads to aminoglycoside resistance through the extrusion of this family compounds. We know the MexXY overexpression needs to be experimentally established and cannot be represented in a graph. PilA is a major pilin protein related to bacterial adherence through type VI pilus machinery. Therefore, PilA has great importance in pathogenesis. The *pilA* gene is also a target, only showing regulatory influence upon itself. The advantageous effect of PilA loss to an MDR strain it is unclear. One can hypothesize that this loss could be somehow compensated by newly acquired genes since CCBH4851 has a chromosome approximately 600 kb larger than PAO1. However, these alterations are common among MDR strains^(29,30)^, and to have a network comprising these features could impact dynamics simulations designed to assess MDR bacteria behaviors based on *P. aeruginosa* CCBH4851 GRN. Overall, the reconstruction included additional regulators, target genes, and new interactions described in literature or included in curated databases since the last *P. aeruginosa* GRN publication^(12)^. Several genes involved in virulence mechanisms were identified, such those associated to the production of proteases and toxins, antimicrobial activity, iron uptake, antiphagocytosis, adherence and quorum sensing. Not only new nodes and connections were added, but previously identified nodes were excluded (by the curation process or lack of homology) and interactions were revisited (due to genes that regulatory effect has been recently elucidated). Two noteworthy examples of included nodes and interactions are the regulatory effect of *fleQ* upon “*psl*” genes, and the regulation of efflux pumps genes *mexA, mexE*, and *oprH* by *brlR*. The “*psl*” (for polysaccharide synthesis locus) cluster comprises 15 genes in tandem related to an exopolysaccharide biosynthesis, important to biofilm formation. The recently functionally characterized transcriptional regulator, BrlR, has a biofilm-specific expression and plays a role in the antibiotic tolerance of biofilms through gene expression modulation of efflux pumps genes^(4)^. Altogether, these alterations influenced directly in the network topological characteristics. However, topology measures of the *P. aeruginosa* CCBH4851 GRN, such as node degree distribution and clustering coefficient, remained consistent with a scale-free network type. The degree distribution followed the power-law distribution (Figure 3B and 3D), meaning that a small number of nodes have many connections and a large number of nodes have a few connections. Also, the correlation of local clustering coefficient with node degree (Figure 3F) showed that nodes with lower degrees have larger local clustering coefficients than nodes with higher degrees. Indeed, construction of several networks representing biological processes reveal similar topological characteristics^(24,31)^. As other mathematical aspects of the network topology *per se* were consistent with the type of network obtained, yet a concern is to make sure these measures are consistent with the biological observations. The reconstructed network showed a low-density value compatible with the fact that networks representing natural phenomena often have low density, which is reflected by their structural and dynamic flexibility^(22)^. The low density observed in the CCBH4851 GRN means that the nodes are not all interconnected. Biologically, in an organism such as *P. aeruginosa* that has an average of 6,000 coding sequences, is not expected that all genes maintain an interaction since they are related to distinct biological process that are not all dependent on each other and are triggered in different growth phases, corroborating this low density. In the same way, the global clustering coefficient and connectivity parameters are affected by these biological behaviors, resulting in the large number of connected components found in the CCBH4851 GRN.

Although some nodes under positive regulation were lost (Table 2, the most common regulatory activity found among CCBH4851 GRN interactions was activation. On the other hand, more than 50% of autoregulation found was negative. This could be a consequence of the increase of negative regulation in overall network interactions. A similar pattern was seen in the regulatory network of another member of gammaproteobacteria class, *Escherichia coli*, the prevalence of negative autoregulation in contrast to the prevalence of positive regulation between transcription factors. The positive mode of regulation is important to ensure continuity of biological processes from beginning to end. Adhesion, cell-to-cell signaling, production of virulence and resistance factors, biofilm formation, secretion of toxins, interaction host-pathogen factors are examples of processes that once started must reach a final stage in order to have the desired effect. In fact, we can observe that genes such as *lasR, rlhR, pvdS, anr, dnr, algU* and others involved in these types of process have demonstrated mostly a positive mode of regulation in the CCBH4851 gene regulatory network. On the other hand, negative cycles are important to life-sustaining cyclic processes such as those involved in cell homeostasis. This is the case of metabolic process where we can observe genes such as *lexA, hutC, iscR, desT, mvat* (although involved in virulence factors biosynthesis, this gene regulates arginine metabolism) and others which negative mode of regulation is the predominant effect^(20)^. Regarding the negative autoregulation, they are linked to cellular stability, providing a rapid response to variation of protein/toxin/metabolite concentrations, saving the energetic cost of unneeded synthesis as well as avoiding undesired effects. Some examples of negative autoregulatory interactions included in the CCBH4851 network are *algZ, lexA, metR, ptxR, rsaL* and others. RsaL is a quorum-sensing repressor; LexA is involved in SOS response; AlgZ is the transcriptional activator of AlgD, involved in alginate production; PtxR affects exotoxin A production; MetR is involved in swarming motility and methionine synthesis; overall, these autoregulatory genes tend to be more upstream in the regulatory chain. Dominancy of activation mode was also revealed when looking to network motifs. Motifs are patterns of topological structures statistically overrepresented in the network. The number of FFL motifs, considering all variations, is 218 for the CCBH4851 GRN and 137 for the GRN published by Galán-Vásquez *et al.* ^(12)^. A common motif often related to transcriptional networks, the coherent feed-forward loop, is abundantly present in the CCBH4851 GRN (Table 2)^(26)^. In particular, the coherent type I FFL motif, where all interactions are positive, are common in both GRNs. They act as sign-sensitive delays, *i.e.*, a circuit that responds rapidly to step-like stimuli in one direction (ON to OFF), and at a delay to steps in the opposite direction (OFF to ON). While the temporary removal of the stimulus ceases the transcription, the expression activation needs a persistent signal to carry on. Although less represented, the incoherent type II FFL motif was also found in CCBH4851 GRN. In contrast to the coherent FFL, they act as a sign-sensitive accelerator, *i.e.*, a circuit that responds rapidly to step-like stimuli on one direction but not in the other direction^(26)^. Overall, the FFL motifs are important to modulate cellular processes according to environmental conditions.

One last characteristic revealed by the topological analysis is the presence of hubs. Hubs are nodes showing a large number of connections, a concept that is inherent of scale-free networks. As expected, CCBH4851 GRN analysis pointed out among the most influential hubs genes such as *lasR, fur, anr, mexT, algU*, known to cause great impact in the gene regulatory systems of *P. aeruginosa*. They are involved in resistance, virulence, and pathogenicity mechanisms. LasR, for instance, directly activates the expression of 99 genes. LasR depends on presence and binding of *N*-3-oxo-dodecanoyl-L-homoserine lactone (C12) to act. Once bound, LasR-C12 coordinate the expression of target genes, including many genes encoding virulence factors and cell density^(32)^. In addition, *fur* is the global regulator for iron uptake; *rpoN*, an alternative sigma factor; *mexT*, the regulator of an efflux pump system and several virulence factors; *anr*, responsible for the regulation of anaerobic adaptation processes; all of them known to control the expression of many genes. We could observe that even though few hubs remained unconnected, most of the influential genes belong to the major connected component. This interaction can be direct as the positive effect of *lasR* on *rlhR* transcription, or indirect when hubs are regulating the same targets, *i.e.*, involved in the regulation of the same processes, as *fur* and *algU* both affecting the expression of *phuR* that codifies a member of a heme uptake system to provide host iron acquisition^(33,34)^. Another example is the regulation of *algU, rpoN* and *cysB* over “*alg*” genes, not direct connected but related through their influence in the alginate biosynthesis, important to the mucoid phenotype of *P. aeruginosa* colonies^(34)^. Direct and indirect interactions reflect the importance of influential genes, not only to their specific targets, but the effects of their targets’ regulation upon following processes, triggering a more pleiotropic effect. If a perturbation is required, a hub can affect more than one pathway, resulting in undesired effects. On the other hand, one of the interconnected nodes related to that hub could be an option to perform a perturbation that results in a specific pathway impairment or improvement. Nevertheless, isolated hubs are equally important. In fact, they are related to process such as zinc uptake (*np20*) and purine metabolism (PA4851_19380), that are fundamental to bacterial survival but can be considered somehow independent of other process and are triggered under particular conditions.

Table 2 compared network statistics between the CCBH4851 GRN and the GRN network published by Galán-Vásquez *et al.* ^(12)^. One clear trend is that the CCBH4851 GRN represents a substantial improvement in terms of network completeness, since is includes more nodes, edges and network motifs when compared to the previously published GRN. Other measures reflect this improvement, such as the global clustering coefficient and the diameter of the network. Other comparisons between networks were presented in Figure 3. Those charts and Table 2 show increased completeness and complexity of the CCBH4851 over the previous network, in particular when comparing clustering coefficients (Figures 3E and 3F).

A concept addressed by Csermely^(35)^ is the plasticity of networks. Plastic networks have some interesting characteristics, such as diffuse core, overlapping modules, fewer hierarchies/more loops, large network entropy, and origin dominance, leading to many attractors. Csermely states that biological plastic networks should be attacked by a “central impact” directed at their hubs, bridges and bottlenecks, since if they are attacked on their periphery the effect of the drug will never reach the center of the network due its efficient dissipation. For this reason, topological characteristics as connected components, motifs, hubs are important to determine the best approach to disturb a network in a way to lead the cell to a desired phenotype. Indeed, it is noteworthy that the total network entropy of the CCBH4851 GRN has increased when compared to the *P. aeruginosa* GRN published in 2011 (Table 2). Therefore, the CCBH4851 GRN has increased plasticity when compared to the GRN described in Galán-Vásquez *et al.* ^(12)^. The increased plasticity may be due to the increased size of the CCBH4851 GRN. Nevertheless, one can argue that this observation may be also related to the fact that the CCBH4851 strain is multidrug-resistant, while the previous network is mostly based on the *P. aeruginosa* PAO1.

This reconstruction of *P. aeruginosa* gene regulatory network can contribute to increase our understanding of this bacterium behavior. As future work, we intend to construct a dynamic model of this network, aiming to help researchers working on experimental drug design and screening, to predict dynamical behaviors in order to have a better understanding of the bacteria lifestyle, also allowing the simulation of normal against stress conditions and eventually leading to the discovery of new potential therapeutic targets and the development of new drugs to combat *P. aeruginosa* infections.

## Supporting information

CCBH4851 GRN

Figure 2 (better resolution)

Figure 4 (better resolution)

BBH algorithm

GRN R code

## Acknowledgements

The authors would like to acknowledge INOVA-FIOCRUZ, FAPERJ and CAPES for the financial support.

## Author’s Contribution

FMF performed the GRN reconstruction. APBN and APDCA coordinated the network curation effort. MTS and FABS designed the overall method. All authors have equally participated in the writing of this manuscript.

