## Supplementary material for "Gene regulatory network inference and analysis of multidrug-resistant *Pseudomonas aeruginosa*": CCBH4851 GRN

| Regulatory gene | Ortholog of the regulatory gene | Target gene | Ortholog of the target gene | Mode of regulation | Reference (PubMed) |
|---|---|---|---|---|---|
| aguR | aguR | aguA | aguA | - | 27242034* |
| aguR | aguR | aguB | aguB | - | 27242034* |
| algD | algD | algD | algD | ? | 18440972* |
| algQ | algQ | lasR | lasR | - | 18440972* |
| algQ | algQ | rhlR | rhlR | - | 18440972* |
| algQ | algQ | rpoD | rpoD | + | 18440972* |
| algR | algR | alg44 | alg44 | + | 22587778 |
| algR | algR | alg8 | alg8 | + | 22587778 |
| algR | algR | algA | algA | + | 22587778 |
| algR | algR | algC | algC | + | 22587778 |
| algR | algR | algD | algD | + | 22587778 |
| algR | algR | algE | algE | + | 22587778 |
| algR | algR | algF | algF | + | 22587778 |
| algR | algR | algG | algG | + | 22587778 |
| algR | algR | algI | algI | + | 22587778 |
| algR | algR | algJ | algJ | + | 22587778 |
| algR | algR | algK | algK | + | 22587778 |
| algR | algR | algL | algL | + | 22587778 |
| algR | algR | algR | algR | + | 22587778 |
| algR | algR | algX | algX | + | 22587778 |
| algR | algR | algZ | algZ | - | 22587778 |
| algR | algR | argB | argB | + | 22587778 |
| algR | algR | argG | argG | - | 22587778 |
| algR | algR | braZ | braZ | + | 22587778 |
| algR | algR | gdhA | gdhA | - | 22587778 |
| algR | algR | hcnA | hcnA | - | 22587778 |
| algR | algR | PA4851_01650 | PA0328 | + | 22587778 |
| algR | algR | ldcA | ldcA | + | 22587778 |
| algR | algR | PA4851_17840 | PA1819 | + | 22587778 |
| algR | algR | PA4851_05160 | PA3934 | + | 22587778 |
| algR | algR | PA4851_29405 | PA5152 | + | 22587778 |
| algR | algR | PA4851_29410 | PA5153 | + | 22587778 |
| algR | algR | rhlA | rhlA | - | 22587778 |
| algR | algR | rhlB | rhlB | - | 22587778 |
| algR | algR | rhlI | rhlI | - | 22587778 |
| algR | algR | speA | speA | ? | 22587778 |
| algR | algR | PA4851_16745 | PA2042 | + | 18440972* |
| algU | algU | alg44 | alg44 | + | 18440972* |
| algU | algU | alg8 | alg8 | + | 18440972* |
| algU | algU | algA | algA | + | 18974177* |
| algU | algU | algB | algB | + | 18974177* |
| algU | algU | algD | algD | + | 18440972* |
| algU | algU | algE | algE | + | 18440972* |
| algU | algU | algF | algF | + | 18440972* |
| algU | algU | algG | algG | + | 18440972* |

| Regulatory gene | Ortholog of the regulatory gene | Target gene | Ortholog of the target gene | Mode of regulation | Reference (PubMed) |
|---|---|---|---|---|---|
| algU | algU | algI | algI | + | 18440972* |
| algU | algU | algJ | algJ | + | 18440972* |
| algU | algU | algK | algK | + | 18440972* |
| algU | algU | algL | algL | + | 18974177* |
| algU | algU | algR | algR | + | 18974177* |
| algU | algU | algU | algU | + | 18440972* |
| algU | algU | algX | algX | + | 18974177* |
| algU | algU | algZ | algZ | + | 22587778 |
| algU | algU | bolA | bolA | + | 22587778 |
| algU | algU | dksA | dksA | + | 22587778 |
| algU | algU | fleQ | fleQ | - | 21863142 |
| algU | algU | PA4851_05320 | ivy/ PA3902 | + | 22587778 |
| algU | algU | lptA | lptA | + | 22587778 |
| algU | algU | PA4851_25200 | lptB | + | 22587778 |
| algU | algU | mucA | mucA | + | 22587778 |
| algU | algU | mucB | mucB | + | 22587778 |
| algU | algU | oprF | oprF | + | 22587778 |
| algU | algU | osmC | osmC | + | 22587778 |
| algU | algU | PA4851_23315 | PA0856 | + | 22587778 |
| algU | algU | PA4851_19010 | PA1592 | + | 22587778 |
| algU | algU | PA4851_08455 | PA3262 | + | 22587778 |
| algU | algU | PA4851_05730 | PA3819 | + | 22587778 |
| algU | algU | PA4851_05070 | PA3952 | + | 22587778 |
| algU | algU | PA4851_30120 | PA5291 | + | 22587778 |
| algU | algU | phuR | phuR | + | 22587778 |
| algU | algU | rpoH | rpoH | + | 22587778 |
| algU | algU | PA4851_22040 | PA1053 | + | 22587778 |
| algU | algU | tal | tal | + | 22587778 |
| algU | algU | fusA | fusA | ? | 27242034*, 18974177*, 22587778 |
| algU | algU | PA4851_22040 | PA1053 | + | 27242034* |
| algU | algU | fliC | fliC | ? | 27242034* |
| algU | algU | osmE | osmE | ? | 27242034* |
| algU | algU | amrZ | amrZ | ? | 27242034* |
| algW | algW | algD | algD | + | 12533483 |
| algW | algW | algU | algU | + | 12533483 |
| algZ | algZ | alg44 | alg44 | + | 18440972* |
| algZ | algZ | alg8 | alg8 | + | 18440972* |
| algZ | algZ | algA | algA | + | 18440972* |
| algZ | algZ | algD | algD | + | 18440972* |
| algZ | algZ | algE | algE | + | 18440972* |
| algZ | algZ | algF | algF | + | 18440972* |
| algZ | algZ | algG | algG | + | 18440972* |
| algZ | algZ | algI | algI | + | 18440972* |
| algZ | algZ | algJ | algJ | + | 18440972* |
| algZ | algZ | algK | algK | + | 18440972* |

| Regulatory gene | Ortholog of the regulatory gene | Target gene | Ortholog of the target gene | Mode of regulation | Reference (PubMed) |
|---|---|---|---|---|---|
| algZ | algZ | algL | algL | + | 18440972* |
| algZ | algZ | algX | algX | + | 18440972* |
| algZ | algZ | algZ | algZ | - | 18440972* |
| algZ | algZ | fleQ | fleQ | - | 18440972* |
| ampR | ampR | ampC | ampC | + | 22587778 |
| ampR | ampR | lasA | lasA | - | 22587778 |
| ampR | ampR | lasB | lasB | + | 22587778 |
| ampR | ampR | lasI | lasI | - | 22587778 |
| ampR | ampR | lasR | lasR | - | 22587778 |
| ampR | ampR | poxB | poxB | - | 22587778 |
| ampR | ampR | rhlR | rhlR | + | 22587778 |
| anr | anr | anr | anr | + | 27242034* |
| anr | anr | arcA | arcA | + | 27242034* |
| anr | anr | arcB | arcB | + | 27242034* |
| anr | anr | arcC | arcC | + | 27242034* |
| anr | anr | azu | azu | + | 27242034* |
| anr | anr | ccoN2 | ccoN2 | + | 27242034* |
| anr | anr | ccoO2 | ccoO2 | + | 27242034* |
| anr | anr | ccoP2 | ccoP2 | + | 27242034* |
| anr | anr | ccoQ2 | ccoQ2 | + | 27242034* |
| anr | anr | cioA | cioA | - | 27242034* |
| anr | anr | cioB | cioB | - | 27242034* |
| anr | anr | coIII | coIII | - | 27242034* |
| anr | anr | coxA | coxA | - | 27242034* |
| anr | anr | coxB | coxB | - | 27242034* |
| anr | anr | dnr | dnr | + | 27242034* |
| anr | anr | hcnA | hcnA | + | 27242034* |
| anr | anr | hcnB | hcnB | + | 27242034* |
| anr | anr | hcnC | hcnC | + | 27242034* |
| anr | anr | hemA | hemA | + | 27242034* |
| anr | anr | hemF | hemF | + | 27242034* |
| anr | anr | hemK | hemK | d | 27242034* |
| anr | anr | hemN | hemN | + | 27242034* |
| anr | anr | moeB | moeB | d | 27242034* |
| anr | anr | murI | murI | d | 27242034* |
| anr | anr | narG | narG | + | 27242034* |
| anr | anr | narH | narH | + | 27242034* |
| anr | anr | narI | narI | + | 27242034* |
| anr | anr | narJ | narJ | + | 27242034* |
| anr | anr | narK1 | narK1 | + | 27242034* |
| anr | anr | narK2 | narK2 | + | 27242034* |
| anr | anr | narL | narL | + | 27242034* |
| anr | anr | narX | narX | + | 27242034* |
| anr | anr | nirQ | nirQ | + | 27242034* |
| anr | anr | nirS | nirS | + | 27242034* |

| Regulatory gene | Ortholog of the regulatory gene | Target gene | Ortholog of the target gene | Mode of regulation | Reference (PubMed) |
|---|---|---|---|---|---|
| anr | anr | norB | norB | + | 27242034* |
| anr | anr | norC | norC | + | 27242034* |
| anr | anr | oprE | oprE | + | 27242034* |
| anr | anr | PA4851_02610 | PA0521 | + | 27242034* |
| anr | anr | PA4851_02615 | PA0522 | + | 27242034* |
| anr | anr | PA4851_02635 | PA0526 | ? | 27242034* |
| anr | anr | PA4851_05190 | PA3928 | ? | 27242034* |
| anr | anr | PA4851_24640 | PA4352 | + | 27242034* |
| anr | anr | prfA | prfA | d | 27242034* |
| anr | anr | arcD | arcD | + | 27242034* |
| anr | anr | aroE | aroE | + | 27242034* |
| anr | anr | PA4851_08205 | PA3309 | ? | 27242034* |
| anr | anr | aer | aer | ? | 27242034* |
| anr | anr | PA4851_16350 | PA2127 | ? | 27242034* |
| anr | anr | PA4851_16360 | PA2126 | ? | 27242034* |
| argR | argR | ldcA | ldcA | + | 18440972* |
| argR | argR | PA4851_17840 | PA1819 | + | 18440972* |
| argR | argR | aotJ | aotJ | + | 18440972* |
| argR | argR | aotM | aotM | + | 18440972* |
| argR | argR | aotP | aotP | + | 18440972* |
| argR | argR | aotQ | aotQ | + | 18440972* |
| argR | argR | arcA | arcA | + | 18440972* |
| argR | argR | arcB | arcB | + | 18440972* |
| argR | argR | arcC | arcC | + | 18440972* |
| argR | argR | arcD | arcD | + | 18440972* |
| argR | argR | argF | argF | - | 18440972* |
| argR | argR | argG | argG | - | 18440972* |
| argR | argR | argR | argR | + | 18440972* |
| argR | argR | aruC | aruC | + | 18440972* |
| argR | argR | aruF | aruF | + | 18440972* |
| argR | argR | braZ | braZ | + | 18440972* |
| argR | argR | carA | carA | - | 18440972* |
| argR | argR | carB | carB | - | 18440972* |
| argR | argR | gdhA | gdhA | - | 18440972* |
| argR | argR | gdhB | gdhB | + | 18440972* |
| argR | argR | gltB | gltB | - | 18440972* |
| argR | argR | gltD | gltD | - | 18440972* |
| argR | argR | greA | greA | - | 18440972* |
| argR | argR | PA4851_01650 | PA0328 | + | 18440972* |
| argR | argR | PA4851_23090 | PA0900 | + | 18440972* |
| argR | argR | PA4851_16745 | PA2042 | + | 18440972* |
| argR | argR | PA4851_07145 | PA3538 | - | 18440972* |
| argR | argR | PA4851_05160 | PA3934 | + | 18440972* |
| argR | argR | PA4851_27360 | PA4754 | - | 18440972* |
| argR | argR | PA4851_27375 | PA4757 | - | 18440972* |

| Regulatory gene | Ortholog of the regulatory gene | Target gene | Ortholog of the target gene | Mode of regulation | Reference (PubMed) |
|---|---|---|---|---|---|
| argR | argR | PA4851_29405 | PA5152 | + | 18440972* |
| argR | argR | PA4851_29410 | PA5153 | + | 18440972* |
| argR | argR | PA4851_29415 | PA5154 | + | 18440972* |
| argR | argR | PA4851_29420 | PA5155 | + | 18440972* |
| argR | argR | PA4851_23135 | PA0891 | + | 18440972* |
| argR | argR | aruG | aruG | + | 18440972* |
| argR | argR | aruB | aruB | + | 18440972* |
| argR | argR | aruD | aruD | + | 18440972* |
| argR | argR | aruE | aruE | + | 18440972* |
| atuR | atuR | atuD | atuD | ? | 27242034* |
| atuR | atuR | atuE | atuE | ? | 27242034* |
| atuR | atuR | atuF | atuF | ? | 27242034* |
| atuR | atuR | atuG | atuG | ? | 27242034* |
| atuR | atuR | atuA | atuA | ? | 27242034* |
| atuR | atuR | atuB | atuB | ? | 27242034* |
| atuR | atuR | atuC | atuC | ? | 27242034* |
| atuR | atuR | atuH | atuH | ? | 27242034* |
| bexR | bexR | aprA | aprA | + | 20041030 |
| bexR | bexR | bexR | bexR | + | 20041030 |
| bexR | bexR | PA4851_02865 | PA0572 | + | 20041030 |
| bexR | bexR | PA4851_21290 | PA1202 | + | 20041030 |
| bexR | bexR | PA4851_21285 | PA1203 | + | 20041030 |
| bexR | bexR | PA4851_21280 | PA1204 | + | 20041030 |
| bexR | bexR | PA4851_21275 | PA1205 | + | 20041030 |
| birA | birA | bioB | bioB | - | 27242034* |
| birA | birA | bioF | bioF | - | 27242034* |
| birA | birA | PA4851_02515 | bioH | - | 27242034* |
| birA | birA | bioC | bioC | - | 27242034* |
| birA | birA | bioD | bioD | - | 27242034* |
| brlR | brlR | brlR | brlR | + | 29967320 |
| brlR | brlR | mexA | mexA | + | 23687276 |
| brlR | brlR | mexE | mexE | + | 23687276 |
| brlR | brlR | oprH | oprH | - | 23935054 |
| cbrB | cbrB | aotJ | aotJ | + | 22587778 |
| cbrB | cbrB | aotM | aotM | + | 22587778 |
| cbrB | cbrB | aotP | aotP | + | 22587778 |
| cbrB | cbrB | aotQ | aotQ | + | 22587778 |
| cbrB | cbrB | cbrB | cbrB | + | 22587778 |
| cbrB | cbrB | spuA | spuA | + | 22587778 |
| cbrB | cbrB | spuB | spuB | + | 22587778 |
| cbrB | cbrB | spuC | spuC | + | 22587778 |
| cbrB | cbrB | spuD | spuD | + | 22587778 |
| cbrB | cbrB | spuE | spuE | + | 22587778 |
| cbrB | cbrB | spuF | spuF | + | 22587778 |
| cbrB | cbrB | spuG | spuG | + | 22587778 |

| Regulatory gene | Ortholog of the regulatory gene | Target gene | Ortholog of the target gene | Mode of regulation | Reference (PubMed) |
|---|---|---|---|---|---|
| cbrB | cbrB | spuH | spuH | + | 22587778 |
| cbrB | cbrB | spuI | spuI | + | 22587778 |
| cbrB | cbrB | PA4851_29115 | PA5104 | ? | 27242034* |
| cbrB | cbrB | PA4851_29100 | PA5101 | ? | 27242034* |
| cbrB | cbrB | PA4851_29110 | PA5103 | ? | 27242034* |
| cbrB | cbrB | PA4851_29105 | PA5102 | ? | 27242034* |
| cbrB | cbrB | aotJ | aotJ | + | 27242034* |
| cbrB | cbrB | PA4851_29090 | PA5099 | ? | 27242034* |
| cbrB | cbrB | PA4851_29060 | PA5093 | ? | 27242034* |
| cbrB | cbrB | PA4851_29080 | PA5097 | ? | 27242034* |
| cbrB | cbrB | PA4851_29075 | PA5096 | ? | 27242034* |
| cbrB | cbrB | PA4851_29070 | PA5095 | ? | 27242034* |
| cbrB | cbrB | PA4851_29065 | PA5094 | ? | 27242034* |
| cbrB | cbrB | hutU | hutU | ? | 27242034* |
| cbrB | cbrB | hutI | hutI | ? | 27242034* |
| cbrB | cbrB | hutH | hutH | ? | 27242034* |
| cbrB | cbrB | exoS | exoS | ? | 27242034* |
| cbrB | cbrB | hutG | hutG | ? | 27242034* |
| cbrB | cbrB | hutC | hutC | ? | 27242034* |
| cdhR | PSPA7_RS29445 | PA4851_30635 | PSPA7_RS29440 (PSPA7_6174) | ? | 27242034* |
| cdhR | PSPA7_RS29445 | cdhC | PSPA7_RS29435 (PSPA7_6173) | ? | 27242034* |
| cifR | cifR | morB | morB | ? | 27242034* |
| cifR | cifR | PA4851_10135 | PA2933 | ? | 27242034* |
| cifR | cifR | cif | cif | ? | 27242034* |
| copR | copR | PA4851_13705 | PA2524 | ? | 27242034* |
| copR | copR | PA4851_13710 | PA2523 | ? | 27242034* |
| copR | copR | czcB | czcB | ? | 27242034* |
| copR | copR | czcC | czcC | ? | 27242034* |
| copR | copR | czcA | czcA | ? | 27242034* |
| copR | copR | ptrA | ptrA | ? | 27242034* |
| crc | crc | zwf | zwf | ? | 30429516 |
| crc | crc | bkdB | bkdB | ? | 30429516 |
| crc | crc | mtlD | mtlD | ? | 30429516 |
| crc | crc | bkdA1 | bkdA1 | ? | 30429516 |
| crc | crc | bkdA2 | bkdA2 | ? | 30429516 |
| crc | crc | pilB | pilB | ? | 27242034* |
| crc | crc | lpdV | lpdV | ? | 27242034* |
| CueR | CueR | PA4851_05230 | PA3920 | + | 24175918* |
| CueR | CueR | PA4851_07220 | PA3523 | + | 24175918* |
| CueR | CueR | PA4851_07225 | PA3522 | + | 24175918* |
| CueR | CueR | PA4851_07230 | PA3521 | + | 24175918* |
| cysB | cysB | alg44 | alg44 | + | 22587778 |
| cysB | cysB | alg8 | alg8 | + | 22587778 |
| cysB | cysB | algA | algA | + | 22587778 |
| cysB | cysB | algD | algD | + | 22587778 |

| Regulatory gene | Ortholog of the regulatory gene | Target gene | Ortholog of the target gene | Mode of regulation | Reference (PubMed) |
|---|---|---|---|---|---|
| cysB | cysB | algE | algE | + | 22587778 |
| cysB | cysB | algF | algF | + | 22587778 |
| cysB | cysB | algG | algG | + | 22587778 |
| cysB | cysB | algI | algI | + | 22587778 |
| cysB | cysB | algJ | algJ | + | 22587778 |
| cysB | cysB | algK | algK | + | 22587778 |
| cysB | cysB | algL | algL | + | 22587778 |
| cysB | cysB | algX | algX | + | 22587778 |
| cysB | cysB | aruG | aruG | + | 22587778 |
| cysB | cysB | aruB | aruB | + | 22587778 |
| cysB | cysB | aruC | aruC | + | 22587778 |
| cysB | cysB | cysB | cysB | - | 22587778 |
| cysB | cysB | PA4851_14870 | PA2355 | + | 22587778 |
| cysB | cysB | msuD | msuD | + | 22587778 |
| cysB | cysB | msuE | msuE | + | 22587778 |
| cysB | cysB | PA4851_00950 | PA0185 | ? | 27242034* |
| cysB | cysB | atsA | atsA | ? | 27242034* |
| desT | desT | fabA | fabA | - | 27242034* |
| desT | desT | PA4851_28040 | PA4889 | - | 27242034* |
| desT | desT | desB | desB | - | 27242034* |
| desT | desT | desT | desT | ? | 27242034* |
| dnr | dnr | anr | anr | + | 22587778 |
| dnr | dnr | aroE | aroE | + | 22587778 |
| dnr | dnr | dnr | dnr | + | 22587778 |
| dnr | dnr | hemA | hemA | + | 22587778 |
| dnr | dnr | hemF | hemF | + | 22587778 |
| dnr | dnr | hemN | hemN | + | 22587778 |
| dnr | dnr | narG | narG | + | 22587778 |
| dnr | dnr | narH | narH | + | 22587778 |
| dnr | dnr | narI | narI | + | 22587778 |
| dnr | dnr | narJ | narJ | + | 22587778 |
| dnr | dnr | narK1 | narK1 | + | 22587778 |
| dnr | dnr | narK2 | narK2 | + | 22587778 |
| dnr | dnr | narL | narL | + | 22587778 |
| dnr | dnr | narX | narX | + | 22587778 |
| dnr | dnr | nirC | nirC | + | 22587778 |
| dnr | dnr | nirD | nirD | + | 22587778 |
| dnr | dnr | nirF | nirF | + | 22587778 |
| dnr | dnr | PA4851_02565 | nirH | + | 22587778 |
| dnr | dnr | nirJ | nirJ | + | 22587778 |
| dnr | dnr | nirL | nirL | + | 22587778 |
| dnr | dnr | nirM | nirM | + | 22587778 |
| dnr | dnr | nirN | nirN | + | 22587778 |
| dnr | dnr | nirQ | nirQ | + | 22587778 |
| dnr | dnr | nirS | nirS | + | 22587778 |

| Regulatory gene | Ortholog of the regulatory gene | Target gene | Ortholog of the target gene | Mode of regulation | Reference (PubMed) |
|---|---|---|---|---|---|
| dnr | dnr | norB | norB | + | 22587778 |
| dnr | dnr | norC | norC | + | 22587778 |
| dnr | dnr | nosD | nosD | + | 22587778 |
| dnr | dnr | nosF | nosF | + | 22587778 |
| dnr | dnr | nosL | nosL | + | 22587778 |
| dnr | dnr | nosR | nosR | + | 22587778 |
| dnr | dnr | nosY | nosY | + | 22587778 |
| dnr | dnr | nosZ | nosZ | + | 22587778 |
| dnr | dnr | PA4851_02555 | PA0510 | + | 22587778 |
| dnr | dnr | PA4851_02570 | PA0513 | + | 22587778 |
| erbR | agmR | exaA | exaA | + | 27242034* |
| erbR | agmR | exaB | exaB | + | 27242034* |
| erbR | agmR | exaC | exaC | + | 27242034* |
| erbR | agmR | eraS | exaD | + | 27242034* |
| erbR | agmR | eraR | exaE | + | 27242034* |
| erbR | agmR | pqqA | pqqA | + | 27242034* |
| erbR | agmR | pqqB | pqqB | + | 27242034* |
| erbR | agmR | pqqC | pqqC | + | 27242034* |
| erbR | agmR | pqqD | pqqD | + | 27242034* |
| erbR | agmR | pqqE | pqqE | + | 27242034* |
| erbR | agmR | pqqH | pqqH | + | 27242034* |
| exsA | exsA | exoS | exoS | + | 18440972* |
| exsA | exsA | exsA | exsA | + | 18440972* |
| exsA | exsA | exsB | exsB | + | 18440972* |
| exsA | exsA | exsC | exsC | + | 18440972* |
| exsA | exsA | exsD | exsD | + | 18440972* |
| exsA | exsA | exsE | exsE | + | 18440972* |
| exsA | exsA | PA4851_05610 | PA3842 | + | 18440972* |
| exsA | exsA | PA4851_05605 | PA3843 | + | 18440972* |
| exsA | exsA | pscB | pscB | + | 18440972* |
| exsA | exsA | pscC | pscC | + | 18440972* |
| exsA | exsA | pscD | pscD | + | 18440972* |
| exsA | exsA | pscE | pscE | + | 18440972* |
| exsA | exsA | pscF | pscF | + | 18440972* |
| exsA | exsA | pscG | pscG | + | 18440972* |
| exsA | exsA | pscH | pscH | + | 18440972* |
| exsA | exsA | pscI | pscI | + | 18440972* |
| exsA | exsA | pscJ | pscJ | + | 18440972* |
| exsA | exsA | pscL | pscL | + | 18440972* |
| exsA | exsA | exoS | exoS | + | 27242034* |
| exsA | exsA | exoT | exoT | ? | 27242034* |
| exsD | exsD | exsA | exsA | - | 27242034* |
| fleQ | fleQ | fleR | fleR | + | 22587778 |
| fleQ | fleQ | fleS | fleS | + | 22587778 |
| fleQ | fleQ | flhA | flhA | + | 22587778 |

| Regulatory gene | Ortholog of the regulatory gene | Target gene | Ortholog of the target gene | Mode of regulation | Reference (PubMed) |
|---|---|---|---|---|---|
| fleQ | fleQ | flhB | flhB | + | 22587778 |
| fleQ | fleQ | fliD | fliD | + | 22587778 |
| fleQ | fleQ | fliE | fliE | + | 22587778 |
| fleQ | fleQ | fliF | fliF | + | 22587778 |
| fleQ | fleQ | fliG | fliG | + | 22587778 |
| fleQ | fleQ | PA4851_21790 | fliH | + | 22587778 |
| fleQ | fleQ | fliI | fliI | + | 22587778 |
| fleQ | fleQ | fliJ | fliJ | + | 22587778 |
| fleQ | fleQ | fliM | fliM | + | 22587778 |
| fleQ | fleQ | fliN | fliN | + | 22587778 |
| fleQ | fleQ | fliO | fliO | + | 22587778 |
| fleQ | fleQ | fliP | fliP | + | 22587778 |
| fleQ | fleQ | fliQ | fliQ | + | 22587778 |
| fleQ | fleQ | fliR | fliR | + | 22587778 |
| fleQ | fleQ | PA4851_21830 | fliS | + | 22587778 |
| fleQ | fleQ | fur | fur | - | 22587778 |
| fleQ | fleQ | pelA | pelA | - | 22587778 |
| fleQ | fleQ | pelB | pelB | - | 22587778 |
| fleQ | fleQ | fleQ | fleQ | ? | 22587778 |
| fleQ | fleQ | PA4851_19770 | PA1442 | + | 22587778 |
| fleQ | fleQ | PA4851_21825 | PA1096 | ? | 27242034* |
| fleQ | fleQ | pslI | pslI | ? | 27242034* |
| fleQ | fleQ | flhF | flhF | ? | 27242034* |
| fleQ | fleQ | pelE | pelE | ? | 27242034* |
| fleQ | fleQ | pelD | pelD | ? | 27242034* |
| fleQ | fleQ | pelG | pelG | ? | 27242034* |
| fleQ | fleQ | pelF | pelF | ? | 27242034* |
| fleQ | fleQ | pelC | pelC | ? | 27242034* |
| fleQ | fleQ | PA4851_14440 | PA2441 | ? | 27242034* |
| fleQ | fleQ | pslK | pslK | ? | 27242034* |
| fleQ | fleQ | pslJ | pslJ | ? | 27242034* |
| fleQ | fleQ | pslH | pslH | ? | 27242034* |
| fleQ | fleQ | pslL | pslL | ? | 27242034* |
| fleQ | fleQ | pslC | pslC | ? | 27242034* |
| fleQ | fleQ | pslB | pslB | ? | 27242034* |
| fleQ | fleQ | pslA | pslA | ? | 27242034* |
| fleQ | fleQ | pslG | pslG | ? | 27242034* |
| fleQ | fleQ | pslF | pslF | ? | 27242034* |
| fleQ | fleQ | pslE | pslE | ? | 27242034* |
| fleQ | fleQ | fleN | fleN | ? | 27242034* |
| fleQ | fleQ | pslD | pslD | ? | 27242034* |
| flgM | flgM | fliA | fliA | - | 22587778 |
| flgM | flgM | fliC | fliC | - | 22587778 |
| fliA | fliA | flgM | flgM | + | 22587778 |
| fliA | fliA | fliC | fliC | + | 22587778 |

| Regulatory gene | Ortholog of the regulatory gene | Target gene | Ortholog of the target gene | Mode of regulation | Reference (PubMed) |
|---|---|---|---|---|---|
| fliA | fliA | PA4851_19770 | PA1442 | + | 22587778 |
| fliA | fliA | PA4851_21840 | PA1093 | ? | 27242034* |
| fliA | fliA | PA4851_07990 | PA3352 | ? | 27242034* |
| fliA | fliA | fliA | fliA | ? | 27242034* |
| fpvR | fpvR | fpvI | fpvI | - | 22587778 |
| fpvR | fpvR | pvdS | pvdS | - | 22587778 |
| fruR | fruR | fruR | fruR | - | 27242034* |
| fruR | fruR | fruK | fruK | - | 27242034* |
| fruR | fruR | fruA | fruA | - | 27242034* |
| fruR | fruR | fruI | fruI | - | 27242034* |
| fur | fur | PA4851_25250 | fagA | - | 22587778 |
| fur | fur | foxI | foxI | - | 22587778 |
| fur | fur | foxR | foxR | - | 22587778 |
| fur | fur | fptA | fptA | + | 22587778 |
| fur | fur | PA4851_03740 | fptB | + | 22587778 |
| fur | fur | fpvR | fpvR | + | 22587778 |
| fur | fur | gbuR | gbuR | - | 22587778 |
| fur | fur | hasAp_P | hasAp_P | + | 22587778 |
| fur | fur | hasAp_N | hasAp_N | - | 22587778 |
| fur | fur | icmP | icmP | + | 22587778 |
| fur | fur | oprL | oprL | - | 22587778 |
| fur | fur | motD | motD | - | 22587778 |
| fur | fur | PA4851_00380 | PA0071 | - | 22587778 |
| fur | fur | PA4851_00385 | PA0072 | - | 22587778 |
| fur | fur | PA4851_22680 | PA0929 | - | 22587778 |
| fur | fur | PA4851_22675 | PA0930 | - | 22587778 |
| fur | fur | PA4851_20800 | PA1300 | - | 22587778 |
| fur | fur | PA4851_20795 | PA1301 | - | 22587778 |
| fur | fur | PA4851_16785 | PA2033 | + | 22587778 |
| fur | fur | PA4851_16780 | PA2034 | + | 22587778 |
| fur | fur | PA4851_07685 | PA3409 | + | 22587778 |
| fur | fur | PA4851_07185 | PA3530 | + | 22587778 |
| fur | fur | PA4851_05335 | PA3899 | - | 22587778 |
| fur | fur | PA4851_05330 | PA3900 | - | 22587778 |
| fur | fur | PA4851_25230 | PA4467 | - | 22587778 |
| fur | fur | PA4851_25240 | PA4469 | - | 22587778 |
| fur | fur | PA4851_25475 | PA4516 | + | 22587778 |
| fur | fur | PA4851_26350 | PA4570 | + | 22587778 |
| fur | fur | PA4851_28070 | PA4895 | + | 22587778 |
| fur | fur | PA4851_28075 | PA4896 | + | 22587778 |
| fur | fur | PA4851_29740 | PA5216 | + | 22587778 |
| fur | fur | PA4851_29745 | PA5217 | + | 22587778 |
| fur | fur | pchA | pchA | - | 22587778 |
| fur | fur | pchB | pchB | - | 22587778 |
| fur | fur | pchC | pchC | - | 22587778 |

| Regulatory gene | Ortholog of the regulatory gene | Target gene | Ortholog of the target gene | Mode of regulation | Reference (PubMed) |
|---|---|---|---|---|---|
| fur | fur | pchD | pchD | - | 22587778 |
| fur | fur | pchE | pchE | - | 22587778 |
| fur | fur | pchF | pchF | - | 22587778 |
| fur | fur | pchR | pchR | - | 22587778 |
| fur | fur | pfeR | pfeR | - | 22587778 |
| fur | fur | phuT | phuT | + | 22587778 |
| fur | fur | hemO | hemO | + | 22587778 |
| fur | fur | PA4851_25470 | piuC | + | 22587778 |
| fur | fur | pvdQ | pvdQ | + | 22587778 |
| fur | fur | pvdS | pvdS | - | 22587778 |
| fur | fur | rplA | rplA | + | 22587778 |
| fur | fur | rplJ | rplJ | + | 22587778 |
| fur | fur | rplL | rplL | + | 22587778 |
| fur | fur | sodM | sodA | - | 22587778 |
| fur | fur | tolA | tolA | - | 22587778 |
| fur | fur | tolB | tolB | - | 22587778 |
| fur | fur | tolQ | tolQ | - | 22587778 |
| fur | fur | tolR | tolR | - | 22587778 |
| fur | fur | toxA | toxA | - | 22587778 |
| fur | fur | PA4851_22485 | ybgC | ? | 22587778 |
| fur | fur | PA4851_27090 | PSPA7_RS25870 | - | 22587778 |
| fur | fur | fpvI | fpvI | - | 22587778 |
| fur | fur | fumC1 | fumC1 | - | 22587778 |
| fur | fur | hasR | hasR | + | 22587778 |
| fur | fur | PA4851_05610 | PA3842 | - | 22587778 |
| fur | fur | phuR | phuR | + | 22587778 |
| fur | fur | PA4851_27105 | PA4709 | + | 22587778 |
| fur | fur | PA4851_27095 | PA4707 | + | 22587778 |
| fur | fur | PA4851_27085 | PA4705 | + | 22587778 |
| fur | fur | toxR | toxR | - | 22587778 |
| fur | fur | PA4851_26895 | PA4675 | ? | 27242034* |
| fur | fur | PA4851_20690 | PA1322 | ? | 27242034* |
| fur | fur | aprA | aprA | ? | 27242034* |
| fur | fur | PA4851_25460 | PA4513 | ? | 27242034* |
| fur | fur | PA4851_07680 | PA3410 | ? | 27242034* |
| fur | fur | PA4851_02375 | PA0473 | ? | 27242034* |
| fur | fur | PA4851_26895 | PA4675 | ? | 27242034* |
| fur | fur | PA4851_20690 | PA1322 | ? | 27242034* |
| fur | fur | aprA | aprA | ? | 27242034* |
| fur | fur | PA4851_25460 | PA4513 | ? | 27242034* |
| fur | fur | PA4851_07680 | PA3410 | ? | 27242034* |
| fur | fur | hemO | hemO | ? | 27242034* |
| fur | fur | PA4851_02375 | PA0473 | ? | 27242034* |
| gacA | gacA | hcnA | hcnA | + | 22587778 |
| gacA | gacA | hcnB | hcnB | + | 22587778 |

| Regulatory gene | Ortholog of the regulatory gene | Target gene | Ortholog of the target gene | Mode of regulation | Reference (PubMed) |
|---|---|---|---|---|---|
| gacA | gacA | hcnC | hcnC | + | 22587778 |
| gacA | gacA | lasR | lasR | + | 22587778 |
| gacA | gacA | PA4851_02645 | RsmY | + | 22587778 |
| gacA | gacA | PA4851_06725 | RsmZ | + | 22587778 |
| gacA | gacA | lasI | lasI | ? | 22587778 |
| gacS | gacS | PA4851_02645 | RsmY | + | 27242034* |
| gacS | gacS | PA4851_06725 | RsmZ | + | 27242034* |
| gbdR | PSPA7_RS29400 | PA4851_30680 | PSPA7_RS29485 (PSPA7_6184) | ? | 27242034* |
| gbdR | PSPA7_RS29400 | soxB | soxB | ? | 27242034* |
| gbdR | PSPA7_RS29400 | soxA | soxA | ? | 27242034* |
| gbdR | PSPA7_RS29400 | soxG | soxG | ? | 27242034* |
| gbdR | PSPA7_RS29400 | dgcA | PSPA7_RS29490 (PSPA7_6185) | ? | 27242034* |
| gbdR | PSPA7_RS29400 | gbcA | PSPA7_RS29550 (PSPA7_6197) | ? | 27242034* |
| gbdR | PSPA7_RS29400 | PA4851_30675 | PSPA7_RS29480 (PSPA7_6182) | ? | 27242034* |
| gbuR | gbuR | glmS | glmS | ? | 27242034* |
| gbuR | gbuR | gpuP | gpuP | + | 27242034* |
| gbuR | gbuR | gpuA | gpuA | d | 27242034* |
| gbuR | gbuR | ptxS | ptxS | + | 27242034* |
| gbuR | gbuR | gbuA | gbuA | d | 27242034* |
| gbuR | gbuR | glpR | glpR | + | 27242034* |
| GlcC | GlcC | glcD | glcD | - | 27242034* |
| GlcC | GlcC | glcE | glcE | - | 27242034* |
| GlcC | GlcC | glcF | glcF | - | 27242034* |
| GlcC | GlcC | PA4851_30430 | PA5352 | - | 27242034* |
| GlcC | GlcC | glcC | glcC | - | 27242034* |
| glmR | PSPA7_RS30295 | glmS | glmS | ? | 18440972* |
| glpR | glpR | erbR | agmR | + | 18440972*, 22587778 |
| glpR | glpR | glpD | glpD | - | 22587778, 18440972* |
| glpR | glpR | glpF | glpF | - | 22587778, 18974177* |
| glpR | glpR | PA4851_06940 | glpK1 | ? | 22587778, 18974177* |
| glpR | glpR | glpT | glpT | - | 22587778, 18974177* |
| glpR | glpR | himA | himA | - | 22587778, 18974177* |
| gntR | gntR | PA4851_15035 | PA2322 | - | 27242034* |
| gntR | gntR | PA4851_15040 | PA2321 | - | 27242034* |
| gntR | gntR | gntR | gntR | - | 27242034* |
| gpuR | gpuR | gpuA | gpuA | + | 24175918* |
| gpuR | gpuR | gpuP | gpuP | + | 24175918* |
| gpuR | gpuR | gpuR | gpuR | + | 24175918* |
| HutC | HutC | hutC | hutC | - | 27242034* |
| HutC | HutC | PA4851_29115 | PA5104 | - | 27242034* |
| HutC | HutC | PA4851_29125 | PA5106 | - | 27242034* |
| HutC | HutC | hutU | hutU | - | 27242034* |
| HutC | HutC | PA4851_29090 | PA5099 | - | 27242034* |
| HutC | HutC | hutH | hutH | - | 27242034* |
| HutC | HutC | PA4851_29080 | PA5097 | - | 27242034* |

| Regulatory gene | Ortholog of the regulatory gene | Target gene | Ortholog of the target gene | Mode of regulation | Reference (PubMed) |
|---|---|---|---|---|---|
| HutC | HutC | PA4851_29075 | PA5096 | - | 27242034* |
| HutC | HutC | PA4851_29070 | PA5095 | - | 27242034* |
| HutC | HutC | PA4851_29065 | PA5094 | - | 27242034* |
| HutC | HutC | PA4851_29060 | PA5093 | - | 27242034* |
| HutC | HutC | hutI | hutI | - | 27242034* |
| HutC | HutC | hutG | hutG | - | 27242034* |
| ihf | ihf | alg44 | alg44 | + | 22587778 |
| ihf | ihf | alg8 | alg8 | + | 22587778 |
| ihf | ihf | algA | algA | + | 22587778 |
| ihf | ihf | algB | algB | + | 22587778 |
| ihf | ihf | algD | algD | + | 22587778 |
| ihf | ihf | algE | algE | + | 22587778 |
| ihf | ihf | algF | algF | + | 22587778 |
| ihf | ihf | algG | algG | + | 22587778 |
| ihf | ihf | algI | algI | + | 22587778 |
| ihf | ihf | algJ | algJ | + | 22587778 |
| ihf | ihf | algK | algK | + | 22587778 |
| ihf | ihf | algL | algL | + | 22587778 |
| ihf | ihf | algX | algX | + | 22587778 |
| ihf | ihf | fleR | fleR | + | 22587778 |
| ihf | ihf | fleS | fleS | + | 22587778 |
| ihf | ihf | fliD | fliD | + | 22587778 |
| ihf | ihf | fumC1 | fumC1 | + | 22587778 |
| ihf | ihf | hemA | hemA | + | 22587778 |
| ihf | ihf | hemK | hemK | + | 22587778 |
| ihf | ihf | lasR | lasR | + | 22587778 |
| ihf | ihf | moeB | moeB | + | 22587778 |
| ihf | ihf | murI | murI | + | 22587778 |
| ihf | ihf | narG | narG | + | 22587778 |
| ihf | ihf | narH | narH | + | 22587778 |
| ihf | ihf | narI | narI | + | 22587778 |
| ihf | ihf | narJ | narJ | + | 22587778 |
| ihf | ihf | narK1 | narK1 | + | 22587778 |
| ihf | ihf | narK2 | narK2 | + | 22587778 |
| ihf | ihf | oprE | oprE | + | 22587778 |
| ihf | ihf | algQ | algQ | + | 22587778 |
| iscR | iscR | iscR | iscR | - | 27242034* |
| iscR | iscR | iscS | iscS | - | 27242034* |
| iscR | iscR | iscU | iscU | - | 27242034* |
| iscR | iscR | iscA | iscA | - | 27242034* |
| iscR | iscR | hscB | hscB | - | 27242034* |
| iscR | iscR | hscA | hscA | - | 27242034* |
| iscR | iscR | fdx2 | fdx2 | - | 27242034* |
| iscR | iscR | PA4851_05785 | PA3808 | - | 27242034* |
| iscR | iscR | PA4851_03365 | PA0665 | - | 27242034* |

| Regulatory gene | Ortholog of the regulatory gene | Target gene | Ortholog of the target gene | Mode of regulation | Reference (PubMed) |
|---|---|---|---|---|---|
| lasI | lasI | rsaL | rsaL | ? | 22587778, 27242034* |
| lasI | lasI | xcpP | xcpP | ? | 22587778, 27242034* |
| lasI | lasI | lasB | lasB | ? | 22587778, 27242034* |
| lasI | lasI | lasI | lasI | ? | 22587778, 27242034* |
| lasI | lasI | gacA | gacA | ? | 27242034* |
| lasI | lasI | xcpW | xcpW | ? | 27242034* |
| lasI | lasI | xcpV | xcpV | ? | 27242034* |
| lasI | lasI | rsaL | rsaL | ? | 27242034* |
| lasI | lasI | xcpT | xcpT | ? | 27242034* |
| lasI | lasI | xcpS | xcpS | ? | 27242034* |
| lasI | lasI | xcpR | xcpR | ? | 27242034* |
| lasI | lasI | xcpQ | xcpQ | ? | 27242034* |
| lasI | lasI | xcpP | xcpP | ? | 27242034* |
| lasI | lasI | xcpZ | xcpZ | ? | 27242034* |
| lasI | lasI | xcpX | xcpX | ? | 27242034* |
| lasI | lasI | qscR | qscR | ? | 27242034* |
| lasI | lasI | lasR | lasR | ? | 27242034* |
| lasI | lasI | rhlB | rhlB | ? | 27242034* |
| lasI | lasI | rhlA | rhlA | ? | 27242034* |
| lasI | lasI | xcpU | xcpU | ? | 27242034* |
| lasI | lasI | rhlI | rhlI | ? | 27242034* |
| lasI | lasI | lasA | lasA | ? | 27242034* |
| lasI | lasI | lasB | lasB | ? | 27242034* |
| lasI | lasI | rhlR | rhlR | ? | 27242034* |
| lasI | lasI | ampR | ampR | ? | 27242034* |
| lasI | lasI | lasI | lasI | ? | 27242034* |
| lasI | lasI | fagA | fagA | ? | 27242034* |
| lasI | lasI | PA4851_25240 | PA4469 | ? | 27242034* |
| lasI | lasI | ptxR | ptxR | ? | 27242034* |
| lasI | lasI | pprB | pprB | ? | 27242034* |
| lasI | lasI | eta | eta | ? | 27242034* |
| lasI | lasI | vfr | vfr | ? | 27242034* |
| lasR | lasR | acpP | acpP | + | 22587778, 27242034* |
| lasR | lasR | PA4851_17590 | PA1869 | + | 22587778, 27242034* |
| lasR | lasR | aprD | aprD | + | 22587778, 27242034* |
| lasR | lasR | aprE | aprE | + | 22587778, 27242034* |
| lasR | lasR | aprF | aprF | + | 22587778, 27242034* |
| lasR | lasR | PA4851_21075 | aprX | + | 22587778, 27242034* |
| lasR | lasR | bphO | bphO | + | 22587778, 27242034* |
| lasR | lasR | bphP | bphP | + | 22587778, 27242034* |
| lasR | lasR | PA4851_08120 | PA3326 | + | 22587778, 27242034* |
| lasR | lasR | flp | flp | + | 22587778, 27242034* |
| lasR | lasR | hcnA | hcnA | + | 22587778, 27242034* |
| lasR | lasR | hcnB | hcnB | + | 22587778, 27242034* |
| lasR | lasR | hcnC | hcnC | + | 22587778, 27242034* |

| Regulatory gene | Ortholog of the regulatory gene | Target gene | Ortholog of the target gene | Mode of regulation | Reference (PubMed) |
|---|---|---|---|---|---|
| lasR | lasR | PA4851_17365 | hvn | + | 22587778, 27242034* |
| lasR | lasR | PA4851_31115 | kinB | + | 22587778, 27242034* |
| lasR | lasR | kynB | kynB | + | 22587778, 27242034* |
| lasR | lasR | lasB | lasB | + | 22587778, 27242034* |
| lasR | lasR | lasI | lasI | + | 22587778, 27242034* |
| lasR | lasR | mexR | mexR | + | 22587778, 27242034* |
| lasR | lasR | mvfR | mvfR | + | 22587778, 27242034* |
| lasR | lasR | nuh | nuh | + | 22587778, 27242034* |
| lasR | lasR | PA4851_00165 | PA0027 | + | 22587778, 27242034* |
| lasR | lasR | PA4851_00170 | PA0028 | + | 22587778, 27242034* |
| lasR | lasR | PA4851_00635 | PA0122 | + | 22587778, 27242034* |
| lasR | lasR | PA4851_00745 | PA0144 | + | 22587778, 27242034* |
| lasR | lasR | PA4851_02865 | PA0572 | + | 22587778, 27242034* |
| lasR | lasR | PA4851_23570 | PA0805 | + | 22587778, 27242034* |
| lasR | lasR | PA4851_23320 | PA0855 | + | 22587778, 27242034* |
| lasR | lasR | PA4851_21505 | PA1159 | + | 22587778, 27242034* |
| lasR | lasR | PA4851_19930 | PA1419 | + | 22587778, 27242034* |
| lasR | lasR | PA4851_18685 | PA1656 | + | 22587778, 27242034* |
| lasR | lasR | PA4851_18680 | PA1657 | + | 22587778, 27242034* |
| lasR | lasR | PA4851_18675 | PA1658 | + | 22587778, 27242034* |
| lasR | lasR | PA4851_18670 | PA1659 | + | 22587778, 27242034* |
| lasR | lasR | ambE | ambE | + | 22587778, 27242034* |
| lasR | lasR | ambD | ambD | + | 22587778, 27242034* |
| lasR | lasR | ambC | ambC | + | 22587778, 27242034* |
| lasR | lasR | ambB | ambB | + | 22587778, 27242034* |
| lasR | lasR | PA4851_12265 | PA2588 | + | 22587778, 27242034* |
| lasR | lasR | PA4851_12250 | PA2591 | + | 22587778, 27242034* |
| lasR | lasR | PA4851_10105 | PA2939 | + | 22587778, 27242034* |
| lasR | lasR | PA4851_07160 | PA3535 | + | 22587778, 27242034* |
| lasR | lasR | PA4851_05310 | PA3904 | + | 22587778, 27242034* |
| lasR | lasR | PA4851_05305 | PA3905 | + | 22587778, 27242034* |
| lasR | lasR | PA4851_05300 | PA3906 | + | 22587778, 27242034* |
| lasR | lasR | PA4851_05295 | PA3907 | + | 22587778, 27242034* |
| lasR | lasR | PA4851_05290 | PA3908 | + | 22587778, 27242034* |
| lasR | lasR | PA4851_26905 | PA4677 | + | 22587778, 27242034* |
| lasR | lasR | cueR | cueR | + | 22587778, 27242034* |
| lasR | lasR | PA4851_29555 | PA5181 | + | 22587778, 27242034* |
| lasR | lasR | PA4851_29580 | PA5184 | + | 22587778, 27242034* |
| lasR | lasR | PA4851_29815 | PA5230 | + | 22587778, 27242034* |
| lasR | lasR | PA4851_29820 | PA5231 | + | 22587778, 27242034* |
| lasR | lasR | PA4851_29825 | PA5232 | + | 22587778, 27242034* |
| lasR | lasR | phnC | phnC | + | 22587778, 27242034* |
| lasR | lasR | phzA1 | phzA1 | + | 22587778, 27242034* |
| lasR | lasR | phzB1 | phzB1 | + | 22587778, 27242034* |
| lasR | lasR | phzC1 | phzC1 | + | 22587778, 27242034* |

| Regulatory gene | Ortholog of the regulatory gene | Target gene | Ortholog of the target gene | Mode of regulation | Reference (PubMed) |
|---|---|---|---|---|---|
| lasR | lasR | phzD1 | phzD1 | + | 22587778, 27242034* |
| lasR | lasR | phzE1 | phzE1 | + | 22587778, 27242034* |
| lasR | lasR | phzF1 | phzF1 | + | 22587778, 27242034* |
| lasR | lasR | phzG1 | phzG1 | + | 22587778, 27242034* |
| lasR | lasR | plcB | plcB | + | 22587778, 27242034* |
| lasR | lasR | pqsA | pqsA | + | 22587778, 27242034* |
| lasR | lasR | pqsB | pqsB | + | 22587778, 27242034* |
| lasR | lasR | pqsC | pqsC | + | 22587778, 27242034* |
| lasR | lasR | pqsD | pqsD | + | 22587778, 27242034* |
| lasR | lasR | pqsE | pqsE | + | 22587778, 27242034* |
| lasR | lasR | pqsH | pqsH | + | 22587778, 27242034* |
| lasR | lasR | pslA | pslA | + | 22587778, 27242034* |
| lasR | lasR | pslB | pslB | + | 22587778, 27242034* |
| lasR | lasR | pslC | pslC | + | 22587778, 27242034* |
| lasR | lasR | pslD | pslD | + | 22587778, 27242034* |
| lasR | lasR | pslE | pslE | + | 22587778, 27242034* |
| lasR | lasR | pslF | pslF | + | 22587778, 27242034* |
| lasR | lasR | pslG | pslG | + | 22587778, 27242034* |
| lasR | lasR | pslH | pslH | + | 22587778, 27242034* |
| lasR | lasR | pslI | pslI | + | 22587778, 27242034* |
| lasR | lasR | pslJ | pslJ | + | 22587778, 27242034* |
| lasR | lasR | pslK | pslK | + | 22587778, 27242034* |
| lasR | lasR | pslL | pslL | + | 22587778, 27242034* |
| lasR | lasR | pvdS | pvdS | + | 22587778, 27242034* |
| lasR | lasR | PA4851_17450 | PA1897 | + | 22587778, 27242034* |
| lasR | lasR | rhlG | rhlG | + | 22587778, 27242034* |
| lasR | lasR | rhlI | rhlI | + | 22587778, 27242034* |
| lasR | lasR | rhlR | rhlR | + | 22587778, 27242034* |
| lasR | lasR | rsaL | rsaL | + | 22587778, 27242034* |
| lasR | lasR | tpbA | tpbA | + | 22587778, 27242034* |
| lasR | lasR | xcpP | xcpP | + | 22587778, 27242034* |
| lasR | lasR | xcpQ | xcpQ | + | 22587778, 27242034* |
| lasR | lasR | ambE | ambE | + | 27242034* |
| lasR | lasR | ambB | ambB | + | 27242034* |
| lasR | lasR | ambC | ambC | + | 27242034* |
| lasR | lasR | phzF2 | phzF2 | + | 27242034* |
| lasR | lasR | rhlA | rhlA | ? | 27242034* |
| lasR | lasR | kynU | kynU | ? | 27242034* |
| lasR | lasR | rhlB | rhlB | ? | 27242034* |
| lasR | lasR | PA4851_12245 | PA2592 | ? | 27242034* |
| lasR | lasR | qteE | qteE | ? | 27242034* |
| lexA | lexA | PA4851_00370 | PA0069 | - | 27242034* |
| lexA | lexA | PA4851_22720 | PA0922 | - | 27242034* |
| lexA | lexA | PA4851_24245 | PA0671 | - | 27242034* |
| lexA | lexA | PA4851_24250 | PA0670 | - | 27242034* |

| Regulatory gene | Ortholog of the regulatory gene | Target gene | Ortholog of the target gene | Mode of regulation | Reference (PubMed) |
|---|---|---|---|---|---|
| lexA | lexA | PA4851_24255 | PA0669 | - | 27242034* |
| lexA | lexA | lexA | lexA | - | 27242034* |
| lexA | lexA | PA4851_09750 | PA3008 | - | 27242034* |
| lexA | lexA | recN | recN | - | 27242034* |
| lexA | lexA | recA | recA | - | 27242034* |
| lexA | lexA | PA4851_06755 | PA3616 | - | 27242034* |
| lexA | lexA | PA4851_22080 | PA1045 | - | 27242034* |
| lexA | lexA | PA4851_17610 | PA1865 | - | 27242034* |
| lexA | lexA | PA4851_17605 | PA1866 | - | 27242034* |
| lexA | lexA | PA4851_15200 | PA2288 | - | 27242034* |
| lexA | lexA | PA4851_07665 | PA3413 | - | 27242034* |
| metR | metR | atuF | atuF | + | 27242034*, 18974177*, 22587778 |
| metR | metR | metH | metH | + | 27242034*, 18974177*, 22587778 |
| metR | metR | ntrC | ntrC | - | 27242034*, 18974177*, 22587778 |
| metR | metR | PA4851_03205 | PA0633 | - | 27242034*, 18974177*, 22587778 |
| metR | metR | PA4851_23355 | PA0848 | - | 27242034*, 18974177*, 22587778 |
| metR | metR | PA4851_04120 | PA4144 | + | 27242034*, 18974177*, 22587778 |
| metR | metR | PA4851_03725 | PA4223 | + | 27242034*, 18974177*, 22587778 |
| metR | metR | pvdL | pvdL | + | 27242034*, 18974177*, 22587778 |
| metR | metR | pvdH | pvdH | + | 27242034*, 18974177*, 22587778 |
| metR | metR | metH | metH | - | 27242034*, 18974177*, 22587778 |
| metR | metR | metR | metR | - | 27242034*, 18974177*, 22587778 |
| metR | metR | metE | metE | - | 27242034*, 18974177*, 22587778 |
| mexR | mexR | mexA | mexA | - | 27242034*, 18974177*, 22587778 |
| mexR | mexR | mexB | mexB | - | 27242034*, 18974177*, 22587778 |
| mexR | mexR | mexT | mexT | - | 27242034*, 18974177*, 22587778 |
| mexR | mexR | nalC | nalC | - | 27242034*, 18974177*, 22587778 |
| mexR | mexR | oprM | oprM | - | 27242034*, 18974177*, 22587778 |
| mexT | mexT | cbpD | cbpD | - | 27242034*, 18974177*, 22587778 |
| mexT | mexT | exsA | exsA | - | 27242034*, 18974177*, 22587778 |
| mexT | mexT | fabH2 | fabH2 | - | 27242034*, 18974177*, 22587778 |
| mexT | mexT | hcnA | hcnA | - | 27242034*, 18974177*, 22587778 |
| mexT | mexT | hcnB | hcnB | - | 27242034*, 18974177*, 22587778 |
| mexT | mexT | hcnC | hcnC | - | 27242034*, 18974177*, 22587778 |
| mexT | mexT | lasB | lasB | - | 27242034*, 18974177*, 22587778 |
| mexT | mexT | lldD | lldD | - | 27242034*, 18974177*, 22587778 |
| mexT | mexT | lldP | lldP | - | 27242034*, 18974177*, 22587778 |
| mexT | mexT | mexE | mexE | + | 27242034*, 18974177*, 22587778 |
| mexT | mexT | mexF | mexF | + | 27242034*, 18974177*, 22587778 |
| mexT | mexT | mexT | mexT | + | 27242034*, 18974177*, 22587778 |
| mexT | mexT | oprD | oprD | - | 27242034*, 18974177*, 22587778 |
| mexT | mexT | oprN | oprN | + | 27242034*, 18974177*, 22587778 |
| mexT | mexT | PA4851_18680 | PA1657 | - | 27242034*, 18974177*, 22587778 |
| mexT | mexT | PA4851_18675 | PA1658 | - | 27242034*, 18974177*, 22587778 |
| mexT | mexT | PA4851_18245 | PA1744 | + | 27242034*, 18974177*, 22587778 |

| Regulatory gene | Ortholog of the regulatory gene | Target gene | Ortholog of the target gene | Mode of regulation | Reference (PubMed) |
|---|---|---|---|---|---|
| mexT | mexT | PA4851_17590 | PA1869 | - | 27242034*, 18974177*, 22587778 |
| mexT | mexT | PA4851_17100 | PA1970 | + | 27242034*, 18974177*, 22587778 |
| mexT | mexT | PA4851_14230 | PA2486 | + | 27242034*, 18974177*, 22587778 |
| mexT | mexT | PA4851_11255 | PA2759 | + | 27242034*, 18974177*, 22587778 |
| mexT | mexT | PA4851_10765 | PA2811 | + | 27242034*, 18974177*, 22587778 |
| mexT | mexT | PA4851_10760 | PA2812 | + | 27242034*, 18974177*, 22587778 |
| mexT | mexT | PA4851_10755 | PA2813 | + | 27242034*, 18974177*, 22587778 |
| mexT | mexT | PA4851_08740 | PA3205 | + | 27242034*, 18974177*, 22587778 |
| mexT | mexT | PA4851_08620 | PA3229 | + | 27242034*, 18974177*, 22587778 |
| mexT | mexT | PA4851_08120 | PA3326 | - | 27242034*, 18974177*, 22587778 |
| mexT | mexT | PA4851_08095 | PA3331 | - | 27242034*, 18974177*, 22587778 |
| mexT | mexT | PA4851_08090 | PA3332 | - | 27242034*, 18974177*, 22587778 |
| mexT | mexT | PA4851_04135 | PA4141 | + | 27242034*, 18974177*, 22587778 |
| mexT | mexT | PA4851_24650 | PA4354 | + | 27242034*, 18974177*, 22587778 |
| mexT | mexT | PA4851_24655 | PA4355 | + | 27242034*, 18974177*, 22587778 |
| mexT | mexT | PA4851_26625 | PA4623 | + | 27242034*, 18974177*, 22587778 |
| mexT | mexT | PA4851_27455 | PA4772 | - | 27242034*, 18974177*, 22587778 |
| mexT | mexT | PA4851_28000 | PA4881 | + | 27242034*, 18974177*, 22587778 |
| mexT | mexT | phnA | phnA | - | 27242034*, 18974177*, 22587778 |
| mexT | mexT | phnB | phnB | - | 27242034*, 18974177*, 22587778 |
| mexT | mexT | pqsA | pqsA | - | 27242034*, 18974177*, 22587778 |
| mexT | mexT | pqsB | pqsB | - | 27242034*, 18974177*, 22587778 |
| mexT | mexT | pqsC | pqsC | - | 27242034*, 18974177*, 22587778 |
| mexT | mexT | pqsD | pqsD | - | 27242034*, 18974177*, 22587778 |
| mexT | mexT | pqsE | pqsE | - | 27242034*, 18974177*, 22587778 |
| mexT | mexT | pscE | pscE | - | 27242034*, 18974177*, 22587778 |
| mexT | mexT | pvdA | pvdA | + | 27242034*, 18974177*, 22587778 |
| mexT | mexT | PA4851_14205 | qrh | + | 27242034*, 18974177*, 22587778 |
| mexT | mexT | rhlA | rhlA | - | 27242034*, 18974177*, 22587778 |
| mexT | mexT | rhlI | rhlI | + | 27242034*, 18974177*, 22587778 |
| mexT | mexT | xenB | xenB | + | 27242034*, 18974177*, 22587778 |
| mmsR | mmsR | mmsB | mmsB | ? | 27242034* |
| mmsR | mmsR | mmsA | mmsA | ? | 27242034* |
| mucA | mucA | algU | algU | - | 27242034*, 18974177*, 22587778 |
| mucB | mucB | algU | algU | - | 27242034*, 18974177*, 22587778 |
| mucC | mucC | algU | algU | - | 27242034*, 18974177*, 22587778 |
| mucD | mucD | algU | algU | - | 27242034*, 18974177*, 22587778 |
| mvfR | mvfR | mvfR | mvfR | - | 27242034*, 18974177*, 22587778 |
| mvfR | mvfR | phnA | phnA | + | 27242034*, 18974177*, 22587778 |
| mvfR | mvfR | phnB | phnB | + | 27242034*, 18974177*, 22587778 |
| mvfR | mvfR | pqsA | pqsA | + | 27242034*, 18974177*, 22587778 |
| mvfR | mvfR | pqsB | pqsB | + | 27242034*, 18974177*, 22587778 |
| mvfR | mvfR | pqsC | pqsC | + | 27242034*, 18974177*, 22587778 |
| mvfR | mvfR | pqsD | pqsD | + | 27242034*, 18974177*, 22587778 |
| mvfR | mvfR | pqsE | pqsE | + | 27242034*, 18974177*, 22587778 |

| Regulatory gene | Ortholog of the regulatory gene | Target gene | Ortholog of the target gene | Mode of regulation | Reference (PubMed) |
|---|---|---|---|---|---|
| mvfR | mvfR | rhlI | rhlI | + | 27242034*, 18974177*, 22587778 |
| mvfR | mvfR | rsmA | rsmA | ? | 27242034* |
| mvfR | mvfR | mexH | mexH | ? | 27242034* |
| mvfR | mvfR | opmD | opmD | ? | 27242034* |
| mvfR | mvfR | mexI | mexI | ? | 27242034* |
| mvfR | mvfR | mexG | mexG | ? | 27242034* |
| nalC | nalC | mexA | mexA | - | 27242034*, 18974177*, 22587778 |
| nalC | nalC | mexB | mexB | - | 27242034*, 18974177*, 22587778 |
| nalC | nalC | oprM | oprM | - | 27242034*, 18974177*, 22587778 |
| nalD | nalD | mexB | mexB | ? | 27242034* |
| nalD | nalD | oprM | oprM | ? | 27242034* |
| nalD | nalD | mexA | mexA | ? | 27242034* |
| narL | narL | hemA | hemA | d | 27242034*, 18974177*, 22587778 |
| narL | narL | hemK | hemK | d | 27242034*, 18974177*, 22587778 |
| narL | narL | moeB | moeB | d | 27242034*, 18974177*, 22587778 |
| narL | narL | murI | murI | d | 27242034*, 18974177*, 22587778 |
| narL | narL | narH | narH | + | 27242034*, 18974177*, 22587778 |
| narL | narL | narI | narI | + | 27242034*, 18974177*, 22587778 |
| narL | narL | narJ | narJ | + | 27242034*, 18974177*, 22587778 |
| narL | narL | narK1 | narK1 | + | 27242034*, 18974177*, 22587778 |
| narL | narL | narK2 | narK2 | + | 27242034*, 18974177*, 22587778 |
| narL | narL | nirQ | nirQ | + | 27242034*, 18974177*, 22587778 |
| narL | narL | prfA | prfA | d | 27242034*, 18974177*, 22587778 |
| narL | narL | narG | narG | ? | 27242034*, 18974177*, 22587778 |
| narL | narL | arcD | arcD | ? | 27242034*, 18974177*, 22587778 |
| narL | narL | arcC | arcC | ? | 27242034*, 18974177*, 22587778 |
| narL | narL | arcB | arcB | ? | 27242034*, 18974177*, 22587778 |
| narL | narL | arcA | arcA | ? | 27242034*, 18974177*, 22587778 |
| nfxB | nfxB | oprJ | oprJ | ? | 27242034* |
| nfxB | nfxB | mexD | mexD | ? | 27242034* |
| nfxB | nfxB | mexC | mexC | ? | 27242034* |
| nfxB | nfxB | nfxB | nfxB | - | 27242034* |
| np20 | np20 | np20 | np20 | - | 27242034* |
| np20 | np20 | znuC | znuC | - | 27242034* |
| np20 | np20 | znuB | znuB | - | 27242034* |
| np20 | np20 | PA4851_23690 | PA0781 | - | 27242034* |
| np20 | np20 | PA4851_06830 | PA3601 | - | 27242034* |
| np20 | np20 | PA4851_06835 | PA3600 | - | 27242034* |
| np20 | np20 | PA4851_31375 | PA5536 | - | 27242034* |
| np20 | np20 | PA4851_31370 | PA5535 | - | 27242034* |
| np20 | np20 | PA4851_31365 | PA5534 | - | 27242034* |
| np20 | np20 | PA4851_27785 | PA4838 | - | 27242034* |
| np20 | np20 | PA4851_31380 | PA5537 | - | 27242034* |
| np20 | np20 | PA4851_31390 | PA5539 | - | 27242034* |
| np20 | np20 | PA4851_31395 | PA5540 | - | 27242034* |

| Regulatory gene | Ortholog of the regulatory gene | Target gene | Ortholog of the target gene | Mode of regulation | Reference (PubMed) |
|---|---|---|---|---|---|
| np20 | np20 | pyrQ | pyrQ | - | 27242034* |
| np20 | np20 | amiA | amiA | - | 27242034* |
| np20 | np20 | PA4851_04525 | PA4063 | - | 27242034* |
| np20 | np20 | PA4851_31185 | PA5498 | - | 27242034* |
| NrdR | NrdR | nrdD | nrdD | - | 27242034* |
| NrdR | NrdR | nrdG | nrdG | - | 27242034* |
| NrdR | NrdR | nrdJb | nrdJb | - | 27242034* |
| NrdR | NrdR | nrdJa | nrdJa | - | 27242034* |
| NrdR | NrdR | nrdA | nrdA | - | 27242034* |
| NrdR | NrdR | nrdB | nrdB | - | 27242034* |
| NrdR | NrdR | topA | topA | - | 27242034* |
| NtrC | NtrC | ntrB | ntrB | + | 27242034*, 18974177*, 22587778 |
| NtrC | NtrC | ntrC | ntrC | + | 27242034*, 18974177*, 22587778 |
| NtrC | NtrC | PA4851_18315 | PA1730 | + | 27242034*, 18974177*, 22587778 |
| NtrC | NtrC | PA4851_18310 | PA1731 | + | 27242034*, 18974177*, 22587778 |
| NtrC | NtrC | PA4851_18305 | PA1732 | + | 27242034*, 18974177*, 22587778 |
| NtrC | NtrC | glnA | glnA | + | 27242034*, 18974177*, 22587778 |
| NtrC | NtrC | glnK | glnK | + | 27242034*, 18974177*, 22587778 |
| NtrC | NtrC | amtB | amtB | + | 27242034*, 18974177*, 22587778 |
| ospR | ospR | PA4851_10675 | PA2826 | - | 27242034* |
| oxyR | oxyR | katB | katB | ? | 27242034* |
| oxyR | oxyR | PA4851_23355 | PA0848 | ? | 27242034* |
| oxyR | oxyR | PA4851_26565 | PA4612 | ? | 27242034* |
| PA4851_00625 | PA0120 | dctA | dctA | - | 27242034* |
| PA4851_00625 | PA0120 | PA4851_00625 | PA0120 | - | 27242034* |
| PA4851_00860 | PA0167 | PA4851_19395 | PA1517 | - | 27242034*, 18974177*, 22587778 |
| PA4851_00860 | PA0167 | PA4851_19400 | PA1516 | - | 27242034*, 18974177*, 22587778 |
| PA4851_00860 | PA0167 | alc | alc | - | 27242034*, 18974177*, 22587778 |
| PA4851_00860 | PA0167 | PA4851_19410 | PA1514 | - | 27242034*, 18974177*, 22587778 |
| PA4851_00860 | PA0167 | PA4851_19415 | PA1513 | - | 27242034*, 18974177*, 22587778 |
| PA4851_00860 | PA0167 | PA4851_00705 | PA0136 | - | 27242034*, 18974177*, 22587778 |
| PA4851_00860 | PA0167 | PA4851_00710 | PA0137 | - | 27242034*, 18974177*, 22587778 |
| PA4851_00860 | PA0167 | PA4851_00715 | PA0138 | - | 27242034*, 18974177*, 22587778 |
| PA4851_00860 | PA0167 | PA4851_00850 | PA0165 | - | 27242034*, 18974177*, 22587778 |
| PA4851_00860 | PA0167 | PA4851_00855 | PA0166 | - | 27242034*, 18974177*, 22587778 |
| PA4851_00860 | PA0167 | PA4851_00860 | PA0167 | - | 27242034*, 18974177*, 22587778 |
| PA4851_00860 | PA0167 | PA4851_00865 | PA0168 | - | 27242034*, 18974177*, 22587778 |
| PA4851_01350 | PA0268 | PA4851_01355 | PA0269 | - | 27242034* |
| PA4851_01350 | PA0268 | PA4851_01360 | PA0270 | - | 27242034* |
| PA4851_02195 | PA0436 | PA4851_02230 | PA0443 | - | 27242034* |
| PA4851_02195 | PA0436 | PA4851_02235 | PA0444 | - | 27242034* |
| PA4851_02195 | PA0436 | dht | dht | - | 27242034* |
| PA4851_02195 | PA0436 | PA4851_02215 | PA0440 | - | 27242034* |
| PA4851_02195 | PA0436 | PA4851_02210 | PA0439 | - | 27242034* |
| PA4851_02195 | PA0436 | codB | codB | - | 27242034* |

| Regulatory gene | Ortholog of the regulatory gene | Target gene | Ortholog of the target gene | Mode of regulation | Reference (PubMed) |
|---|---|---|---|---|---|
| PA4851_02195 | PA0436 | codA | codA | - | 27242034* |
| PA4851_02445 | modE | modA | modA | - | 27242034* |
| PA4851_02445 | modE | modB | modB | - | 27242034* |
| PA4851_02445 | modE | modC | modC | - | 27242034* |
| PA4851_02745 | PA0547 | PA4851_02745 | PA0547 | - | 27242034*, 18974177*, 22587778 |
| PA4851_02745 | PA0547 | metK | metK | - | 27242034*, 18974177*, 22587778 |
| PA4851_04015 | PA4165 | PA4851_04010 | PA4166 | - | 27242034* |
| PA4851_04180 | PA4132 | cysI | cysI | - | 27242034* |
| PA4851_04180 | PA4132 | PA4851_04195 | PA4129 | - | 27242034* |
| PA4851_04180 | PA4132 | PA4851_04175 | PA4133 | - | 27242034* |
| PA4851_06045 | PA3757 | PA4851_06045 | PA3757 | - | 27242034* |
| PA4851_06045 | PA3757 | PA4851_06040 | PA3758 | - | 27242034* |
| PA4851_06045 | PA3757 | PA4851_06035 | PA3759 | - | 27242034* |
| PA4851_06045 | PA3757 | PA4851_06030 | PA3760 | - | 27242034* |
| PA4851_06045 | PA3757 | PA4851_06025 | PA3761 | - | 27242034* |
| PA4851_06345 | PA3697 | fhp | fhp | + | 27242034*, 18974177*, 22587778 |
| PA4851_06385 | PA3689 | PA4851_06380 | PA3690 | + | 27242034* |
| PA4851_07825 | PA3381 | PA4851_07825 | PA3381 | - | 27242034* |
| PA4851_07825 | PA3381 | PA4851_07830 | PA3380 | - | 27242034* |
| PA4851_07825 | PA3381 | PA4851_07835 | PA3379 | - | 27242034* |
| PA4851_07825 | PA3381 | PA4851_07840 | PA3378 | - | 27242034* |
| PA4851_07825 | PA3381 | PA4851_07845 | PA3377 | - | 27242034* |
| PA4851_07825 | PA3381 | PA4851_07850 | PA3376 | - | 27242034* |
| PA4851_07825 | PA3381 | PA4851_07855 | PA3375 | - | 27242034* |
| PA4851_07825 | PA3381 | PA4851_07860 | PA3374 | - | 27242034* |
| PA4851_07825 | PA3381 | PA4851_07865 | PA3373 | - | 27242034* |
| PA4851_07825 | PA3381 | PA4851_07870 | PA3372 | - | 27242034* |
| PA4851_08520 | PA3249 | PA4851_08520 | PA3249 | - | 27242034* |
| PA4851_08520 | PA3249 | PA4851_08515 | PA3250 | - | 27242034* |
| PA4851_08520 | PA3249 | PA4851_08510 | PA3251 | - | 27242034* |
| PA4851_08520 | PA3249 | PA4851_08505 | PA3252 | - | 27242034* |
| PA4851_08520 | PA3249 | PA4851_08500 | PA3253 | - | 27242034* |
| PA4851_08520 | PA3249 | PA4851_08495 | PA3254 | - | 27242034* |
| PA4851_08520 | PA3249 | PA4851_08490 | PA3255 | - | 27242034* |
| PA4851_08520 | PA3249 | PA4851_10810 | PA2802 | - | 27242034* |
| PA4851_08520 | PA3249 | PA4851_10805 | PA2803 | - | 27242034* |
| PA4851_08520 | PA3249 | PA4851_10800 | PA2804 | - | 27242034* |
| PA4851_08520 | PA3249 | PA4851_19125 | PA1144 | - | 27242034* |
| PA4851_08520 | PA3249 | PA4851_21585 | PA1143 | - | 27242034* |
| PA4851_08520 | PA3249 | PA4851_21590 | PA1142 | - | 27242034* |
| PA4851_08845 | PA3184 | PA4851_08845 | PA3184 | - | 27242034* |
| PA4851_08845 | PA3184 | edd | edd | - | 27242034* |
| PA4851_08845 | PA3184 | glk | glk | - | 27242034* |
| PA4851_08845 | PA3184 | gltR | gltR | - | 27242034* |
| PA4851_08845 | PA3184 | gltS | gltS | - | 27242034* |

| Regulatory gene | Ortholog of the regulatory gene | Target gene | Ortholog of the target gene | Mode of regulation | Reference (PubMed) |
|---|---|---|---|---|---|
| PA4851_08845 | PA3184 | gapA | gapA | - | 27242034* |
| PA4851_08845 | PA3184 | zwf | zwf | - | 27242034* |
| PA4851_08845 | PA3184 | pgl | pgl | - | 27242034* |
| PA4851_08845 | PA3184 | PA4851_08860 | PA3181 | - | 27242034* |
| PA4851_11865 | fhpR | fhp | fhp | + | 27242034*, 18974177*, 22587778 |
| PA4851_11865 | fhpR | PA4851_11865 | fhpR | - | 27242034*, 18974177*, 22587778 |
| PA4851_11865 | fhpR | ppyR | ppyR | + | 27242034*, 18974177*, 22587778 |
| PA4851_11865 | fhpR | PA4851_11880 | PA2662 | + | 27242034*, 18974177*, 22587778 |
| PA4851_11910 | bqsR/PA2657 | phnA | phnA | + | 27242034*, 18974177*, 22587778 |
| PA4851_11910 | bqsR/PA2657 | pqsA | pqsA | + | 27242034*, 18974177*, 22587778 |
| PA4851_11910 | bqsR/PA2657 | rhlA | rhlA | + | 27242034*, 18974177*, 22587778 |
| PA4851_11910 | bqsR/PA2657 | rhlB | rhlB | + | 27242034*, 18974177*, 22587778 |
| PA4851_11915 | bqsS/PA2656 | phnA | phnA | + | 27242034*, 18974177*, 22587778 |
| PA4851_11915 | bqsS/PA2656 | pqsA | pqsA | + | 27242034*, 18974177*, 22587778 |
| PA4851_11915 | bqsS/PA2656 | rhlA | rhlA | + | 27242034*, 18974177*, 22587778 |
| PA4851_11915 | bqsS/PA2656 | rhlB | rhlB | + | 27242034*, 18974177*, 22587778 |
| PA4851_12250 | PA2591 | pprB | pprB | - | 27242034*, 18974177*, 22587778 |
| PA4851_14400 | PA2449 | gcvH2 | gcvH2 | + | 27242034* |
| PA4851_14400 | PA2449 | gcvP2 | gcvP2 | + | 27242034* |
| PA4851_14400 | PA2449 | glyA2 | glyA2 | + | 27242034* |
| PA4851_14400 | PA2449 | sdaA | sdaA | + | 27242034* |
| PA4851_14400 | PA2449 | gcvT2 | gcvT2 | + | 27242034* |
| PA4851_15145 | PA2299 | PA4851_15145 | PA2299 | - | 27242034* |
| PA4851_15145 | PA2299 | PA4851_15150 | PA2298 | - | 27242034* |
| PA4851_15145 | PA2299 | PA4851_15155 | PA2297 | - | 27242034* |
| PA4851_15145 | PA2299 | PA4851_15160 | PA2296 | - | 27242034* |
| PA4851_15145 | PA2299 | PA4851_15165 | PA2295 | - | 27242034* |
| PA4851_15145 | PA2299 | PA4851_15170 | PA2294 | - | 27242034* |
| PA4851_15145 | PA2299 | PA4851_15175 | PA2293 | - | 27242034* |
| PA4851_15145 | PA2299 | PA4851_15180 | PA2292 | - | 27242034* |
| PA4851_16790 | PA2032 | PA4851_16795 | PA2031 | - | 27242034* |
| PA4851_16900 | PA2010 | PA4851_16900 | PA2010 | - | 27242034* |
| PA4851_16900 | PA2010 | hmgA | hmgA | - | 27242034* |
| PA4851_16900 | PA2010 | fahA | fahA | - | 27242034* |
| PA4851_16900 | PA2010 | maiA | maiA | - | 27242034* |
| PA4851_16900 | PA2010 | PA4851_16920 | PA2006 | - | 27242034* |
| PA4851_19280 | PA1539 | PA4851_19280 | PA1539 | - | 27242034* |
| PA4851_19280 | PA1539 | PA4851_19285 | PA1538 | - | 27242034* |
| PA4851_19280 | PA1539 | PA4851_19290 | PA1537 | - | 27242034* |
| PA4851_19380 | PA1520 | PA4851_10110 | PA2938 | - | 27242034*, 18974177*, 22587778 |
| PA4851_19380 | PA1520 | PA4851_19380 | PA1520 | - | 27242034*, 18974177*, 22587778 |
| PA4851_19380 | PA1520 | PA4851_19390 | PA1518 | - | 27242034*, 18974177*, 22587778 |
| PA4851_19380 | PA1520 | gcl | gcl | - | 27242034*, 18974177*, 22587778 |
| PA4851_19380 | PA1520 | PA4851_19475 | PA1501 | - | 27242034*, 18974177*, 22587778 |
| PA4851_19380 | PA1520 | PA4851_19480 | PA1500 | - | 27242034*, 18974177*, 22587778 |

| Regulatory gene | Ortholog of the regulatory gene | Target gene | Ortholog of the target gene | Mode of regulation | Reference (PubMed) |
|---|---|---|---|---|---|
| PA4851_19380 | PA1520 | PA4851_00850 | PA0165 | - | 27242034*, 18974177*, 22587778 |
| PA4851_19380 | PA1520 | PA4851_19465 | PA1503 | - | 27242034*, 18974177*, 22587778 |
| PA4851_19380 | PA1520 | PA4851_19445 | PA1507 | - | 27242034*, 18974177*, 22587778 |
| PA4851_19380 | PA1520 | PA4851_19395 | PA1517 | - | 27242034*, 18974177*, 22587778 |
| PA4851_19380 | PA1520 | PA4851_19400 | PA1516 | - | 27242034*, 18974177*, 22587778 |
| PA4851_19380 | PA1520 | PA4851_19410 | PA1514 | - | 27242034*, 18974177*, 22587778 |
| PA4851_19380 | PA1520 | PA4851_19415 | PA1513 | - | 27242034*, 18974177*, 22587778 |
| PA4851_19380 | PA1520 | PA4851_02390 | PA0476 | - | 27242034*, 18974177*, 22587778 |
| PA4851_19380 | PA1520 | PA4851_19385 | PA1519 | - | 27242034*, 18974177*, 22587778 |
| PA4851_19380 | PA1520 | alc | alc | - | 27242034*, 18974177*, 22587778 |
| PA4851_19460 | PA1504 | xdhA | xdhA | - | 27242034*, 18974177*, 22587778 |
| PA4851_19460 | PA1504 | xdhB | xdhB | - | 27242034*, 18974177*, 22587778 |
| PA4851_19460 | PA1504 | PA4851_19370 | PA1522 | - | 27242034*, 18974177*, 22587778 |
| PA4851_19460 | PA1504 | PA4851_19375 | PA1521 | - | 27242034*, 18974177*, 22587778 |
| PA4851_19460 | PA1504 | PA4851_19385 | PA1519 | - | 27242034*, 18974177*, 22587778 |
| PA4851_20955 | PA1269 | PA4851_20955 | PA1269 | + | 27242034* |
| PA4851_20955 | PA1269 | PA4851_20960 | PA1268 | + | 27242034* |
| PA4851_20955 | PA1269 | PA4851_20965 | PA1267 | + | 27242034* |
| PA4851_20955 | PA1269 | PA4851_21000 | PA1260 | + | 27242034* |
| PA4851_20955 | PA1269 | PA4851_21005 | PA1259 | + | 27242034* |
| PA4851_20955 | PA1269 | PA4851_21010 | PA1258 | + | 27242034* |
| PA4851_20955 | PA1269 | PA4851_21015 | PA1257 | + | 27242034* |
| PA4851_20955 | PA1269 | PA4851_21020 | PA1256 | + | 27242034* |
| PA4851_20955 | PA1269 | PA4851_21025 | PA1255 | + | 27242034* |
| PA4851_20955 | PA1269 | PA4851_21030 | PA1254 | + | 27242034* |
| PA4851_20955 | PA1269 | PA4851_21035 | PA1253 | + | 27242034* |
| PA4851_22055 | PA1050 | PA4851_22050 | PA1051 | + | 27242034* |
| PA4851_22055 | PA1050 | PA4851_22045 | PA1052 | + | 27242034* |
| PA4851_22615 | pqrR | PA4851_22620 | PA0941 | - | 27242034* |
| PA4851_22615 | pqrR | PA4851_22625 | PA0940 | - | 27242034* |
| PA4851_22615 | pqrR | PA4851_22630 | PA0939 | - | 27242034* |
| PA4851_22615 | pqrR | PA4851_22615 | pqrR | + | 27242034* |
| PA4851_23610 | PA0797 | PA4851_23610 | PA0797 | - | 27242034* |
| PA4851_23610 | PA0797 | prpB | prpB | - | 27242034* |
| PA4851_23610 | PA0797 | prpC | prpC | - | 27242034* |
| PA4851_23610 | PA0797 | PA4851_23625 | PA0794 | - | 27242034* |
| PA4851_23610 | PA0797 | PA4851_23630 | PA0793 | - | 27242034* |
| PA4851_23610 | PA0797 | prpD | prpD | - | 27242034* |
| PA4851_23700 | PA0779 | fhp | fhp | + | 27242034*, 18974177*, 22587778 |
| PA4851_23700 | PA0779 | PA4851_11865 | fhpR | - | 27242034*, 18974177*, 22587778 |
| PA4851_24455 | mvaT | cupA1 | cupA1 | - | 27242034*, 18974177*, 22587778 |
| PA4851_24455 | mvaT | cupB1 | cupB1 | - | 27242034*, 18974177*, 22587778 |
| PA4851_24455 | mvaT | cupC1 | cupC1 | - | 27242034*, 18974177*, 22587778 |
| PA4851_24455 | mvaT | ptxS | ptxS | + | 27242034*, 18974177*, 22587778 |
| PA4851_25390 | psdR | PA4851_25395 | PA4500 | - | 27242034* |

| Regulatory gene | Ortholog of the regulatory gene | Target gene | Ortholog of the target gene | Mode of regulation | Reference (PubMed) |
|---|---|---|---|---|---|
| PA4851_25390 | psdR | PA4851_25410 | dppB | - | 27242034* |
| PA4851_25390 | psdR | PA4851_25415 | dppC | - | 27242034* |
| PA4851_25390 | psdR | PA4851_25420 | dppD | - | 27242034* |
| PA4851_25390 | psdR | PA4851_25425 | dppF | - | 27242034* |
| PA4851_26805 | PA4659 | PA4851_26795 | PA4657 | - | 27242034* |
| PA4851_26805 | PA4659 | PA4851_26800 | PA4658 | - | 27242034* |
| PA4851_26805 | PA4659 | PA4851_26805 | PA4659 | - | 27242034* |
| PA4851_26805 | PA4659 | phr | phr | - | 27242034* |
| PA4851_27440 | PA4769 | PA4851_27440 | PA4769 | - | 27242034*, 18974177*, 22587778 |
| PA4851_27440 | PA4769 | lldP | lldP | - | 27242034*, 18974177*, 22587778 |
| PA4851_27440 | PA4769 | lldD | lldD | - | 27242034*, 18974177*, 22587778 |
| PA4851_27440 | PA4769 | PA4851_27455 | PA4772 | - | 27242034*, 18974177*, 22587778 |
| PA4851_28125 | PA4906 | vanA | vanA | - | 27242034* |
| PA4851_28125 | PA4906 | vanB | vanB | - | 27242034* |
| PA4851_28125 | PA4906 | PA4851_28110 | PA4903 | - | 27242034* |
| PA4851_28175 | PA4916 | nadD | nadD | - | 27242034*, 18974177*, 22587778 |
| PA4851_28175 | PA4916 | PA4851_28175 | PA4916 | - | 27242034*, 18974177*, 22587778 |
| PA4851_28175 | PA4916 | PA4851_28185 | PA4918 | - | 27242034*, 18974177*, 22587778 |
| PA4851_28175 | PA4916 | pncB1 | pncB1 | - | 27242034*, 18974177*, 22587778 |
| PA4851_28175 | PA4916 | nadE | nadE | - | 27242034*, 18974177*, 22587778 |
| PA4851_30850 | PA5431 | PA4851_30855 | PA5432 | - | 27242034* |
| PA4851_30850 | PA5431 | PA4851_30860 | PA5433 | - | 27242034* |
| PA4851_30880 | PSPA7_RS29685 | PA4851_30870 | oadA | ? | 27242034*, 18974177*, 22587778 |
| PA4851_30880 | PSPA7_RS29685 | PA4851_00880 | PSPA7_RS01190 | ? | 27242034*, 18974177*, 22587778 |
| PA4851_30880 | PSPA7_RS29685 | norC | norC | ? | 27242034*, 18974177*, 22587778 |
| PA4851_30880 | PSPA7_RS29685 | PA4851_30880 | PSPA7_RS29685 | ? | 27242034*, 18974177*, 22587778 |
| PA4851_30880 | PSPA7_RS29685 | nosZ | nosZ | ? | 27242034*, 18974177*, 22587778 |
| PA4851_30885 | PA5438 | zwf | zwf | - | 27242034*, 18974177*, 22587778 |
| PA4851_30885 | PA5438 | aceE | aceE | - | 27242034*, 18974177*, 22587778 |
| PA4851_30885 | PA5438 | aceF | aceF | - | 27242034*, 18974177*, 22587778 |
| PA4851_30885 | PA5438 | PA4851_30885 | PA5438 | - | 27242034*, 18974177*, 22587778 |
| PA4851_30885 | PA5438 | PA4851_08840 | PA3185 | - | 27242034*, 18974177*, 22587778 |
| PA4851_31225 | PA5506 | PA4851_31225 | PA5506 | - | 27242034* |
| PA4851_31225 | PA5506 | PA4851_31230 | PA5507 | - | 27242034* |
| PA4851_31225 | PA5506 | PA4851_31235 | PA5508 | - | 27242034* |
| PA4851_31225 | PA5506 | PA4851_31240 | PA5509 | - | 27242034* |
| PA4851_31225 | PA5506 | PA4851_31245 | PA5510 | - | 27242034* |
| pchR | pchR | fptA | fptA | + | 27242034*, 18974177*, 22587778 |
| pchR | pchR | pchA | pchA | + | 27242034*, 18974177*, 22587778 |
| pchR | pchR | pchB | pchB | + | 27242034*, 18974177*, 22587778 |
| pchR | pchR | pchC | pchC | + | 27242034*, 18974177*, 22587778 |
| pchR | pchR | pchD | pchD | + | 27242034*, 18974177*, 22587778 |
| pchR | pchR | pchE | pchE | + | 27242034*, 18974177*, 22587778 |
| pchR | pchR | pchF | pchF | + | 27242034*, 18974177*, 22587778 |
| pchR | pchR | pchR | pchR | - | 27242034*, 18974177*, 22587778 |

| Regulatory gene | Ortholog of the regulatory gene | Target gene | Ortholog of the target gene | Mode of regulation | Reference (PubMed) |
|---|---|---|---|---|---|
| pepA | pepA | algD | algD | ? | 27242034*, 18974177*, 22587778 |
| pfeR | pfeR | pfeA | pfeA | + | 27242034*, 18974177*, 22587778 |
| phhR | phhR | phhA | phhA | + | 27242034* |
| phhR | phhR | phhB | phhB | + | 27242034* |
| phhR | phhR | phhC | phhC | + | 27242034* |
| phhR | phhR | phhR | phhR | - | 27242034* |
| phhR | phhR | hpd | hpd | + | 27242034* |
| phhR | phhR | phhR | phhR | + | 27242034* |
| phhR | phhR | phhA | phhA | + | 27242034* |
| phhR | phhR | phhB | phhB | + | 27242034* |
| phhR | phhR | phhC | phhC | + | 27242034* |
| phoB | phoB | PA4851_27815 | PSPA7_RS26570 (PSPA7_5564) | ? | 27242034* |
| phoB | phoB | phoU | phoU | ? | 27242034* |
| phoP | phoP | PA4851_23420 | PA0836 | - | 27242034*, 18974177*, 22587778 |
| phoP | phoP | gabD | gabD | + | 27242034*, 18974177*, 22587778 |
| phoP | phoP | gabT | gabT | + | 27242034*, 18974177*, 22587778 |
| phoP | phoP | PA4851_07225 | PA3522 | + | 27242034*, 18974177*, 22587778 |
| phoP | phoP | oprH | oprH | + | 27242034*, 18974177*, 22587778 |
| phoP | phoP | PA4851_22725 | PA0921 | + | 27242034*, 18974177*, 22587778 |
| phoP | phoP | PA4851_21320 | PA1196 | - | 27242034*, 18974177*, 22587778 |
| phoP | phoP | PA4851_08205 | PA3309 | - | 27242034*, 18974177*, 22587778 |
| phoP | phoP | PA4851_06585 | PA3649 | + | 27242034*, 18974177*, 22587778 |
| phoP | phoP | PA4851_04785 | PA4010 | + | 27242034*, 18974177*, 22587778 |
| phoP | phoP | PA4851_04780 | PA4011 | + | 27242034*, 18974177*, 22587778 |
| phoP | phoP | PA4851_25160 | PA4453 | + | 27242034*, 18974177*, 22587778 |
| phoP | phoP | PA4851_25165 | PA4454 | + | 27242034*, 18974177*, 22587778 |
| phoP | phoP | PA4851_25170 | PA4455 | + | 27242034*, 18974177*, 22587778 |
| phoP | phoP | PA4851_28185 | PA4918 | - | 27242034*, 18974177*, 22587778 |
| phoP | phoP | phoP | phoP | + | 27242034*, 18974177*, 22587778 |
| phoP | phoP | phoQ | phoQ | + | 27242034*, 18974177*, 22587778 |
| phoP | phoP | sodB | sodB | + | 27242034*, 18974177*, 22587778 |
| phoP | phoP | tpbA | tpbA | + | 27242034*, 18974177*, 22587778 |
| phoP | phoP | ackA | ackA | - | 27242034* |
| phoP | phoP | arnT | arnT | ? | 27242034* |
| phoP | phoP | arnC | arnC | ? | 27242034* |
| phoP | phoP | arnB | arnB | ? | 27242034* |
| phoP | phoP | arnA | arnA | ? | 27242034* |
| phoP | phoP | arnF | arnF | ? | 27242034* |
| phoP | phoP | arnE | arnE | ? | 27242034* |
| phoP | phoP | arnD | arnD | ? | 27242034* |
| phoP | phoP | PA4851_07040 | PA3559 | ? | 27242034* |
| phoP | phoP | PA4851_20580 | PA1343 | ? | 27242034* |
| phoQ | phoQ | algR | algR | - | 27242034*, 18974177*, 22587778 |
| phoQ | phoQ | arnB | arnB | - | 27242034*, 18974177*, 22587778 |
| phoQ | phoQ | pmrA | pmrA | - | 27242034*, 18974177*, 22587778 |

| Regulatory gene | Ortholog of the regulatory gene | Target gene | Ortholog of the target gene | Mode of regulation | Reference (PubMed) |
|---|---|---|---|---|---|
| pilR | pilR | pilA | pilA | - | 27242034* |
| pilR | pilR | pilR | pilR | + | 27242034* |
| pilR | pilR | pilS | pilS | + | 27242034* |
| pmrA | pmrA | cysT | cysT | + | 27242034*, 18974177*, 22587778 |
| pmrA | pmrA | dnr | dnr | - | 27242034*, 18974177*, 22587778 |
| pmrA | pmrA | PA4851_24675 | feoA | + | 27242034*, 18974177*, 22587778 |
| pmrA | pmrA | PA4851_24670 | feoB | + | 27242034*, 18974177*, 22587778 |
| pmrA | pmrA | metK | metK | + | 27242034*, 18974177*, 22587778 |
| pmrA | pmrA | mexG | mexG | - | 27242034*, 18974177*, 22587778 |
| pmrA | pmrA | mexH | mexH | - | 27242034*, 18974177*, 22587778 |
| pmrA | pmrA | mexI | mexI | - | 27242034*, 18974177*, 22587778 |
| pmrA | pmrA | mgtA | mgtA | + | 27242034*, 18974177*, 22587778 |
| pmrA | pmrA | mgtE | mgtE | + | 27242034*, 18974177*, 22587778 |
| pmrA | pmrA | PA4851_01035 | PA0201 | + | 27242034*, 18974177*, 22587778 |
| pmrA | pmrA | PA4851_14850 | PA2359 | + | 27242034*, 18974177*, 22587778 |
| pmrA | pmrA | PA4851_07500 | PA3446 | + | 27242034*, 18974177*, 22587778 |
| pmrA | pmrA | PA4851_07260 | PA3515 | - | 27242034*, 18974177*, 22587778 |
| pmrA | pmrA | PA4851_07255 | PA3516 | - | 27242034*, 18974177*, 22587778 |
| pmrA | pmrA | PA4851_07250 | PA3517 | - | 27242034*, 18974177*, 22587778 |
| pmrA | pmrA | PA4851_07245 | PA3518 | - | 27242034*, 18974177*, 22587778 |
| pmrA | pmrA | PA4851_27460 | PA4773 | + | 27242034*, 18974177*, 22587778 |
| pmrA | pmrA | PA4851_27465 | PA4774 | + | 27242034*, 18974177*, 22587778 |
| pmrA | pmrA | PA4851_27470 | PA4775 | + | 27242034*, 18974177*, 22587778 |
| pmrA | pmrA | PA4851_27500 | PA4781 | + | 27242034*, 18974177*, 22587778 |
| pmrA | pmrA | PA4851_27505 | PA4782 | + | 27242034*, 18974177*, 22587778 |
| pmrA | pmrA | PA4851_27705 | PA4822 | + | 27242034*, 18974177*, 22587778 |
| pmrA | pmrA | PA4851_27710 | PA4823 | + | 27242034*, 18974177*, 22587778 |
| pmrA | pmrA | PA4851_27715 | PA4824 | + | 27242034*, 18974177*, 22587778 |
| pmrA | pmrA | PA4851_27725 | PA4826 | + | 27242034*, 18974177*, 22587778 |
| pmrA | pmrA | pcoA | pcoA | - | 27242034*, 18974177*, 22587778 |
| pmrA | pmrA | pcoB | pcoB | - | 27242034*, 18974177*, 22587778 |
| pmrA | pmrA | pmrA | pmrA | + | 27242034*, 18974177*, 22587778 |
| pmrA | pmrA | pmrB | pmrB | + | 27242034*, 18974177*, 22587778 |
| pmrA | pmrA | PA4851_24590 | PA0202 | + | 27242034*, 18974177*, 22587778 |
| pmrA | pmrA | PA4851_19170 | PA1559 | + | 27242034*, 18974177*, 22587778 |
| pmrA | pmrA | cueR | cueR | + | 27242034*, 18974177*, 22587778 |
| pmrA | pmrA | arnE | arnE | ? | 27242034* |
| pmrA | pmrA | arnD | arnD | ? | 27242034* |
| pmrA | pmrA | arnT | arnT | ? | 27242034* |
| pmrA | pmrA | arnC | arnC | ? | 27242034* |
| pmrA | pmrA | arnB | arnB | ? | 27242034* |
| pmrA | pmrA | arnA | arnA | ? | 27242034* |
| pmrA | pmrA | arnF | arnF | ? | 27242034* |
| pmrA | pmrA | PA4851_07040 | PA3559 | ? | 27242034* |
| pprB | pprB | rpoS | rpoS | + | 27242034*, 18974177*, 22587778 |

| Regulatory gene | Ortholog of the regulatory gene | Target gene | Ortholog of the target gene | Mode of regulation | Reference (PubMed) |
|---|---|---|---|---|---|
| pprB | pprB | PA4851_12250 | PA2591 | + | 27242034*, 18974177*, 22587778 |
| pprB | pprB | rsaL | rsaL | ? | 27242034* |
| pprB | pprB | tadZ | tadZ | ? | 27242034* |
| pprB | pprB | flp | flp | ? | 27242034* |
| pprB | pprB | rcpC | rcpC | ? | 27242034* |
| pprB | pprB | fppA | fppA | ? | 27242034* |
| pprB | pprB | pprB | pprB | ? | 27242034* |
| pprB | pprB | rcpA | rcpA | ? | 27242034* |
| pprB | pprB | lasB | lasB | ? | 27242034* |
| pprB | pprB | pprA | pprA | ? | 27242034* |
| pprB | pprB | PA4851_24350 | PA4294 | ? | 27242034* |
| pprB | pprB | lasI | lasI | ? | 27242034* |
| pprB | pprB | PA4851_24370 | PA4298 | ? | 27242034* |
| pprB | pprB | tadA | tadA | ? | 27242034* |
| pprB | pprB | tadG | tadG | ? | 27242034* |
| pprB | pprB | tadC | tadC | ? | 27242034* |
| pprB | pprB | tadB | tadB | ? | 27242034* |
| ppyR | ppyR | lasB | lasB | - | 27242034*, 18974177*, 22587778 |
| psrA | psrA | etfA | etfA | - | 27242034*, 18974177*, 22587778 |
| psrA | psrA | etfB | etfB | - | 27242034*, 18974177*, 22587778 |
| psrA | psrA | exoS | exoS | + | 27242034*, 18974177*, 22587778 |
| psrA | psrA | exsA | exsA | + | 27242034*, 18974177*, 22587778 |
| psrA | psrA | exsB | exsB | + | 27242034*, 18974177*, 22587778 |
| psrA | psrA | exsC | exsC | + | 27242034*, 18974177*, 22587778 |
| psrA | psrA | exsE | exsE | + | 27242034*, 18974177*, 22587778 |
| psrA | psrA | PA4851_18280 | fadB | + | 27242034*, 18974177*, 22587778 |
| psrA | psrA | mmsR | mmsR | + | 27242034*, 18974177*, 22587778 |
| psrA | psrA | PA4851_02535 | PA0506 | - | 27242034*, 18974177*, 22587778 |
| psrA | psrA | PA4851_10030 | PA2953 | - | 27242034*, 18974177*, 22587778 |
| psrA | psrA | PA4851_06860 | PA3595 | ? | 27242034*, 18974177*, 22587778 |
| psrA | psrA | psrA | psrA | - | 27242034*, 18974177*, 22587778 |
| psrA | psrA | rpoS | rpoS | + | 27242034*, 18974177*, 22587778 |
| psrA | psrA | PA4851_02540 | PA0507 | - | 27242034*, 18974177*, 22587778 |
| psrA | psrA | PA4851_02545 | PA0508 | - | 27242034*, 18974177*, 22587778 |
| psrA | psrA | PA4851_17780 | PA1831 | - | 27242034*, 18974177*, 22587778 |
| psrA | psrA | PA4851_17785 | PA1830 | - | 27242034*, 18974177*, 22587778 |
| psrA | psrA | faoA | faoA | - | 27242034*, 18974177*, 22587778 |
| psrA | psrA | algQ | algQ | - | 27242034*, 18974177*, 22587778 |
| ptxR | ptxR | pqsA | pqsA | - | 27242034*, 18974177*, 22587778 |
| ptxR | ptxR | pqsB | pqsB | - | 27242034*, 18974177*, 22587778 |
| ptxR | ptxR | pqsC | pqsC | - | 27242034*, 18974177*, 22587778 |
| ptxR | ptxR | pqsD | pqsD | - | 27242034*, 18974177*, 22587778 |
| ptxR | ptxR | ptxS | ptxS | + | 27242034*, 18974177*, 22587778 |
| ptxR | ptxR | pvcA | pvcA | + | 27242034*, 18974177*, 22587778 |
| ptxR | ptxR | pvcB | pvcB | + | 27242034*, 18974177*, 22587778 |

| Regulatory gene | Ortholog of the regulatory gene | Target gene | Ortholog of the target gene | Mode of regulation | Reference (PubMed) |
|---|---|---|---|---|---|
| ptxR | ptxR | pvcC | pvcC | + | 27242034*, 18974177*, 22587778 |
| ptxR | ptxR | pvcD | pvcD | + | 27242034*, 18974177*, 22587778 |
| ptxR | ptxR | rhlI | rhlI | - | 27242034*, 18974177*, 22587778 |
| ptxR | ptxR | toxA | toxA | + | 27242034*, 18974177*, 22587778 |
| ptxS | ptxS | PA4851_15325 | PA2263 | + | 27242034*, 18974177*, 22587778 |
| ptxS | ptxS | PA4851_15335 | PA2261 | ? | 27242034*, 18974177*, 22587778 |
| ptxS | ptxS | ptxR | ptxR | - | 27242034*, 18974177*, 22587778 |
| ptxS | ptxS | ptxS | ptxS | - | 27242034*, 18974177*, 22587778 |
| ptxS | ptxS | PA4851_15320 | PA2264 | - | 27242034*, 18974177*, 22587778 |
| ptxS | ptxS | PA4851_15315 | PA2265 | - | 27242034*, 18974177*, 22587778 |
| ptxS | ptxS | PA4851_15310 | PA2266 | - | 27242034*, 18974177*, 22587778 |
| ptxS | ptxS | PA4851_15330 | PA2262 | + | 27242034*, 18974177*, 22587778 |
| ptxS | ptxS | PA4851_15340 | PA2260 | + | 27242034*, 18974177*, 22587778 |
| pvdS | pvdS | pvdS | pvdS | ? | 27242034*, 18974177*, 22587778 |
| pvdS | pvdS | cat | cat | + | 27242034*, 18974177*, 22587778 |
| pvdS | pvdS | grx | grx | + | 27242034*, 18974177*, 22587778 |
| pvdS | pvdS | imm2 | imm2 | ? | 27242034*, 18974177*, 22587778 |
| pvdS | pvdS | PA4851_14690 | PA2390 | + | 27242034*, 18974177*, 22587778 |
| pvdS | pvdS | PA4851_14685 | opmQ | + | 27242034*, 18974177*, 22587778 |
| pvdS | pvdS | PA4851_01740 | PA0346 | + | 27242034*, 18974177*, 22587778 |
| pvdS | pvdS | PA4851_23505 | PA0818 | + | 27242034*, 18974177*, 22587778 |
| pvdS | pvdS | pvdR | PA2389 | + | 27242034*, 18974177*, 22587778 |
| pvdS | pvdS | PA4851_14675 | PA2393 | + | 27242034*, 18974177*, 22587778 |
| pvdS | pvdS | PA4851_14640 | PA2402 | + | 27242034*, 18974177*, 22587778 |
| pvdS | pvdS | PA4851_14590 | PA2411 | + | 27242034*, 18974177*, 22587778 |
| pvdS | pvdS | PA4851_14585 | PA2412 | + | 27242034*, 18974177*, 22587778 |
| pvdS | pvdS | PA4851_14510 | PA2427 | + | 27242034*, 18974177*, 22587778 |
| pvdS | pvdS | PA4851_13670 | PA2531 | + | 27242034*, 18974177*, 22587778 |
| pvdS | pvdS | PA4851_05855 | PA3794 | ? | 27242034*, 18974177*, 22587778 |
| pvdS | pvdS | PA4851_24830 | PA4390 | ? | 27242034*, 18974177*, 22587778 |
| pvdS | pvdS | PA4851_27760 | PA4833 | + | 27242034*, 18974177*, 22587778 |
| pvdS | pvdS | PA4851_29245 | PA5130 | + | 27242034*, 18974177*, 22587778 |
| pvdS | pvdS | PA4851_29395 | PA5150 | + | 27242034*, 18974177*, 22587778 |
| pvdS | pvdS | PA4851_29610 | PA5190 | ? | 27242034*, 18974177*, 22587778 |
| pvdS | pvdS | piv | prpL | + | 27242034*, 18974177*, 22587778 |
| pvdS | pvdS | ptxR | ptxR | + | 27242034*, 18974177*, 22587778 |
| pvdS | pvdS | pvcA | pvcA | + | 27242034*, 18974177*, 22587778 |
| pvdS | pvdS | pvcB | pvcB | + | 27242034*, 18974177*, 22587778 |
| pvdS | pvdS | pvcC | pvcC | + | 27242034*, 18974177*, 22587778 |
| pvdS | pvdS | pvcD | pvcD | + | 27242034*, 18974177*, 22587778 |
| pvdS | pvdS | pvdA | pvdA | + | 27242034*, 18974177*, 22587778 |
| pvdS | pvdS | PA4851_14655 | pvdE | + | 27242034*, 18974177*, 22587778 |
| pvdS | pvdS | pvdF | pvdF | + | 27242034*, 18974177*, 22587778 |
| pvdS | pvdS | pvdG | pvdG | + | 27242034*, 18974177*, 22587778 |
| pvdS | pvdS | pvdH | pvdH | + | 27242034*, 18974177*, 22587778 |

| Regulatory gene | Ortholog of the regulatory gene | Target gene | Ortholog of the target gene | Mode of regulation | Reference (PubMed) |
|---|---|---|---|---|---|
| pvdS | pvdS | PA4851_14670 | pvdN | + | 27242034*, 18974177*, 22587778 |
| pvdS | pvdS | pys2 | pys2 | ? | 27242034*, 18974177*, 22587778 |
| pvdS | pvdS | toxR | toxR | + | 27242034*, 18974177*, 22587778 |
| pvdS | pvdS | toxA | toxA | + | 27242034*, 18974177*, 22587778 |
| pvdS | pvdS | PA4851_14390 | PA2452 | + | 27242034*, 18974177*, 22587778 |
| pvdS | pvdS | pvdL | pvdL | + | 27242034*, 18974177*, 22587778 |
| qscR | qscR | rsaL | rsaL | - | 27242034*, 18974177*, 22587778 |
| qscR | qscR | PA4851_17450 | PA1897 | + | 27242034*, 18974177*, 22587778 |
| qscR | qscR | phzA2 | phzA2 | ? | 27242034* |
| qscR | qscR | rubA1 | rubA1 | ? | 27242034* |
| qscR | qscR | lasI | lasI | ? | 27242034* |
| qscR | qscR | phzF2 | phzF2 | ? | 27242034* |
| qscR | qscR | phzB1 | phzB1 | ? | 27242034* |
| qscR | qscR | phzB2 | phzB2 | ? | 27242034* |
| qscR | qscR | hcnA | hcnA | ? | 27242034* |
| qscR | qscR | hcnC | hcnC | ? | 27242034* |
| qscR | qscR | hcnB | hcnB | ? | 27242034* |
| qscR | qscR | phzD1 | phzD1 | ? | 27242034* |
| qscR | qscR | rhlI | rhlI | ? | 27242034* |
| qscR | qscR | phzA1 | phzA1 | ? | 27242034* |
| qscR | qscR | lasB | lasB | ? | 27242034* |
| qscR | qscR | PA4851_17455 | PA1896 | ? | 27242034* |
| qscR | qscR | PA4851_17460 | PA1895 | ? | 27242034* |
| qscR | qscR | PA4851_17465 | PA1894 | ? | 27242034* |
| qscR | qscR | PA4851_17470 | PA1893 | ? | 27242034* |
| qscR | qscR | PA4851_17475 | PA1892 | ? | 27242034* |
| qscR | qscR | PA4851_17480 | PA1891 | ? | 27242034* |
| qscR | qscR | phzG2 | phzG2 | ? | 27242034* |
| qscR | qscR | phzG1 | phzG1 | ? | 27242034* |
| qscR | qscR | phzC1 | phzC1 | ? | 27242034* |
| qscR | qscR | phzE1 | phzE1 | ? | 27242034* |
| RbsR | RbsR | rbsB | rbsB | - | 27242034* |
| RbsR | RbsR | rbsA | rbsA | - | 27242034* |
| RbsR | RbsR | rbsC | rbsC | - | 27242034* |
| RbsR | RbsR | rbsR | rbsR | - | 27242034* |
| RbsR | RbsR | rbsK | rbsK | - | 27242034* |
| recA | recA | recA | recA | ? | 27242034* |
| rhlR | rhlR | acpP | acpP | + | 27242034*, 18974177*, 22587778 |
| rhlR | rhlR | PA4851_17590 | PA1869 | + | 27242034*, 18974177*, 22587778 |
| rhlR | rhlR | exoS | exoS | - | 27242034*, 18974177*, 22587778 |
| rhlR | rhlR | hcnA | hcnA | + | 27242034*, 18974177*, 22587778 |
| rhlR | rhlR | hcnB | hcnB | + | 27242034*, 18974177*, 22587778 |
| rhlR | rhlR | hcnC | hcnC | + | 27242034*, 18974177*, 22587778 |
| rhlR | rhlR | lecA | lecA | + | 27242034*, 18974177*, 22587778 |
| rhlR | rhlR | migA | migA | + | 27242034*, 18974177*, 22587778 |

| Regulatory gene | Ortholog of the regulatory gene | Target gene | Ortholog of the target gene | Mode of regulation | Reference (PubMed) |
|---|---|---|---|---|---|
| rhlR | rhlR | mvfR | mvfR | - | 27242034*, 18974177*, 22587778 |
| rhlR | rhlR | phzA1 | phzA1 | + | 27242034*, 18974177*, 22587778 |
| rhlR | rhlR | phzB1 | phzB1 | + | 27242034*, 18974177*, 22587778 |
| rhlR | rhlR | phzC1 | phzC1 | + | 27242034*, 18974177*, 22587778 |
| rhlR | rhlR | phzD1 | phzD1 | + | 27242034*, 18974177*, 22587778 |
| rhlR | rhlR | phzE1 | phzE1 | + | 27242034*, 18974177*, 22587778 |
| rhlR | rhlR | phzF1 | phzF1 | + | 27242034*, 18974177*, 22587778 |
| rhlR | rhlR | phzG1 | phzG1 | + | 27242034*, 18974177*, 22587778 |
| rhlR | rhlR | pqsA | pqsA | - | 27242034*, 18974177*, 22587778 |
| rhlR | rhlR | pqsB | pqsB | - | 27242034*, 18974177*, 22587778 |
| rhlR | rhlR | pqsC | pqsC | - | 27242034*, 18974177*, 22587778 |
| rhlR | rhlR | pqsD | pqsD | - | 27242034*, 18974177*, 22587778 |
| rhlR | rhlR | pqsE | pqsE | - | 27242034*, 18974177*, 22587778 |
| rhlR | rhlR | rhlA | rhlA | + | 27242034*, 18974177*, 22587778 |
| rhlR | rhlR | rhlB | rhlB | - | 27242034*, 18974177*, 22587778 |
| rhlR | rhlR | rhlG | rhlG | ? | 27242034*, 18974177*, 22587778 |
| rhlR | rhlR | rhlI | rhlI | + | 27242034*, 18974177*, 22587778 |
| rhlR | rhlR | rhlR | rhlR | - | 27242034*, 18974177*, 22587778 |
| rhlR | rhlR | rpoS | rpoS | - | 27242034*, 18974177*, 22587778 |
| rhlR | rhlR | xcpW | xcpW | ? | 27242034* |
| rhlR | rhlR | xcpV | xcpV | ? | 27242034* |
| rhlR | rhlR | xcpU | xcpU | ? | 27242034* |
| rhlR | rhlR | xcpT | xcpT | ? | 27242034* |
| rhlR | rhlR | xcpS | xcpS | ? | 27242034* |
| rhlR | rhlR | xcpR | xcpR | ? | 27242034* |
| rhlR | rhlR | xcpQ | xcpQ | ? | 27242034* |
| rhlR | rhlR | phzF2 | phzF2 | ? | 27242034* |
| rhlR | rhlR | xcpZ | xcpZ | ? | 27242034* |
| rhlR | rhlR | xcpX | xcpX | ? | 27242034* |
| rhlR | rhlR | chiC | chiC | ? | 27242034* |
| rhlR | rhlR | PA4851_00920 | PA0179 | ? | 27242034* |
| rhlR | rhlR | lasA | lasA | ? | 27242034* |
| rhlR | rhlR | lasB | lasB | ? | 27242034* |
| rhlR | rhlR | lasI | lasI | ? | 27242034* |
| rhlR | rhlR | phzG2 | phzG2 | ? | 27242034* |
| rhlR | rhlR | xcpP | xcpP | ? | 27242034* |
| rocA1 | rocA1 | cupB1 | cupB1 | + | 27242034*, 18974177*, 22587778 |
| rocA1 | rocA1 | cupC1 | cupC1 | - | 27242034*, 18974177*, 22587778 |
| rocA1 | rocA1 | cupB4 | cupB4 | ? | 27242034* |
| rocA1 | rocA1 | rocR | rocR | ? | 27242034* |
| rocA1 | rocA1 | cupB2 | cupB2 | ? | 27242034* |
| rocA1 | rocA1 | rocS1 | rocS1 | ? | 27242034* |
| rocA1 | rocA1 | cupB6 | cupB6 | ? | 27242034* |
| rocA1 | rocA1 | cupB5 | cupB5 | ? | 27242034* |
| rocA1 | rocA1 | cupC3 | cupC3 | ? | 27242034* |

| Regulatory gene | Ortholog of the regulatory gene | Target gene | Ortholog of the target gene | Mode of regulation | Reference (PubMed) |
|---|---|---|---|---|---|
| rocA1 | rocA1 | cupB3 | cupB3 | ? | 27242034* |
| rocA1 | rocA1 | cupC2 | cupC2 | ? | 27242034* |
| rocA1 | rocA1 | rocA1 | rocA1 | ? | 27242034* |
| roxR | roxR | ccoN1 | ccoN1 | + | 27242034*, 18974177*, 22587778 |
| roxR | roxR | ccoN2 | ccoN2 | + | 27242034*, 18974177*, 22587778 |
| roxR | roxR | ccoO1 | ccoO1 | + | 27242034*, 18974177*, 22587778 |
| roxR | roxR | ccoO2 | ccoO2 | + | 27242034*, 18974177*, 22587778 |
| roxR | roxR | ccoP1 | ccoP1 | + | 27242034*, 18974177*, 22587778 |
| roxR | roxR | ccoP2 | ccoP2 | + | 27242034*, 18974177*, 22587778 |
| roxR | roxR | ccoQ1 | ccoQ1 | + | 27242034*, 18974177*, 22587778 |
| roxR | roxR | ccoQ2 | ccoQ2 | + | 27242034*, 18974177*, 22587778 |
| roxR | roxR | cioA | cioA | + | 27242034*, 18974177*, 22587778 |
| roxR | roxR | cioB | cioB | + | 27242034*, 18974177*, 22587778 |
| roxR | roxR | coxA | coxA | - | 27242034*, 18974177*, 22587778 |
| roxR | roxR | coxB | coxB | - | 27242034*, 18974177*, 22587778 |
| roxR | roxR | cyoA | cyoA | + | 27242034*, 18974177*, 22587778 |
| roxR | roxR | cyoB | cyoB | + | 27242034*, 18974177*, 22587778 |
| roxR | roxR | cyoC | cyoC | + | 27242034*, 18974177*, 22587778 |
| roxR | roxR | cyoD | cyoD | + | 27242034*, 18974177*, 22587778 |
| roxR | roxR | cyoE | cyoE | + | 27242034*, 18974177*, 22587778 |
| roxR | roxR | roxR | roxR | + | 27242034*, 18974177*, 22587778 |
| roxS | roxS | ccoN1 | ccoN1 | + | 27242034*, 18974177*, 22587778 |
| roxS | roxS | ccoN2 | ccoN2 | + | 27242034*, 18974177*, 22587778 |
| roxS | roxS | ccoO1 | ccoO1 | + | 27242034*, 18974177*, 22587778 |
| roxS | roxS | ccoO2 | ccoO2 | + | 27242034*, 18974177*, 22587778 |
| roxS | roxS | ccoP1 | ccoP1 | + | 27242034*, 18974177*, 22587778 |
| roxS | roxS | ccoP2 | ccoP2 | + | 27242034*, 18974177*, 22587778 |
| roxS | roxS | ccoQ1 | ccoQ1 | + | 27242034*, 18974177*, 22587778 |
| roxS | roxS | ccoQ2 | ccoQ2 | + | 27242034*, 18974177*, 22587778 |
| roxS | roxS | cioA | cioA | + | 27242034*, 18974177*, 22587778 |
| roxS | roxS | cioB | cioB | + | 27242034*, 18974177*, 22587778 |
| roxS | roxS | coxA | coxA | - | 27242034*, 18974177*, 22587778 |
| roxS | roxS | coxB | coxB | - | 27242034*, 18974177*, 22587778 |
| roxS | roxS | cyoA | cyoA | + | 27242034*, 18974177*, 22587778 |
| roxS | roxS | cyoB | cyoB | + | 27242034*, 18974177*, 22587778 |
| roxS | roxS | cyoC | cyoC | + | 27242034*, 18974177*, 22587778 |
| roxS | roxS | cyoD | cyoD | + | 27242034*, 18974177*, 22587778 |
| roxS | roxS | cyoE | cyoE | + | 27242034*, 18974177*, 22587778 |
| rpoD | rpoD | aotJ | aotJ | + | 27242034*, 18974177*, 22587778 |
| rpoD | rpoD | fleQ | fleQ | + | 27242034*, 18974177*, 22587778 |
| rpoD | rpoD | ptxR | ptxR | + | 27242034*, 18974177*, 22587778 |
| rpoD | rpoD | rpoS | rpoS | ? | 27242034*, 18974177*, 22587778 |
| rpoD | rpoD | rpoD | rpoD | ? | 27242034*, 18974177*, 22587778 |
| rpoN | rpoN | alg44 | alg44 | + | 27242034*, 18974177*, 22587778 |
| rpoN | rpoN | alg8 | alg8 | + | 27242034*, 18974177*, 22587778 |

| Regulatory gene | Ortholog of the regulatory gene | Target gene | Ortholog of the target gene | Mode of regulation | Reference (PubMed) |
|---|---|---|---|---|---|
| rpoN | rpoN | algA | algA | + | 27242034*, 18974177*, 22587778 |
| rpoN | rpoN | algB | algB | ? | 27242034*, 18974177*, 22587778 |
| rpoN | rpoN | algD | algD | d | 27242034*, 18974177*, 22587778 |
| rpoN | rpoN | algE | algE | + | 27242034*, 18974177*, 22587778 |
| rpoN | rpoN | algF | algF | + | 27242034*, 18974177*, 22587778 |
| rpoN | rpoN | algG | algG | + | 27242034*, 18974177*, 22587778 |
| rpoN | rpoN | algI | algI | + | 27242034*, 18974177*, 22587778 |
| rpoN | rpoN | algJ | algJ | + | 27242034*, 18974177*, 22587778 |
| rpoN | rpoN | algK | algK | + | 27242034*, 18974177*, 22587778 |
| rpoN | rpoN | algL | algL | + | 27242034*, 18974177*, 22587778 |
| rpoN | rpoN | algX | algX | + | 27242034*, 18974177*, 22587778 |
| rpoN | rpoN | fhp | fhp | + | 27242034*, 18974177*, 22587778 |
| rpoN | rpoN | PA4851_11865 | fhpR | + | 27242034*, 18974177*, 22587778 |
| rpoN | rpoN | fleN | fleN | + | 27242034*, 18974177*, 22587778 |
| rpoN | rpoN | fleR | fleR | + | 27242034*, 18974177*, 22587778 |
| rpoN | rpoN | fleS | fleS | + | 27242034*, 18974177*, 22587778 |
| rpoN | rpoN | flgB | flgB | + | 27242034*, 18974177*, 22587778 |
| rpoN | rpoN | flgC | flgC | + | 27242034*, 18974177*, 22587778 |
| rpoN | rpoN | flgD | flgD | + | 27242034*, 18974177*, 22587778 |
| rpoN | rpoN | flgE | flgE | + | 27242034*, 18974177*, 22587778 |
| rpoN | rpoN | flgF | flgF | + | 27242034*, 18974177*, 22587778 |
| rpoN | rpoN | flgG | flgG | + | 27242034*, 18974177*, 22587778 |
| rpoN | rpoN | flgH | flgH | + | 27242034*, 18974177*, 22587778 |
| rpoN | rpoN | flgI | flgI | + | 27242034*, 18974177*, 22587778 |
| rpoN | rpoN | flgJ | flgJ | + | 27242034*, 18974177*, 22587778 |
| rpoN | rpoN | flgK | flgK | + | 27242034*, 18974177*, 22587778 |
| rpoN | rpoN | flgL | flgL | + | 27242034*, 18974177*, 22587778 |
| rpoN | rpoN | flhA | flhA | + | 27242034*, 18974177*, 22587778 |
| rpoN | rpoN | flhB | flhB | + | 27242034*, 18974177*, 22587778 |
| rpoN | rpoN | flhF | flhF | + | 27242034*, 18974177*, 22587778 |
| rpoN | rpoN | fliD | fliD | + | 27242034*, 18974177*, 22587778 |
| rpoN | rpoN | fliE | fliE | + | 27242034*, 18974177*, 22587778 |
| rpoN | rpoN | fliF | fliF | + | 27242034*, 18974177*, 22587778 |
| rpoN | rpoN | fliG | fliG | + | 27242034*, 18974177*, 22587778 |
| rpoN | rpoN | PA4851_21790 | fliH | + | 27242034*, 18974177*, 22587778 |
| rpoN | rpoN | fliI | fliI | + | 27242034*, 18974177*, 22587778 |
| rpoN | rpoN | fliJ | fliJ | + | 27242034*, 18974177*, 22587778 |
| rpoN | rpoN | PA4851_19770 | PA1442 | + | 27242034*, 18974177*, 22587778 |
| rpoN | rpoN | fliM | fliM | + | 27242034*, 18974177*, 22587778 |
| rpoN | rpoN | fliN | fliN | + | 27242034*, 18974177*, 22587778 |
| rpoN | rpoN | fliO | fliO | + | 27242034*, 18974177*, 22587778 |
| rpoN | rpoN | fliP | fliP | + | 27242034*, 18974177*, 22587778 |
| rpoN | rpoN | fliQ | fliQ | + | 27242034*, 18974177*, 22587778 |
| rpoN | rpoN | fliR | fliR | + | 27242034*, 18974177*, 22587778 |
| rpoN | rpoN | PA4851_21830 | fliS | + | 27242034*, 18974177*, 22587778 |

| Regulatory gene | Ortholog of the regulatory gene | Target gene | Ortholog of the target gene | Mode of regulation | Reference (PubMed) |
|---|---|---|---|---|---|
| rpoN | rpoN | oprE | oprE | + | 27242034*, 18974177*, 22587778 |
| rpoN | rpoN | wbpI | orfK | + | 27242034*, 18974177*, 22587778 |
| rpoN | rpoN | wbpK | orfM | + | 27242034*, 18974177*, 22587778 |
| rpoN | rpoN | wbpL | orfN | + | 27242034*, 18974177*, 22587778 |
| rpoN | rpoN | PA4851_22010 | phaG | + | 27242034*, 18974177*, 22587778 |
| rpoN | rpoN | proC | proC | + | 27242034*, 18974177*, 22587778 |
| rpoN | rpoN | rhlI | rhlI | - | 27242034*, 18974177*, 22587778 |
| rpoN | rpoN | rhlR | rhlR | + | 27242034*, 18974177*, 22587778 |
| rpoN | rpoN | PA4851_30400 | sadB | ? | 27242034*, 18974177*, 22587778 |
| rpoN | rpoN | glnA | glnA | ? | 27242034*, 18974177*, 22587778 |
| rpoN | rpoN | gacA | gacA | ? | 27242034* |
| rpoN | rpoN | rhlB | rhlB | ? | 27242034* |
| rpoN | rpoN | rhlA | rhlA | ? | 27242034* |
| rpoN | rpoN | ureB | ureB | ? | 27242034* |
| rpoN | rpoN | ureA | ureA | ? | 27242034* |
| rpoN | rpoN | ureC | ureC | ? | 27242034* |
| rpoS | rpoS | azu | azu | + | 27242034*, 18974177*, 22587778 |
| rpoS | rpoS | bphO | bphO | + | 27242034*, 18974177*, 22587778 |
| rpoS | rpoS | bphP | bphP | + | 27242034*, 18974177*, 22587778 |
| rpoS | rpoS | cioA | cioA | + | 27242034*, 18974177*, 22587778 |
| rpoS | rpoS | cioB | cioB | + | 27242034*, 18974177*, 22587778 |
| rpoS | rpoS | coIII | coIII | + | 27242034*, 18974177*, 22587778 |
| rpoS | rpoS | coxA | coxA | + | 27242034*, 18974177*, 22587778 |
| rpoS | rpoS | coxB | coxB | + | 27242034*, 18974177*, 22587778 |
| rpoS | rpoS | exsA | exsA | + | 27242034*, 18974177*, 22587778 |
| rpoS | rpoS | exsB | exsB | + | 27242034*, 18974177*, 22587778 |
| rpoS | rpoS | exsC | exsC | + | 27242034*, 18974177*, 22587778 |
| rpoS | rpoS | exsE | exsE | + | 27242034*, 18974177*, 22587778 |
| rpoS | rpoS | rhlI | rhlI | - | 27242034*, 18974177*, 22587778 |
| rpoS | rpoS | hcnA | hcnA | ? | 27242034*, 18974177*, 22587778 |
| rpoS | rpoS | phzC1 | phzC1 | ? | 27242034*, 18974177*, 22587778 |
| rpoS | rpoS | phzF1 | phzF1 | ? | 27242034*, 18974177*, 22587778 |
| rpoS | rpoS | lasB | lasB | ? | 27242034*, 18974177*, 22587778 |
| rpoS | rpoS | lecA | lecA | ? | 27242034* |
| rpoS | rpoS | aer2 | aer2 | ? | 27242034* |
| rpoS | rpoS | phzF2 | phzF2 | ? | 27242034* |
| rpoS | rpoS | phzD1 | phzD1 | ? | 27242034* |
| rpoS | rpoS | phzB1 | phzB1 | ? | 27242034* |
| rpoS | rpoS | phzG1 | phzG1 | ? | 27242034* |
| rpoS | rpoS | hcnC | hcnC | ? | 27242034* |
| rpoS | rpoS | hcnB | hcnB | ? | 27242034* |
| rpoS | rpoS | PA4851_17285 | PA1930 | ? | 27242034* |
| rpoS | rpoS | cttP | cttP | ? | 27242034* |
| rpoS | rpoS | PA4851_00920 | PA0179 | ? | 27242034* |
| rpoS | rpoS | PA4851_00915 | PA0178 | ? | 27242034* |

| Regulatory gene | Ortholog of the regulatory gene | Target gene | Ortholog of the target gene | Mode of regulation | Reference (PubMed) |
|---|---|---|---|---|---|
| rpoS | rpoS | PA4851_00910 | PA0177 | ? | 27242034* |
| rpoS | rpoS | phzA1 | phzA1 | ? | 27242034* |
| rpoS | rpoS | phzG2 | phzG2 | ? | 27242034* |
| rpoS | rpoS | lasA | lasA | ? | 27242034* |
| rpoS | rpoS | PA4851_13455 | PA2573 | ? | 27242034* |
| rpoS | rpoS | phzE1 | phzE1 | ? | 27242034* |
| rsaL | rsaL | phzA1 | phzA1 | - | 27242034*, 18974177*, 22587778 |
| rsaL | rsaL | lasI | lasI | - | 27242034*, 18974177*, 22587778 |
| rsaL | rsaL | lasB | lasB | ? | 27242034*, 18974177*, 22587778 |
| rsaL | rsaL | rsaL | rsaL | - | 27242034*, 18974177*, 22587778 |
| rsaL | rsaL | phzM | phzM | - | 27242034*, 18974177*, 22587778 |
| rsaL | rsaL | hcnA | hcnA | - | 27242034*, 18974177*, 22587778 |
| rsaL | rsaL | phzF1 | phzF1 | ? | 27242034* |
| rsaL | rsaL | phzF2 | phzF2 | ? | 27242034* |
| rsaL | rsaL | phzB1 | phzB1 | ? | 27242034* |
| rsaL | rsaL | phzC1 | phzC1 | ? | 27242034* |
| rsaL | rsaL | phzE1 | phzE1 | ? | 27242034* |
| rsaL | rsaL | phzD1 | phzD1 | ? | 27242034* |
| rsaL | rsaL | phzG1 | phzG1 | ? | 27242034* |
| rsaL | rsaL | lasI | lasI | ? | 27242034* |
| rsaL | rsaL | phzG2 | phzG2 | ? | 27242034* |
| rsaL | rsaL | hcnC | hcnC | ? | 27242034* |
| rsaL | rsaL | hcnB | hcnB | ? | 27242034* |
| soxR | soxR | PA4851_06240 | PA3718 | + | 27242034*, 18974177*, 22587778 |
| soxR | soxR | mexG | mexG | + | 27242034*, 18974177*, 22587778 |
| soxR | soxR | mexH | mexH | + | 27242034*, 18974177*, 22587778 |
| soxR | soxR | mexI | mexI | + | 27242034*, 18974177*, 22587778 |
| soxR | soxR | opmD | opmD | + | 27242034*, 18974177*, 22587778 |
| soxR | soxR | PA4851_15270 | PA2274 | + | 27242034*, 18974177*, 22587778 |
| soxR | soxR | soxR | soxR | + | 27242034*, 18974177*, 22587778 |
| toxR | toxR | oprL | oprL | + | 27242034*, 18974177*, 22587778 |
| toxR | toxR | PA4851_05610 | PA3842 | + | 27242034*, 18974177*, 22587778 |
| toxR | toxR | motD | motD | + | 27242034*, 18974177*, 22587778 |
| toxR | toxR | tolA | tolA | + | 27242034*, 18974177*, 22587778 |
| toxR | toxR | tolB | tolB | + | 27242034*, 18974177*, 22587778 |
| toxR | toxR | tolQ | tolQ | + | 27242034*, 18974177*, 22587778 |
| toxR | toxR | tolR | tolR | + | 27242034*, 18974177*, 22587778 |
| toxR | toxR | toxA | toxA | - | 27242034*, 18974177*, 22587778 |
| tpbA | tpbA | PA4851_04145 | PA4139 | - | 27242034*, 18974177*, 22587778 |
| tpbA | tpbA | PA4851_26630 | PA4624 | - | 27242034*, 18974177*, 22587778 |
| tpbA | tpbA | PA4851_26635 | PA4625 | - | 27242034*, 18974177*, 22587778 |
| tpbA | tpbA | pelA | pelA | - | 27242034*, 18974177*, 22587778 |
| tpbA | tpbA | pelB | pelB | - | 27242034*, 18974177*, 22587778 |
| tpbA | tpbA | pelC | pelC | - | 27242034*, 18974177*, 22587778 |
| tpbA | tpbA | pelD | pelD | - | 27242034*, 18974177*, 22587778 |

| Regulatory gene | Ortholog of the regulatory gene | Target gene | Ortholog of the target gene | Mode of regulation | Reference (PubMed) |
|---|---|---|---|---|---|
| tpbA | tpbA | pelE | pelE | - | 27242034*, 18974177*, 22587778 |
| tpbA | tpbA | pelF | pelF | - | 27242034*, 18974177*, 22587778 |
| tpbA | tpbA | pelG | pelG | - | 27242034*, 18974177*, 22587778 |
| tpbA | tpbA | tpbA | tpbA | - | 27242034*, 18974177*, 22587778 |
| tpbA | tpbA | tpbB | tpbB | - | 27242034*, 18974177*, 22587778 |
| trpI | trpI | trpA | trpA | + | 27242034* |
| trpI | trpI | trpB | trpB | + | 27242034* |
| trpI | trpI | trpI | trpI | - | 27242034* |
| vfr | vfr | cbpA | cbpA | + | 27242034*, 18974177*, 22587778 |
| vfr | vfr | fleQ | fleQ | - | 27242034*, 18974177*, 22587778 |
| vfr | vfr | lasR | lasR | + | 27242034*, 18974177*, 22587778 |
| vfr | vfr | ptxR | ptxR | + | 27242034*, 18974177*, 22587778 |
| vfr | vfr | toxR | toxR | + | 27242034*, 18974177*, 22587778 |
| vfr | vfr | rhlR | rhlR | d | 27242034*, 18974177*, 22587778 |
| vfr | vfr | toxA | toxA | + | 27242034*, 18974177*, 22587778 |
| vfr | vfr | vfr | vfr | + | 27242034*, 18974177*, 22587778 |
| vfr | vfr | pilP | pilP | ? | 27242034* |
| vfr | vfr | plcH | plcH | ? | 27242034* |
| vfr | vfr | PA4851_03305 | PA0653 | ? | 27242034* |
| vfr | vfr | plcN | plcN | ? | 27242034* |
| vfr | vfr | plcR | plcR | ? | 27242034* |
| vfr | vfr | pilM | pilM | ? | 27242034* |
| vfr | vfr | pilO | pilO | ? | 27242034* |
| vfr | vfr | pilN | pilN | ? | 27242034* |
| vfr | vfr | alg8 | alg8 | ? | 27242034* |
| vfr | vfr | algZ | algZ | ? | 27242034* |
| vfr | vfr | alg44 | alg44 | ? | 27242034* |
| vfr | vfr | algJ | algJ | ? | 27242034* |
| vfr | vfr | algK | algK | ? | 27242034* |
| vfr | vfr | algI | algI | ? | 27242034* |
| vfr | vfr | algL | algL | ? | 27242034* |
| vfr | vfr | lasI | lasI | ? | 27242034* |
| vfr | vfr | algA | algA | ? | 27242034* |
| vfr | vfr | algF | algF | ? | 27242034* |
| vfr | vfr | algG | algG | ? | 27242034* |
| vfr | vfr | algD | algD | ? | 27242034* |
| vfr | vfr | algE | algE | ? | 27242034* |
| vfr | vfr | exoT | exoT | ? | 27242034* |
| vfr | vfr | argH | argH | ? | 27242034* |
| vfr | vfr | pbpG | pbpG | ? | 27242034* |
| vfr | vfr | algX | algX | ? | 27242034* |
| vqsM | vqsM | pprB | pprB | + | 27242034*, 18974177*, 22587778 |
| vqsM | vqsM | rpoS | rpoS | + | 27242034*, 18974177*, 22587778 |
| vqsM | vqsM | PA4851_12250 | PA2591 | + | 27242034*, 18974177*, 22587778 |
| vreI | vreI | PA4851_24145 | PA0691 | + | 27242034* |

| Regulatory gene | Ortholog of the regulatory gene | Target gene | Ortholog of the target gene | Mode of regulation | Reference (PubMed) |
|---|---|---|---|---|---|
| vreI | vreI | PA4851_14345 | tpsB | + | 27242034* |

*Information retrieved from databases.*
