## Supplementary figures and images for "Gene regulatory network inference and analysis of multidrug-resistant *Pseudomonas aeruginosa*"

### Figure 2 (better resolution)

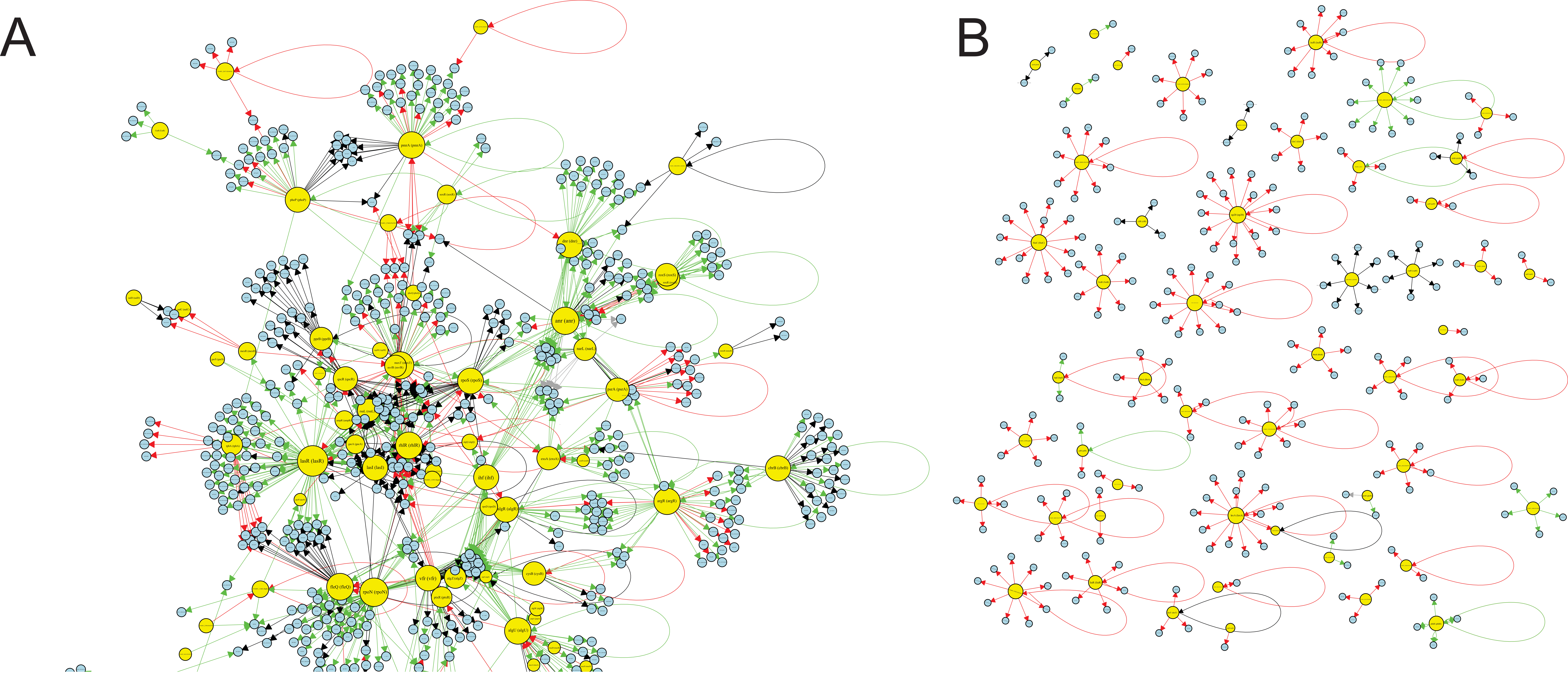

### Figure 4 (better resolution)

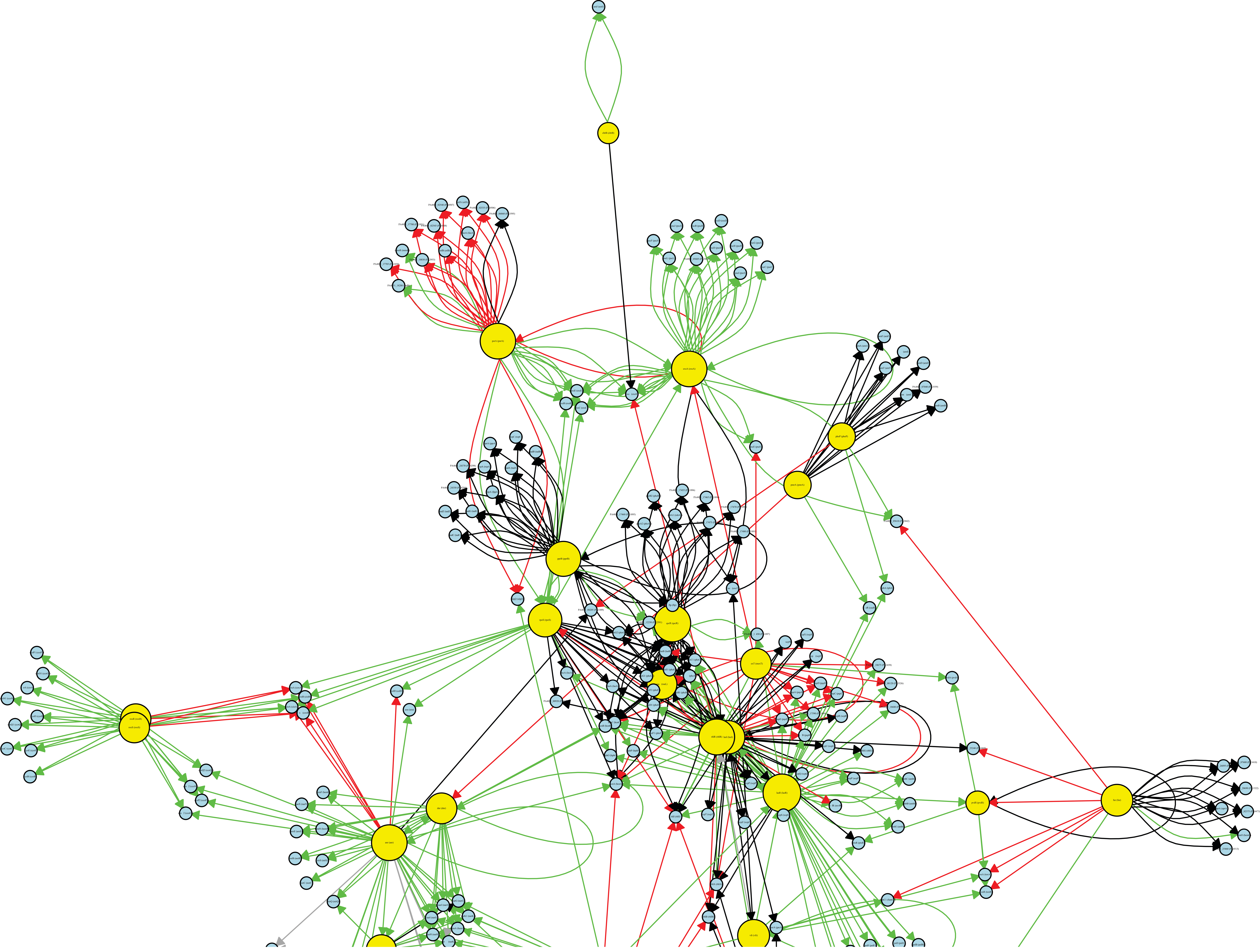
