## Supplementary material for "Gene regulatory network inference and analysis of multidrug-resistant *Pseudomonas aeruginosa*": BBH algorithm

```
Usage = """RBH BLASTOUTPUT1 BLASTOUTPUT2 RBH-list-outfile """

import sys, re

if len(sys.argv) < 3:
      print(Usage)

debug = 9

infl1 = sys.argv[1]
infl2 = sys.argv[2]
outfile = sys.argv[3]

#parse first BLAST results
FL1 = open(infl1, 'r')
D1 = {} #dictionary for BLAST file ONE
for Line in FL1:
      if ( Line[0] != '#' ):
            Line.strip()
            Elements = re.split('\t', Line)
            queryId = Elements[0]
            subjectId = Elements[1]
            if ( not ( queryId in D1.keys() ) ):
                  D1[queryId] = subjectId  #pick the first hit

if (debug): D1.keys()

#parse second BLAST results
FL2 = open(infl2, 'r')
D2 = {}
for Line in FL2:
      if ( Line[0] != '#' ):
            Line.strip()
            Elements = re.split('\t', Line)
            queryId = Elements[0]
            subjectId = Elements[1]
            if ( not ( queryId in D2.keys() ) ):
                  D2[queryId] = subjectId  #pick the first hit

if (debug): D2.keys()

#Now, pick the share pairs

SharedPairs={}
for id1 in D1.keys():
      value1 = D1[id1]
      if ( value1 in D2.keys() ):
            if ( id1 == D2[value1] ) : #a shared best reciprocal
pair
                  SharedPairs[value1] = id1

if (debug): SharedPairs

#outfl = open("_out.csv", "w")
outfl = open( outfile, 'w')

for k1 in SharedPairs.keys():
      line = k1 + '\t' + SharedPairs[k1] + '\n'
```

```
    outfl.write(line)

outfl.close()

print("Done. RBH from", sys.argv[1], "and", sys.argv[2], "are in",
sys.argv[3])
```
