## Supplementary material for "Gene regulatory network inference and analysis of multidrug-resistant *Pseudomonas aeruginosa*": GRN R code

```
# packages 1
library(dplyr)
library(tibble)
library(readr)
# packages 2
library(igraph)
library(scales)


dados <-
  read_csv2("GRN.csv")

dados


c1 <-
  dados$`Regulator (TF or sigma)` %>%
  strsplit(" ") %>%
  unlist()

c1.TF <- c1[gtools::odd(1:length(c1))]
c1.ortologo.TF <- c1[gtools::even(1:length(c1))]


dados$`Regulator (TF)` <- c1.TF
dados$`orthologs of TF` <- c1.ortologo.TF


nrow(dados)  ==
  sum(paste(dados$`Regulator (TF)`,dados$`orthologs of TF`) ==
dados$`Regulator (TF or sigma)`)


dados <-
  dados %>%
  select(`Regulator (TF)`,
         `Target gene`,
         `mode of regulation`,
         `orthologs of TF`,
         `Ortholog of the target gene`
  )


dados$`orthologs of TF` <- gsub("\\(|\\)", "",
                                dados$`orthologs of TF`)



rm(c1, c1.ortologo.TF,c1.TF)


auxTF <- dados[,c(1,4)] %>% setNames(c("gene_CCBH4851",
"orthologs"))
auxTarget <- dados[,c(2,5)] %>% setNames(c("gene_CCBH4851",
"orthologs"))

vert <-
  dplyr::union(auxTF,auxTarget) %>%
```

```
  filter(!is.na(gene_CCBH4851) )

vert$rotulo <- paste0(vert$gene_CCBH4851,
                      " (",
                      vert$orthologs,
                      ")")


arestas <-
  dados[,c(1,2,3)] %>%
  filter(!is.na(`Target gene`))


Rede <- graph_from_data_frame(d = arestas,
                              directed = TRUE,
                              vertices = vert
)



V(Rede)$color <- ifelse(V(Rede)$name %in% auxTF$gene_CCBH4851,
                        "yellow", "lightblue")


codificacao <- '"+" = "green" ; "-" = "red" ; "?" = "white" ; "d" =
"darkgrey"'
E(Rede)$color <- car::Recode(E(Rede)$`mode of regulation`,
                             codificacao)


V(Rede)$size <- 2+ log(1+degree(Rede, mode = "out"))

V(Rede)$name <- V(Rede)$rotulo


set.seed(1234)
#
plot(Rede,
     layout=layout_nicely,
     vertex.label.dist=0,
     vertex.label.color='black',
     vertex.label.font=0.05,
     vertex.label.cex=0.7,
     edge.arrow.size=0.10,edge.arrow.width=1.8,
     edge.width=0.6
)
title(sub="RRG CCBH4851", cex.sub = 0.75, font.sub = 3, col.sub =
"black")


set.seed(2397)

plot(Rede,
     layout=layout_with_fr,
     vertex.label.dist=0,
     vertex.label.color='black',
     vertex.label.font=1,
     vertex.label.cex=1,
```

```
     edge.arrow.size=0.025,edge.arrow.width=0.7,
     edge.width=0.06
)
title(sub="RRG CCBH4851", cex.sub = 0.75, font.sub = 3, col.sub =
"black")


V(Rede)$name <- vert$gene_CCBH4851

nrow(vert)

nrow(arestas)

sum(V(Rede)$color == "yellow")


sum(V(Rede)$color == "lightblue")



scientific(graph.density(Rede, loops=TRUE))

p.kin <- degree_distribution(Rede, mode="in")
p.kin.na <- ifelse(p.kin == 0,NA,p.kin)
min.kin <- min(degree(Rede, mode="in"))
max.kin <-  max(degree(Rede, mode="in"))
plot(min.kin:max.kin, p.kin.na,
     xlab= "k-in (grau input)", ylab= "P(k-in)", type="h")
title(sub="Figura 2: DistribuiÃ§Ã£o de Grau k-in",
      cex.sub = 0.75, font.sub = 3, col.sub = "black")


x.in <- log10(min.kin:max.kin)
y.in <- log10(p.kin.na)
data.in <-
  data.frame(X=x.in,Y=y.in) %>%
  filter(!is.na(X) & !is.na(Y) & X != -Inf)
ajuste.in <- lm(Y~X,data=data.in)
log.A.in <- ajuste.in$coefficients[1]
A.in <- 10^(log.A.in)
gama.in <- -ajuste.in$coefficients[2]
#
plot(x.in, y.in,
     xlab= "log(k-in)", ylab= "log P(k-in)")
plotrix::ablineclip(log.A.in, -gama.in, x1= 0,x2=log10(max.kin))


p.kout <- degree_distribution(Rede, mode="out")
p.kout.na <- ifelse(p.kout == 0,NA,p.kout)
min.kout <- min(degree(Rede, mode="out"))
max.kout <- max(degree(Rede, mode="out"))
#
plot(min.kout:max.kout, p.kout.na,
     xlab= "k-out (grau output)", ylab= "P(k-out)", type="h")
title(sub="Figura 3: DistribuiÃ§Ã£o de Grau k-out",
      cex.sub = 0.75, font.sub = 3, col.sub = "black")

plot(min.kout:max.kout, p.kout.na,
```

```
     xlab= "k-out (grau output)", ylab= "P(k-out)",
type="h",log="y")
title(sub="Figura 4: DistribuiÃ§Ã£o de Grau k-out",
      cex.sub = 0.75, font.sub = 3, col.sub = "black")


x.out <- log10(min.kout:max.kout)
y.out <- log10(p.kout.na)
data.out <-
  data.frame(X=x.out,Y=y.out) %>%
  filter(!is.na(X) & !is.na(Y) & X != -Inf)
ajuste.out <- lm(Y~X,data=data.out)
log.A.out <- ajuste.out$coefficients[1]
A.out <- 10^(log.A.out)
gama.out <- -ajuste.out$coefficients[2]
#
plot(x.out, y.out,
     xlab= "log(k-out)", ylab= "log P(k-out)")
title(sub="Figura 5: DistribuiÃ§Ã£o de Grau k-out log-log",
      cex.sub = 0.75, font.sub = 3, col.sub = "black")
plotrix::ablineclip(log.A.out, -gama.out, x1= 0,x2=log10(max.kout))


CoeffCluster.global <-
scientific(transitivity(Rede,type="globalundirected"))

CoeffCluster.medio <- scientific(transitivity(Rede,type="average"))

CoeffCluster.i <- transitivity(Rede,type="localundirected",
                               isolates = "zero")

hist(CoeffCluster.i,
     xlab= "coeficiente de clusterizaÃ§Ã£o local", ylab=
"frequÃªncia", main=NULL)
title(sub="Figura 6: DistribuiÃ§Ã£o Total de Coef.
ClusterizaÃ§Ã£o",
      cex.sub = 0.75, font.sub = 3, col.sub = "black")

propCzero <-table(CoeffCluster.i)[1]/nrow(vert)


propChum <-
table(CoeffCluster.i)[nrow(table(CoeffCluster.i))]/nrow(vert)



hist(ifelse(CoeffCluster.i ==0 | CoeffCluster.i ==1, NA,
CoeffCluster.i),
     xlab= "coeficiente de clusterizaÃ§Ã£o local", ylab=
"frequÃªncia", main=NULL)
title(sub="Figura 2: DistribuiÃ§Ã£o Parcial de Coef.
ClusterizaÃ§Ã£o",
      cex.sub = 0.75, font.sub = 3, col.sub = "black")


k.i <- degree(Rede,mode="all")
```

```
C.k.i <- CoeffCluster.i
#
plot(k.i, C.k.i,
     xlab="k (grau total)", ylab= "C(k) (coef. cluster. por grau
k)")
title(sub="Figura 7: Coef. ClusterizaÃ§Ã£o em funÃ§Ã£o do grau k
dos vÃ©rtices ",
      cex.sub = 0.75, font.sub = 3, col.sub = "black")


C.k.i.filtrado <- transitivity(Rede,type="localundirected",
                               isolates = "NaN")

plot(k.i, C.k.i.filtrado,
     xlab="k (grau total)", ylab= "C(k) (coef. cluster. por grau
k)")
title(sub="Figura 7: Coef. ClusterizaÃ§Ã£o em funÃ§Ã£o do grau k
dos vÃ©rtices ",
      cex.sub = 0.75, font.sub = 3, col.sub = "black")


count_components(Rede)


components(Rede)$csize

plot(components(Rede)$csize)


reguladores.por.grupo <- c()
for(i in 1:count_components(Rede)){
  grupo <-
names(components(Rede)$membership)[components(Rede)$membership ==
i]
  reguladores.por.grupo<-
    c(reguladores.por.grupo,
      sum(grupo %in% auxTF$gene_CCBH4851))
}


propor.reguladores.por.grupo <-
reguladores.por.grupo/components(Rede)$csize


probab.n.reguladores.por.grupo <-
  table(reguladores.por.grupo)/count_components(Rede)

plot(probab.n.reguladores.por.grupo,
     log= "xy",
     ylim=c(0.01,1))


tab.modoRegula <-
  table(arestas$`mode of regulation`)
tab.modoRegula <- tab.modoRegula[order(tab.modoRegula)]

auto_regul <-
  filter(arestas, `Regulator (TF)` == `Target gene`)
```

```
#
tab.modo.Auto_regul <-
  table(auto_regul$`mode of regulation`)
tab.modo.Auto_regul <-
tab.modo.Auto_regul[order(tab.modo.Auto_regul)]

diameter(Rede,directed = TRUE,unconnected = TRUE)
diameter(Rede,directed = FALSE,unconnected = TRUE)

mean_distance(Rede, directed = TRUE, unconnected = TRUE)
mean_distance(Rede, directed = FALSE, unconnected = TRUE)


triad_census(Rede)

triad_census(Rede)[9]

Rede2 <- graph_from_data_frame(d = filter(arestas,
                                          `mode of regulation` ==
"+" |
                                            `mode of regulation` ==
"-"),
                               directed = TRUE, vertices = vert )

triad_census(Rede2)

triad_census(Rede2)[9]


Rede3 <- graph_from_data_frame(d = filter(arestas,
                                          `mode of regulation` ==
"+" ),
                               directed = TRUE, vertices = vert
)


triad_census(Rede3)

triad_census(Rede3)[9]


Rede4 <- graph_from_data_frame(d = filter(arestas,
                                          `mode of regulation` ==
"-" ),
                               directed = TRUE, vertices = vert
)

triad_census(Rede4)

triad_census(Rede4)[9]


V(Rede)$name <- V(Rede)$rotulo



hubs.em.ordem.dec <-
(hub_score(Rede)$vector)[order((hub_score(Rede)$vector),
```

```
                                                         decreasing =
TRUE)]

authority.em.ordem.dec <-

authority_score(Rede)$vector[order((authority_score(Rede)$vector),
                                          decreasing = TRUE)]


hubs.em.ordem.dec[1:10]

authority.em.ordem.dec[1:10]

k.hubs.em.ordem.dec <- degree(Rede, mode="out")[order(degree(Rede,
mode="out"),
                                                            decreasing =
TRUE)]
k.hubs.em.ordem.dec[1:33]
```
